# Breaking the mold: The first report on germ-free adult marine medaka (*Oryzias melastigma*) models

**DOI:** 10.1101/2023.04.10.536225

**Authors:** Pan-Pan Jia, Yi-Fan Yang, Wei-Guo Li, Jin-Jing Duan, Yan Wang, De-Sheng Pei

**Author notes:** Corresponding author. and (D.S.P).

## Abstract

Marine medaka (*Oryzias melastigma*) animal models play critical roles in environmental and human health by facilitating evaluation of pollutant toxicity and building of disease models. The fish gut microbiota contributes to host health and physiological metabolism, especially special bacterial strains and their functions in marine organisms. However, the distribution of the gut microbiota during medaka growth and development is still unclear, and successful generation of a germ-free (GF) marine medaka model has not been reported to date. In this study, we investigated the microbial composition with the major phyla and genera of marine fish at different life stages, as well as the isolated culturable intestinal bacteria, and then identified them by sequencing of the16S rRNA V3-V4 region. Importantly, the early stage model (larvae) of GF marine medaka without feeding and long-term (from juvenile to early adult stages) GF fish fed GF brine shrimp (*Artemia* sp.) were first generated. Moreover, the basic indexes and behavioral ability of GF fish showed weaker and delayed developmental changes compared to conventionally raised (CR) marine medaka at the same life stages. Notably, the significant differences in the histopathological characteristics of immune organs, intestinal tissues and the reproductive system were observed between GF and CR early-adult and adult fish. Furthermore, the transcriptomic profiles of the screened critical genes in signaling pathways in GF and CR marine medaka were also explored to illustrate the developmental impacts of the absence of the intestinal microbiota during the host growth. Comprehensively, our study provided novel insights into the intestinal microbiota distribution of CR fish during growth, and GF marine medaka from the larval to adult stages *via* GF fish food preparation. The histopathological and transcriptomic differences indicated the potential microbial regulation on growth, and application prospects of GF medaka fish models to clarify the relationships of intestinal bacterial functions to host health in the future.

**Significance:** The generation and application of germ-free (GF) fish models are mostly limited to the early life stages with innate immunity and without feeding. Marine medaka (*Oryzias melastigma*) is a critical animal for evaluating environmental toxicity and human disease models. The gut microbiota contributes to host growth and development, but GF model of this organism has not been successfully generated. In this study, we revealed for the first time the distribution of the gut microbiota in marine medaka during growth and generated GF fish from the larval to adult stages with GF *Artemia* provided daily as food. According to the basic indexes, weaker behavioral ability, smaller immune organs, reproductive system, intestinal tissues, and transcriptome, the delayed development and differences indicated the negative influences of the absence of the microbiota in GF medaka, compared to conventionally raised (CR) fish at the same life stages. All these results provided novel insights into the application of GF medaka models to define intestinal bacterial functions in the host.

**Graphical abstract:** 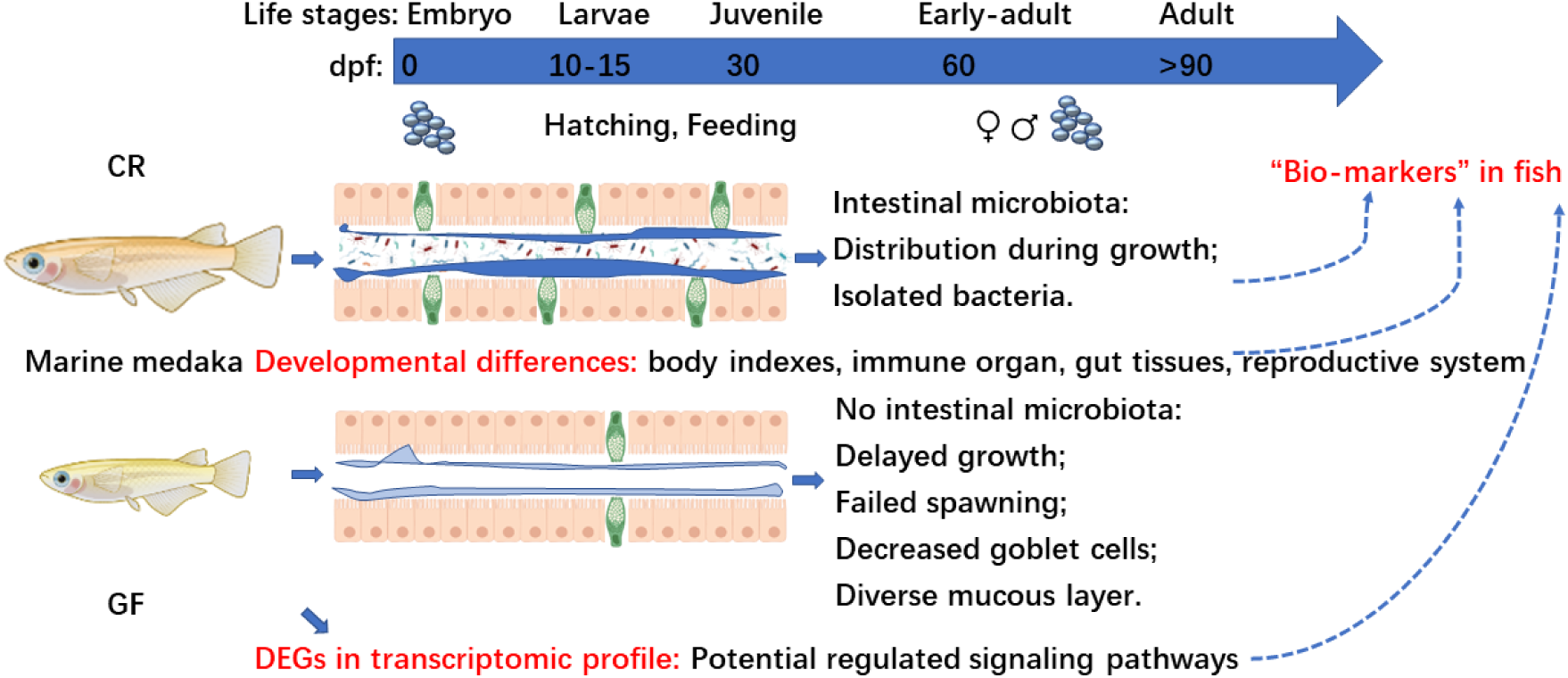

This work revealed the distribution of the gut microbiota in marine medaka during growth, and successfully generated GF marine medaka models from larvae to adults with GF *Artemia* as food, which indicated the delayed development in the absence of the microbiota in GF fish. Moreover, the histopathological analysis presented further evidence of developmental differences in immune organs, intestinal villi, goblet cells, gonad tissues and cell maturation between GF and CR fish at various life stages. Finally, the transcriptomic profile showed the significantly differentially regulated genes, which combined with the major bacteria can be potential “biomarkers” to explore the inner mechanisms or signaling pathways of GF fish models for studying host development and health.

## Introduction

Marine medaka (*Oryzias melastigma*) is a specie native to the coastal waters of Pakistan, China, Korea, Japan, India, Burma and Thailand, and is a useful and sensitive fish model for monitoring ocean and estuary environments contaminated with heavy metals, endocrine disruptors and organic pollutants (1–3). Marine medaka (*O. melastigma*) and Japanese medaka (*Oryzias latipes*) belong to the order Beloniformes, family Adrianichthyidae, and genus Oryzias and have been extensively used worldwide in ecotoxicological studies to evaluate various environmental pollutants and their potential mechanisms of toxicity (4, 5). As reported in previous studies, omics analyses and genome sequencing of marine medaka were recently conducted, providing new insights for further research (6, 7). Germ-free (GF) freshwater zebrafish (*Danio rerio*) were previously reported by Rawls *et al*. to evaluate host-microbe interactions using gene expression analysis in gnotobiotic aquatic animals (8). Aquatic animals in gnotobiotic systems (that is, the rearing of animals in axenic conditions or with a known microflora) is still rare but is required for understanding the mechanisms involved in host-microbe interactions and to evaluate new treatments for disease control (9). Analysis of the effects of age showed that sexually immature (juvenile) fish had a more abundant of gut microbiota than elderly adult fish. However, the certain questions regarding the gut microbiota profile of marine medaka during growth and development, the bacteria that can be isolated and the especial functions, and the generation of GF marine fish models from larvae to adults, as well as GF food for the fish, remain unanswered. Thus, research on the microbial profile of marine medaka, developmental differences, and the development of GF food are still urgently needed and will be significant for understanding intestinal bacterial functions and host and environmental health.

*O. melastigma* models are usually applied in marine ecotoxicology studies of neurotoxicity, embryo toxicity, cardiac toxicity, immunotoxicity, endocrine disruption, and metabolism alterations after exposure to contaminants and other environmental stressors. The marine medaka model has advantages of (1) easy breeding in the laboratory of small adult fish (usually 2-4 cm in size), exhibiting distinct sexual dimorphism with anal fin morphology, and a short generation time (2-3 months), (2) high spawning ability and transparent embryos 10-30 eggs/day with the fertilization rate of over 80%, (3) adaptation capacity for a wide range of salinity, temperature, and water quality conditions, and (4) an identified genome, etc (1, 4, 6, 7). GF fish, including zebrafish and marine medaka models. Thus, it is possible to study the interactions between microbes and hosts by implanting specific microbes at key points, during which microbial genotypes and community competition can be analyzed, and how microbial colonization or metabolic products affect the host’s biological processes, including gene expression, development, physiology, immunity, diet and longevity, and physiological behaviors can be explored (10). The microbial community plays key roles in vertebrate host development, including by affecting the immune system, providing nutrients, and stimulating cell renewal in the intestinal epithelium (11, 12). In zebrafish, previous studies have proven the importance of understanding culturable gut microbiota diversity, how intestinal bacterial communities assemble in tandem with host development (from early to adult stages), and how spatial and temporal species-richness relationships in bacterial community assembly and dynamics during life are shaped (13, 14). However, the community profile as well as the isolation and identification of the gut microbiota of marine medaka has not been sufficiently studied, especially the changes in microbial diversity and richness that occur during the different developmental stages. In addition, GF marine medaka from larval to adult life stages will provide important novel insights into this animal model and its application.

Medaka fish can be distinguished into different life stages, including the embryo, larva, juvenile (sexually immature), early adult (begin to spawn), and sexually mature adult in terms of the growth and development of host organs and physiological systems, which are similar to the stages observed for zebrafish, with the same generation time of 2-3 months (15–17). Marine medaka embryos hatch to larvae at approximately 12-16 dpf, and then the later larvae turn to juveniles at 25-30 dpf, and from the young fish grow to the early adult with the possible spawn ability at 60 dpf, and finally reach to adult at 70 dpf to 3 months (18–20). GF fish are rescued from the newly-born embryos to larvae to investigate the degenerative changes that occur in the late life stages in absence of microbes. However, most studies on standardized GF fish models are usually performed on early stage of larvae, which lack a mature immune system, such as in GF zebrafish larvae (< 7 dpf) without feeding when the yolk is not completely absorbed (21–25). Currently, several constraints still hamper the widespread generation and application of GF and gnotobiotic fish from early life stages to juvenile and adult models for research purposes involving the diversity of disinfection/sterilization methods used to produce and maintain axenic organisms and the difficulty in assuring a complete axenic state in culture systems (9). Among the specific foods special for fish from larval opening period to adulthood, the *Artemia* including nauplii are the most widely used live feed for marine fish and shellfish (26). *Artemia* exhibit a particular form of reproduction (oviparous reproduction) under adverse environmental conditions, wherein developing embryos are enclosed in multilayer shells (also called cysts) to against exterior aggressions, make the GF models possible by treatment of strong oxidizing agents (e.g. sodium hypochlorite) at high concentrations efficiently (9, 27, 28). Previously, GF *Artemia* models were used in environmental toxicity and bacterial function studies, such as for monitoring the probiotic effect of *Aeromonas hydrophila* on gut development in GF *Artemia* nauplii, and for studying the gnotobiotic feed chain for GF fish, such as larval sea bass (*Dicentrarchus labrax* L.), according to a previously described generation method (29, 30). The development of gnotobiotic models including GF *Artemia* cysts and nauplii was described by Sorgeloos *et al* in 1986 and later by Marques *et al* in 2004, but the methods were complicated (27, 31). However, the generation approaches of GF *Artemia* models as GF fish food, which requiring newly and freshly hatched and feed daily, are still limited to wide application due to the challenges posed by the complex treatment steps and long incubation times. Thus, novel axenic experiments including quick sterilization and simple and shorter incubation for GF *Artemia* were explored in this study, which providing much support for the successful obtain of chronic GF fish models.

The intestinal microbiota of animals, including fish models, is considered as a critical indicator of environmental and organism health (10). However, the composition and functional mechanisms of the gut microbiota in important fish models during growth and development still need to be clearly elucidated. Therefore, in this study, the developing microbial community and culturable and isolated strains of marine medaka were identified. Furthermore, the use of GF *Artemia* as the food for GF marine medaka models was explored, facilitating the successful generation of GF marine medaka from larval to juvenile and adult stages *via* optimization of the culture conditions with the advantages, for example, the sea-salt medium inhibited the bacteria survival and prolonged the GF *Artemia* live time compared with the freshwater. Importantly, the growth and developmental indexes and immune organs with histopathological differences between GF and conventionally raised (CR) models were also analyzed in depth during various fish life stages to illustrate the key functions of the intestinal microbiota in host health. Finally, all of the investigations in this study will provide comprehensive knowledge or a deeper understanding of marine medaka model and will potentially serve as an important reference for the development of GF animal models and their application in biomedicine and other life science fields.

## Results

### Gut microbiota of female and male marine medaka during development

In this study, we collected marine medaka of both sexes at different life stages (F-1, F-3, F-5, M-1, M-3, M-5) and measured the body length (BL), body weight (BW), intestinal weight, and calculated the body mass index [BMI, body weight (mg)/body length (cm)^2^] and intestinal somatic index [ISI, intestinal weight (mg)*100%/body weight (mg)] (**Fig. 1**). During growth, the BL and BW of female and male fish at 3 and 5 months-old showed significant increases (*p*<0.01 and *p*<0.001) compared to those of 1 month-old juveniles, respectively. Moreover, the BMI and ISI of marine medaka exhibited no significant differences among various life stages (**Fig. 1A**). As the basic indexes during development, the BL, BW, BMI and ISI were selected as the key environmental factors for evaluating the relationship of the intestinal microbial changes among various groups divided by life stage.

**Figure 1.**
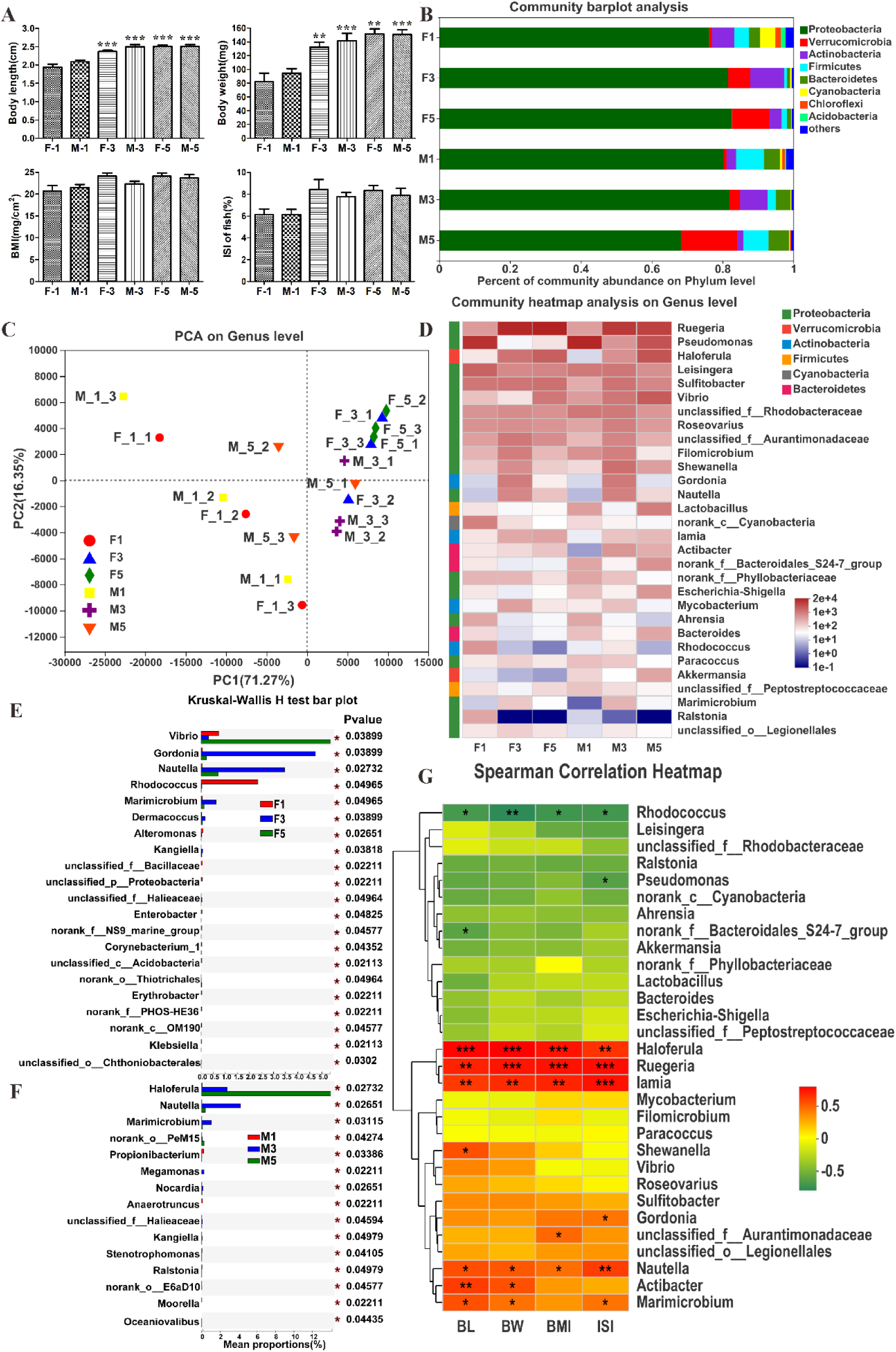
Distribution and significantly changed genera of the gut microbiota in female and male marine medaka during growth and development. (A) The body length (BL), body weight (BW), intestinal weight, body mass index (BMI), and intestinal somatic index (ISI) of marine medaka at the different life stages in both gender (F-1, F-3, F-5, M-1, M-3, M-5). (B) The dominant phyla of the gut microbiota in female and male marine medaka from juvenile to adults with the average percent of community abundance. (C) The PCA based on the genus level showed the similarity and distinction of juvenile fish with adults. (D) The heatmap presented the top 30 major genera of the gut microbiota with the belonged phylum in juvenile to adult marine medaka of both sexes. The multiple-group comparison with the Kruskal-Wallis H test proved that there were the 21 significantly changed genera in female (E) and 15 significantly changed genera in male (F) marine medaka at different life stages. (G) The Spearman correlation heatmap presenting the top 30 species showed the correlation of major genera with the developmental indexes of fish during growth. The symbols *, **, and *** indicated *p*<0.05, 0.01, and 0.001, respectively, with significant differences between different groups or between indexes and species.

To explore the microbial composition and functions, the gut microbiota in marine medaka from juvenile to adult stages was analyzed. In this study, we obtained high-quality sequences with an average length of 433 bp belonging to 1 Domain, 1 Kingdom, 43 Phylum, 93 Class, 200 Order, 381 Family, 795 Genus, 1318 Species, and 2181 OTUs from a total of 18 samples. Among the alpha diversity indexes, the sobs, Shannon, ACE and Chao indexes were not changed among groups, but Simpson index showed a significant difference between F-5 and M-5 with a *p* value of 0.019 (shown in **Table S2;** the common and specific OTUs were shown in **Fig. S1**). In female and male marine medaka, the dominant phyla of gut microbiota from the juvenile to adult stages were Proteobacteria (68.23-81.58%), Verrucomicrobia (0.73-15.77%), Actinobacteria (1.86-9.57%), Firmicutes (0.82-7.82%), Bacteroidetes (0.89-5.79%), Cyanobacteria (0.15-4.30%), Chloroflexi (0.06-1.56%), and Acidobacteria (0.003-1.287%), with the average percent of community abundance (**Fig. 1B**). Among them, the phylum Verrucomicrobia was enriched in both female and male fish during growth and development. In contrast, the phyla Firmicutes, Cyanobacteria, Chloroflexi, and Acidobacteria were depleted in both female and male fish during growth. In the Beta diversity analysis, the PCA analysis based on the genus level showed the similarity of samples from female and male marine medaka and the distinction of fish at juvenile stage with adult fish at 3 and 5 months post fertilization (mpf), which indicating changes in the intestinal community in both gender during development (**Fig. 1C**). Moreover, the phylogenetic tree of the gut microbiota of marine medaka at 1, 3 and 5 mpf showed the distances and abundances of different bacteria in the six groups at the class level, which belonged to Alphaproteobacteria, Gammaproteobacteria, Verrucomicrobia, and Actinobacteria (**Fig. S2**). Furthermore, the heatmap with the top 30 genera of the gut microbiota showed the major genera and the associated phyla based on relative abundance in marine medaka from the juvenile to adult stages (**Fig. 1D**). Among the major genera, *Ruegeria*, *Haloferula*, *Sulfitobacter*, *Vibrio*, *unclassified_f_Aurantimonadaceae*, *Nautella*, *Iamia*, *Actibacter*, *Marimicrobium*, and *unclassified_o_Legionellales* belonging to the phyla of Proteobacteria, Verrucomicrobia, Actinobacteria and Bacteroidetes, were increased with relative abundances of 2.10-46.04-54.28% and 1.85-29.31-23.53%, 0.27-5.99-10.37% and 0.05-2.75-13.99%, 5.62-5.73-6.54% and 1.29-5.99-3.47%, 0.72-0.30-5.35% and 1.96-5.51-11.50%, 1.81-5.28-2.98% and 0.90-3.80-1.17%, 0.04-3.46-0.70% and 0.03-4.22-0.40%, 0.28-1.41-1.64% and 0.09-0.69-0.91%, 0.27-0.38-0.67% and 0.01-2.40-1.25%, 0.04-0.62-0.13% and 0.0-1.06-0.11%, 0.26-0.52-0.29% and 0.07-0.24-0.15%, correspondingly in female and male fish from 1 to 3 and 5 mpf. In contrast, the abundances for major genera such as *Pseudomonas*, *Leisingera*, *norank_c_Cyanobacteria*, *norank_f_Phyllobacteriaceae*, *Ahrensia*, and *Ralstonia*, belonging to phyla of Proteobacteria and Cyanobacteria, were decreased with ranges of 35.30-0.11-0.30% and 47.40-2.34-14.60%, 9.26-3.31-3.72% and 5.08-5.49-2.86%, 4.24-0.34-0.13% and 0.39-0.19-0.11%, 0.99-0.98-0.32% and 1.56-0.70-0.12%, 1.46-0.08-0.11% and 1.46-0.11-0.04%, 1.53-0.00-0.00% and 0.05-0.00-0.00% in female and male fish, respectively, during growth (**Fig. 1D**). Based on the community analysis, we hypothesized that the dominant phyla and major genera of marine medaka changed during development from the juvenile (1 month) to adult stage and finally tended to be stable from the early adult (3 mpf) to later life stage (5 months).

The significantly changed genera in the gut microbiota of female and male marine medaka at different life stages and their correlation with developmental indexes were further analyzed and presented with the top 30 species and their p values (**Fig. 1**). In females, the multiple-group comparison with the Kruskal-Wallis H test proved that the abundances of 21 genera, namely, *Vibrio*, *Gordonia*, *Nautella*, *Rhodococcus*, *Marimicrobium*, *Dermacoccus*, *Alteromonas*, *Kangiella*, *unclassified_f_Bacillaceae, unclassified_p_Proteobacteria*, *unclassified_f_Halieaceae*, *Enterobacter*, *norank_f_NS9_marine_group*, *Corynebacterium_1*, *nclassified_c_Acidobacteria*, *norank_o_Thiotrichales*, *Erythrobacter*, *norank_f_PHOS-HE36*, *norank_c_OM190*, *Klebsiella*, and *unclassified_o_Chthoniobacterales*, were significantly changed (*p*<0.05) in marine medaka at different life stages (**Fig. 1E**). In males, the abundances of 15 genera, namely, *Haloferula*, *Nautella*, *Marimicrobium*, *norank_o_PeM15*, *Propionibacterium*, *Megamonas*, *Nocardia*, *Anaerotruncus*, *unclassified_f_Halieaceae*, *Kangiella*, *Stenotrophomonas*, *Ralstonia*, *norank_o_E6aD10*, *Moorella*, and *Oceaniovalibus*, were significantly different (*p*<0.05) among various life stage groups. (**Fig. 1F**). Furthermore, the dominant genera with the top 30 species were analyzed by a two-group comparison by Students’ test, and the differences in a total of 9 groups divided by life stage and sex were compared (the mean abundance and significant changes with p values were shown in **Table S3**). In details, in group 1 (F1 vs. F3), the bacterial genera *Ruegeria*, *Haloferula*, *Nautella*, *Iamia*, *Paracoccus*, *Marimicrobium*, *Dermacoccus*, *Kangiella*, *Aureimonas*, and *Idiomarina* were significantly increased with *p* values <0.05 and 0.001, while the bacterial genera *unclassified_p_Proteobacteria*, *Rothia*, *Lawsonella*, *norank_p_WS6*, *[Eubacterium]_hallii_group*, and *Klebsiella* were significantly decreased with *p* values <0.05 and 0.01. In group 2 (F1 vs. F5), the bacterial genera *Ruegeria*, *Iamia*, *norank_o_PeM15*, *norank_o_Thiotrichales*, and *Gelria* were significantly enriched, with *p* values <0.05, 0.01, and 0.001, while, the bacterial genera *unclassified_p_Proteobacteria*, *Rothia*, *Lawsonella*, *Klebsiella*, and *Parasutterella* were significantly depleted, with *p* values <0.05 and 0.01. However, in group 3 (F1 vs. M1), only *Lawsonella* was significantly increased with a *p* value <0.05, and *Moorella* was significantly decreased with a *p* value <0.05. In group 4 (F3 vs. F5), the bacterial genera *Vibrio*, *norank_o_Thiotrichales*, and *Vogesella* were significantly increased with *p* values <0.05 and 0.01, while *Filomicrobium*, *Nautella*, *Marimicrobium*, *norank_f_OCS116_clade*, *Dermacoccus*, *Dietzia*, *Marixanthomonas*, *norank_f_Chlamydiaceae*, *Marinococcus*, and *Rheinheimera* were significantly decreased with *p* values <0.05 and 0.01. In group 5 (F3 vs. M3), the bacterial genera *Roseovarius* and *Fusicatenibacte* were significantly increased, with *p* values <0.05, while *Dermacoccus*, *Dietzia*, *norank_o_JG30-KF-CM45*, *Exiguobacterium*, *Marinococcus*, *Janibacter*, *Sphingobium*, and *Idiomarina* were significantly decreased with *p* values <0.05 and 0.01. In group 6 (F5 vs. M5), four bacteria of *Ruegeria*, *Filomicrobium*, *norank_o_Thiotrichales*, and *Vogesella* were significantly decreased with *p* values <0.05 and 0.01. In group 7 (M1 vs. M3), the bacterial genera *Ruegeria*, *Sulfitobacter*, *unclassified_f_Aurantimonadaceae*, *Nautella*, *Haloferula*, *Marimicrobium*, *Iamia*, *unclassified_o_Legionellales*, *Dietzia*, and *norank_o_Thiotrichales* were significantly increased with *p* values <0.05 and 0.01, while *Moorella* was significantly decreased with *p* value <0.05. In group 8 (M1 vs. M5), the bacterial generra *Ruegeria*, *Haloferula*, *Actibacter*, *Iamia*, *norank_f_Erysipelotrichaceae*, *unclassified_o_Cellvibrionales*, *Aeromicrobium*, *Exiguobacterium*, and *Oceaniovalibus* were significantly enriched with *p* values <0.05, 0.01, and 0.001, while *Rhodococcus* and *Moorella* were significantly depleeted with *p* values <0.05 and 0.01. In group 9 (M3 vs. M5), the genera *Haloferula*, *norank_f_Erysipelotrichaceae*, *Stenotrophomonas*, and *Oceaniovalibus* were significantly increased with *p* values <0.05, and *Roseovarius*, *unclassified_f_Aurantimonadaceae*, *Nautella*, *Marimicrobium*, *Paracoccus*, *norank_f_OCS116_clade*, *Pseudonocardia*, *norank_f_Chlamydiaceae*, *Ilumatobacter*, *unclassified_f_Intrasporangiaceae*, and *Aureispira* were significantly decreased with *p* values <0.05 and 0.01. Interestingly, the significantly changed genera with a mean abundance of 0% potentially indicated bacterial appearance or disappearance during the life stages (**Table S3**). For example, *Moorella* disappeared during development in M3 and M5 a mean value of 0% compared to that in M1 fish (0.012%). Notably, Spearman correlation analysis of all the samples at different life stages was performed to explore the relationship of major genera with the developmental indexes of fish during growth, and the heatmap represented the top 30 species based on abundance (**Fig. 1G**). The major genera *Haloferula*, *Ruegeria*, *Iamia*, *Nautella*, *Actibacter*, and *Marimicrobium* were observed to be positively and significantly related to the BL, BW, BMI and ISI developmental indexes during fish life, and *Rhodococcus* was negatively related to the indexes at all life stages.

To explore the predicted functions of the gut microbiota in marine medaka, 16S sequencing data was analyzed by the Bugbase (https://BugBase.cs.umn.edu/index.html) tool, and then microbial phenotypes involving the major 8 types including Gram_Negative, Gram_Positive, Biofilm Forming, Pathogenic, Mobile Element Containing, Oxygen Utilizing including Aerobic, Anaerobic, Facultatively_Anaerobic, and Oxidative Stress Tolerant, were predicted (**Fig. S3**). In details, the dominant genera *Haloferula*, *Sulfitobacter*, *Pseudomonas*, *Leisingera*, and *Filomicrobium* contributed to the Gram_Negative phenotypes, and the genera *Gordonia*, *Iamia*, and *Mycobacterium* contributed to the Gram_Positive phenotype, of the intestinal microbiota in female and male fish during growth (**Fig. S3A** and **B**). Moreover, the dominant genera *Haloferula*, *Pseudomonas*, *Leisingera*, *Gordonia*, and *Iamia* contributed to the Aerobic phenotype, and the genera *Sulfitobacter*, *Roseovarius*, *Vibrio*, *unclassified_f_Rhodobacteraceae*, and *Nautella* mainly contributed to the Facultatively-Anaerobic phenotype of the intestinal microbiota in female and male fish from 1 to 3 and 5 mpf (**Fig. S3C** and **D**). Importantly, the genera *norank_f_Bacteroidales-S24-7_group*, *unclassified_f_Peptostreptococcaceae*, *Blautia*, *Gelria*, and *Bacteroides* mainly contributed to the Anaerobic phenotype, and the genera *Pseudomonas*, *Leisingera*, *Vibrio*, *norank_f_Phyllobacteriaceae*, and *Ahrensia* criticaly contributed to the Potentially-Pathogenic phenotype of intestinal microbiota in female and male fish at various life stages (**Fig. S3E** and **F**). Finally, the dominant genera *Sulfitobacter*, *Leisingera*, *Filomicrobium*, *Roseovarius*, and *unclassified_f_Rhodobacteraceae* contributed to the Forms_Biofilms phenotype, and the genera *Pseudomonas*, *Vibrio*, *norank_f_Phyllobacteriaceae*, *Ahrensia*, and *Escherichia-Shigella* mainly contributed to the Contains_Mobile_Elements of the intestinal microbiota of female and male fish from the juvenile to adult stages (**Fig. S3G** and **H**). Interestingly, the bacterial functions during development can be predicted, for example, the genus *Vibrio* was enriched in the adult period, possibly leading to higher pathogenicity and mobile element levels in the fish. In the F1 and M1 fish, the abundance of human disease and organismal systems pathway was higher than that in the fish at 3 and 5 mpf (**Fig. S4**). Furthermore, according to the KEGG analysis of abundance of level 2 pathways, the amino acid metabolism, metabolism of other amino acids, membrane transport, cell growth and death, and digestive system pathways were significant different between the females and males at various life stages (F1 vs. F3, F1 vs. F5, M1 vs. M3, M1 vs. M5) with *p* values <0.05, 0.01, and 0.001, respectively (**Table S4**). Interestingly, infectious disease (bacterial and viral), the cancer (specific types), and the cardiovascular disease were significantly increased with life stage in female and male fish (**Table S4**).

### Isolated and identified bacterial strains from the intestines of adult marine medaka

In this study, the isolation approach and the subcultures of individual bacterial strains from the intestinal microbiota in marine medaka were optimized (**Fig. S5A**). Based on the morphological distinction, the bacterial colonies were selected and purified before identification (**Fig. S5B**). After 16S rRNA gene-based sequencing (data not shown), NCBI-BLAST searching using the bacterial database and comparison with similar bacterial strains were performed. We summarized and listed he best-matched bacterial strain (name, version, source, submission and so on), identity, sequence length, and sample information (**Table S5** and several genera shown in **Fig. S5C**). In our work, a total of 22 bacterial colonies belonging to 10 genera with the identity >98% were indicated. In details, the colony of the No.1 sample belonged to the *Pseudomonas* genus, with the best-matched bacteria being *Pseudomonas khazarica sp*. nov., a polycyclic aromatic hydrocarbon-degrading bacterium isolated from Khazar Sea sediments, with 99.79% identity and 1416 bp length; BLAST analysis of the No. 8, 9, 13, and 20 samples showed that they were the same or similar with 100 or 99.93 % identity (**Table S5**). The colony of the No.2 sample belonged to the *Shewanella* genus, with the best-matched species being *Shewanella seohaensis sp*. nov., isolated from tidal flat sediment, with 99.86% identity and a length of 1421 bp; the No.16 sample was indicated to be the same, with 100% identity (**Table S5**). The colony of the No.3 sample possibly belonged to the genus *Vibrio*, with the best-matched bacteria of *Vibrio owensii sp.* nov., isolated from cultured crustaceans in Australia, with 99.65% identity and a length of 1428 bp. While, the colony of the No.4 sample also belonged to the *Vibrio* genus, but the best-matched bacteria was *Vibrio alfacsensis sp*. nov., isolated from marine organisms, with 99.93% identity and 1435 bp length (**Table S5**). However, the identity of No.3 and No.4 was 96.38%, which indicated that they were different bacterial species with low similarity. The colony of the No.5 sample also belonged to *Vibrio* genus with the best-matched bacteria *Vibrio alginolyticus* strain, with 99.72% identity and 1429 bp length, and the No.17 sample was similar, with 99.79% identity (**Table S5**). The colony of the No.6 sample seemed to belong to *Vibrio* genus with the best-matched bacteria *Vibrio caribbeanicus sp*. nov., isolated from the marine sponge Scleritoderma cyanea, with 98.60% identity and 1431 bp length. As for the No.7, it was blasted to be best matched with *Staphylococcus succinus sp*. nov., isolated from Dominican amber, with 99.93% identity and 1429 bp length; the No.10 and No.11 samples were the same, with 100% similarity (**Table S5**). The colony of the No.12 sample belonged to the genus *Dietzia*, with the best-matched bacteria *Dietzia aurantiaca sp.* nov., isolated from a human clinical specimen, with 99.06% identity and a length of1375 bp; the same samples No.14, No.15, No.21, and No.11 were also highly similar to No.12 with 100% similarity (**Table S5**). Interestingly, the No.18 sample was highly matched with *Vibrio fluvialis* strain NBRC 103150, with 100% identity and 1433 bp length, but it also appeared to be very similar to *Allomonas enterica* strain LMG 21754, with 99.65% identity and a length of 1433 bp, which indicated that it belonged to the genus *Allomonas*. The comparison of the No.18 sample to the previous *Vibrio* samples of No.3, No.4, No.5, No.6, and No.17 showed few common sequences, indicating that they were different species. Finally, the colony of the No.19 sample was best matched to *Bacillus cereus* ATCC 14579, with 99.79% identity and 1444 bp length by NCBI-BLAST (**Table S5**). Among the genera, the major strains belonged to *Pseudomonas*, *Shewanella*, and *Vibrio* in dominant phylum of Proteobacteria in marine medaka.

### Generation and detection of GF *Artemia* models for fish feeding

In the present study, we modified the model generation process to be highly effective and simple for feeding fish daily, and the sample detection method were optimized to ensure the sterile conditions (**Fig. 2**). In detail, as shown in the flow chart, the protocol mainly included the treatment of *Artemia* cysts, sterile incubation in bottles, washing of the newly hatched GF *Artemia* nauplii, preparation of solution two times to obtain clean and active GF *Artemia*, and feeding after detection (**Fig. 2A**). After treatment, the normal CR cysts and GF cysts showed no significant difference in shape, but a slight color change and a much cleaner and smoother GF cysts surface were observed under a microscope (**Fig. 2B**). Importantly, GF *Artemia* showed a high hatching rate and survival rate after 24 h of incubation, with fewer abnormal and static nauplii present at the bottom. The newly hatched GF *Artemia* as well as the empty shells, incompletely hatched cysts, and eggs that failed to hatch were observed under higher magnification, and the developmental indexes were measured for each replicate group (**Fig. 2C**). Sterile detection of samples, including normal and treated cysts, incubated *Artemia* nauplii, and hatching brine, was performed under aerobic and anaerobic conditions to ensure that GF models without bacteria were obtained (**Fig. 2D**). The detection under aerobic and anaerobic conditions of samples collected from the GF *Artemia* models showed that the normal cysts contained bacteria accompanied by colonies and liquid turbidity, while the treated cysts and sterile incubated nauplii were germ free. Furthermore, the results for samples detection using plates and tubes were recorded from 24 to 168 h (details summarized in **Table S6**).

**Figure 2.**
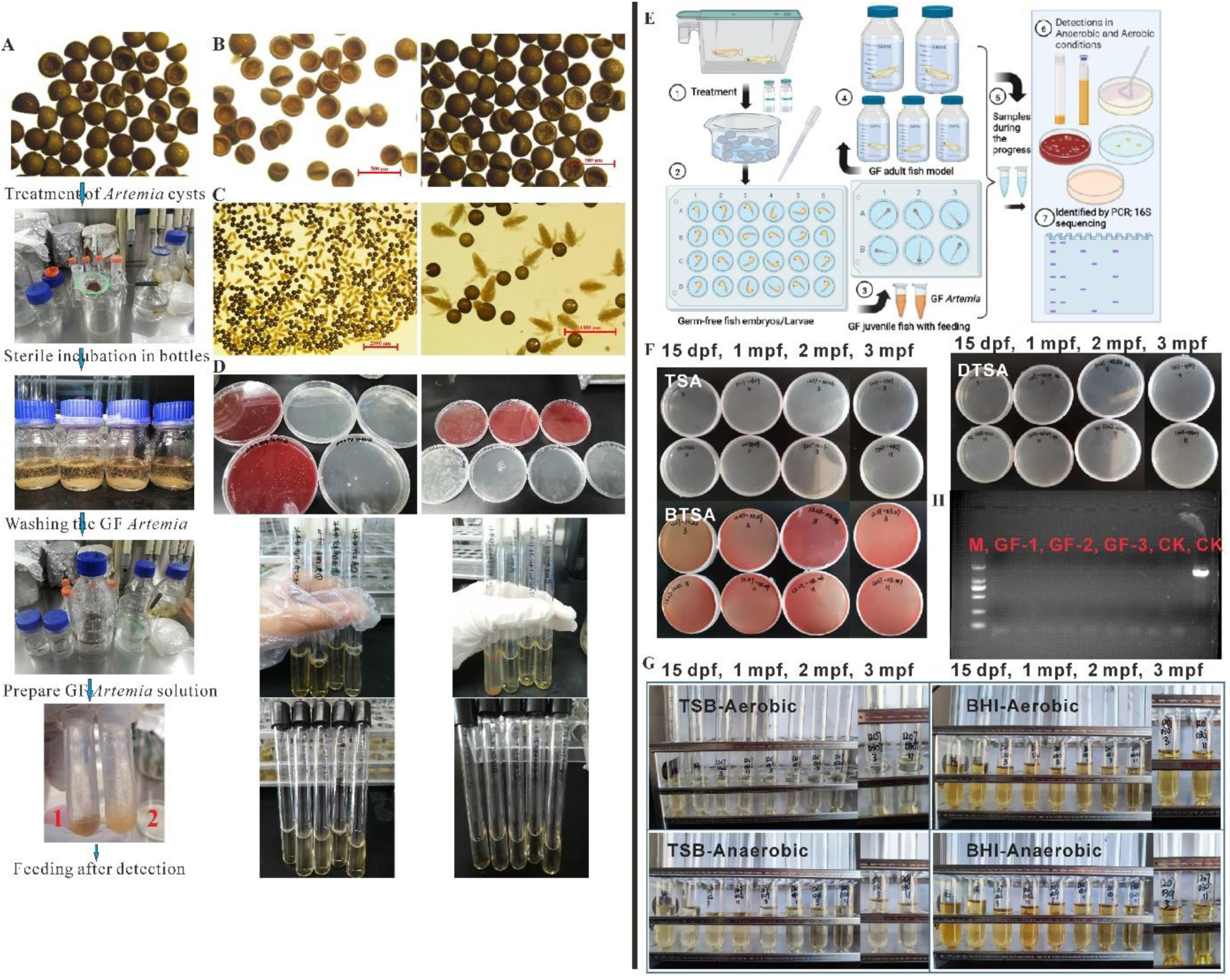
Generation and identification of GF *Artemia* food and GF marine medaka models from larval to juvenile and adult stages. (A) The flow chart of the protocol mainly included the novel treatment and incubation of *Artemia* cysts, sterile incubation in bottles, washing of the newly hatched GF *Artemia* nauplii, preparation of solution for two times to obtain clean and active GF *Artemia*, and feeding after detection. (B) Normal cysts and treated cysts at 50× magnification and 500 μm scale bar. (C) It represented the newly hatched GF *Artemia* nauplii with 10× and 50× magnification (2000 μm and 1000 μm scale bar, respectively). (D) Detection under aerobic and anaerobic conditions of samples collected from the GF *Artemia* models, which showed that the normal cysts contained bacteria, and the treated cysts and sterile incubated nauplii were germ free. (E) Flow chart of the generation and identification of GF marine medaka during different life stages. (F) The pictures showed the detection results for the collected samples from 1 to 3 mpf GF and CR models on TSA plates, blood TSA (BTSA) plates and double layer TSA (DTSA) plates after 7 days of incubation and observation. (G) The pictures showed the detection results for the collected samples from 1 to 3 mpf GF and CR models in TSB and BHI medium with both aerobic and anaerobic conditions after 7 days of incubation and observation. (H) PCR of the collected samples from 1 to 3 mpf GF and CR models by using the 27F and 1492R bacterial primers.

Moreover, the developmental indexes of GF *Artemia* were observed and measured after incubation, such as the average BL (shown in **Table S7**). First, the average hatching rate (counted as hatched *Artemia*, eggs that failed to hatch or shells, and incomplete hatched cysts) of GF *Artemia* at 24 h was as high as 88.15% among 10 replicate groups, with N=30-50 *Artemia* nauplii and cysts per group. The survival rate (recorded as swimming *Artemia* and dead or unmoved nauplii) had an average value of 77.38%, and the average malformation rate was 1.87% among 10 replicate groups, with N=30-50 *Artemia* nauplii per group. Moreover, the average BL was 546.70 µm among 10 replicate groups, with N=10 *Artemia* nauplii per group, and BW was 3.80 mg/100 *Artemia* nauplii per group. Those findings support that the GF *Artemia* models were suitable for the feeding requirements of fish models from larvae.

### Generation and identification of GF marine medaka from larval to adult stages

In this study, we first prepared and compared sterile microparticle food and shelled shrimp eggs as feed for fish larvae (**Fig. S6**), and identified them by multiple detection methods (shown in **Fig. S7** and **Table S8**) before feeding. Then, according to the optimized protocol, GF marine medaka models were generated under sterile conditions (**Fig. 2E**). Moreover, GF marine medaka fed GF *Artemia* after hatching from the larval to juvenile, early-adult and adult stages were maintained and identified by multiple methods (shown in **Fig. 2F-G-H**, and samples listed in **Table S9**). After detection, the GF fish models were collected to examine the developmental indexes and for histopathological and transcriptomic analyses.

### Developmental indexes of GF and CR marine medaka larvae

In this study, the basic developmental indexes of GF and CR fish were recorded, and the images of marine medaka at different life stages were taken by a microscope with a white light background (**Fig. 3**). Meanwhile, the BL and fin length of marine medaka at 1, 2 and 3 mpf stages were measured as described by Jae *et al*. (32). After treatment, the survival rate and germ-free rate of the fish models were 100% at the beginning, but the survival rate decreased to 80.93% at 7 dpf, and to 59.72% at 14 dpf, and to 34.72% at 21 dpf, and to 28.82% at 1 mpf, and to 24.67% at 2 mpf, and finally to 16.67% at 3 mpf during the culture process (n=96 embryos/larvae, three replicates). While the germ-free rate decreased to 91.87%, 91.28%, and 92.03% at 7, 14 and 21 dpf, respectively, during feeding and increased to be stable at 100% from early adulthood to 3 mpf (**Fig. 3A**). The average hatching rate of GF fish at early life stages 12 dpf was 59.72% lower than that of the CR group (66.32%), and the number of heart beats/10s of marine medaka at 1 mpf was significantly lower (13.95) in the GF group than in the CR group (15.20) (**Fig. 3A**). Moreover, the BL of GF fish was significantly (*p*<0.001) lower (7.97, 12.13, and 15.79 mm at 1, 2, and 3 mpf, respectively) than that in the CR group (11.03, 17.61, and 29.09 mm, respectively) (**Fig. 3B**). The BW of GF fish was significantly lower (7.67, 25.83, and 59.29 mg at 1, 2, and 3 mpf, respectively) than that in the CR group (11.18, 45.71, and 114.93 mg, respectively) (**Fig. 3B**). According to the morphometric hallmarks of marine medaka (**Fig. 3C**), the fish length between parts was measured by microscopy analysis software from the pictures (with 7.0 × and 2 mm scales). At 1 mpf, the LS, DHAD, DHDC, DHAA, DADAA, DAAPA, LFRsA, and LFRsD of GF fish have average values of 6.77, 5.51, 6.82, 3.82, 1.91, 1.44, 0.63, and 0.52 mm, respectively, and showed significant (*p*<0.05, 0.01, 0.001) inhibition compared to the CR groups, which had values of 9.26, 7.41, 9.16, 5.30, 2.66, 2.59, 1.23, and 1.09 mm, respectively. However, fin dimorphism was observed between males and females during development, and there were no obvious differences between the GF and CR groups in late larvae and juvenile fish (**Fig. 3D**). At 2 mpf, the LS, DHAD, DHDC, DHAA, DADAA, DAAPA, LFRsA, and LFRsD of GF fish were averaged to 9.54, 7.74, 9.52, 5.63, 2.66, 1.44, 1.00, and 1.02 mm, respectively, which significantly (*p*<0.001) lower compared to the CR groups with 14.07, 11.49, 13.96, 8.38, 4.20, 3.86, 1.78, and 1.93 mm. But, the fin dimorphism between males and females appeared to some extent in GF fish but not as obvious in CR fish at early adult stages (**Fig. 3E**). At 3 mpf, the average DHAD, DHAA, DADAA, DAAPA, LFRsA, and LFRsD of GF fish were 9.97, 7.20, 3.56, 3.43, 1.46, and 1.65 mm, respectively, which showed significant (*p*<0.001) inhibition compared to the CR groups with 17.68, 14.20, 7.50, 6.63, 3.45, and 3.12 mm, respectively. However, the fin dimorphism between males and females indicated developmental maturity, with obvious differences in both GF and CR fish at the adult life stage (**Fig. 3F**). The LS and DHDC of CR-3 marine medaka could not be measured by computer software because the adult fish were too large to take an unabridged photo in the minimum magnification horizon. However, the GF fish were smaller and could be photographed in one horizon, with the LS and DHDC determined as being 12.13 mm and 12.20 mm, respectively, at 3 mpf. Examination of the fish images indicated that development of the GF marine medaka from the larval to juvenile and adult stages was significantly delayed with distinctly smaller in size than the CR fish at the same life stages, which was consistent with the developmental indexes of BL and BW, and LS, DHAD, DHDC, DHAA, DADAA, DAAPA, LFRsA, and LFRsD. As of the time of sampling, the GF fish at 3 mpf could not spawn, while the CR fish began to produce embryos from 2.5 mpf, indicating an at least half-month of developmental delay between them.

**Figure 3.**
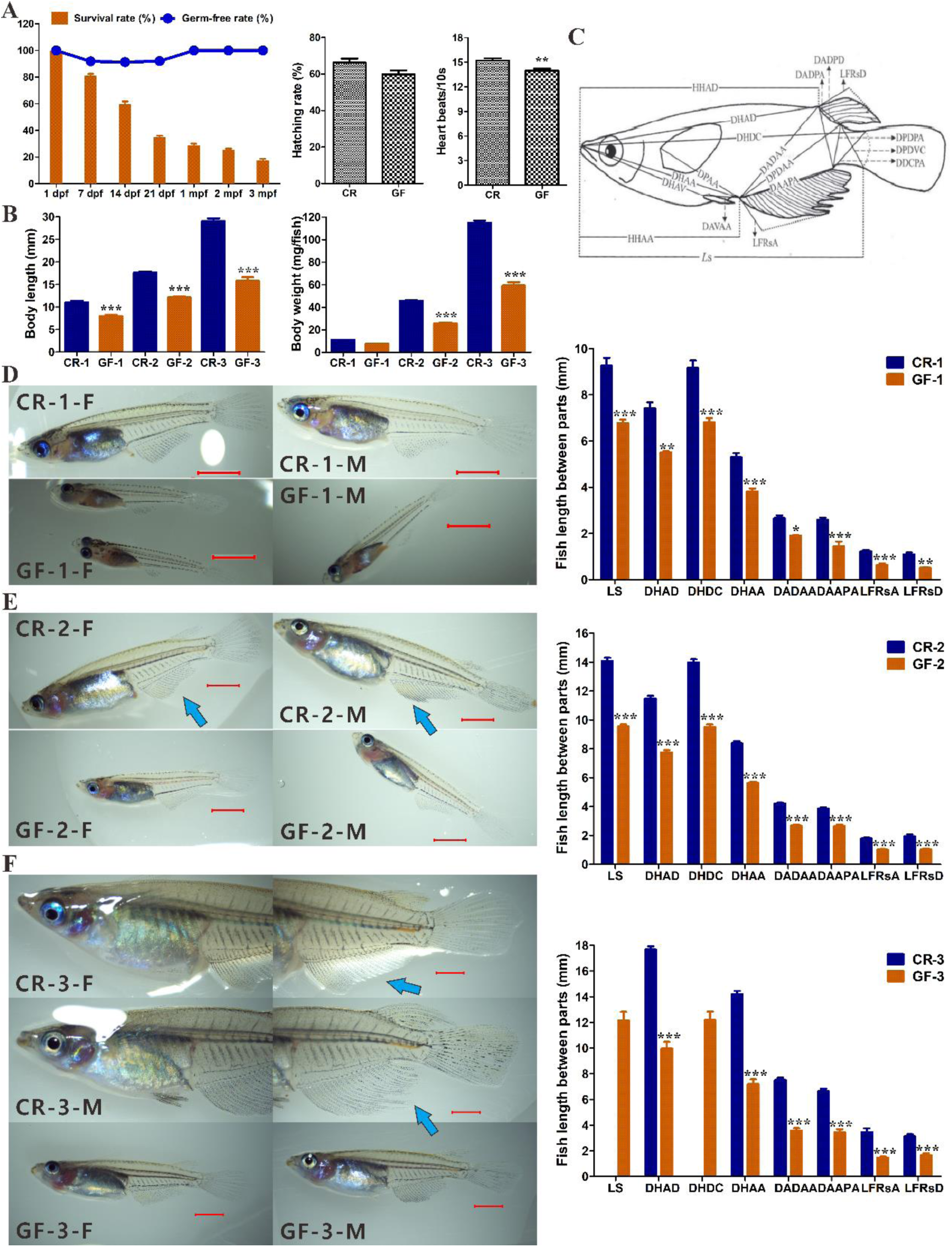
Developmental indexes of GF and CR marine medaka at different life stages. (A) The survival rate and germ-free rate of GF fish at 1, 7, 14, and 21 dpf, and 1, 2, and 3 mpf, the hatching rate of CR and GF at 13 dpf, and the heart beats/10s of CR and GF fish at 1 mpf. (B) The body length (mm) and body weight (mg) of the different life stages of marine medaka (CR-1, GF-1, CR-2, GF-2, CR-3, GF-3). (C) The morphometric measurements between each landmark for the marine medaka were analyzed as described by Jae *et al*. (D-E-F) Images (with 7× and 2 mm scale) of female and male marine medaka from juvenile to adult stages under a microscope with a white light background. The basic fish length between parts, including the LS, DHAD, DHDC, DHAA, DADAA, DAAPA, LFRsA, and LFRsD were recorded and compared the GF and CR groups. The LS and DHDC of CR-3 marine medaka were impossible to be measured by computer software, because the adult fish were too large to take an unabridged photo in the minimum magnification horizon. Among them, the body length was the total length from head to tail, and the LS was standard length. The DHAD was the direct distance between the most anterior extension of the head and the anterior insertion of the first dorsal fin. The DHDC was the direct distance between the most anterior extension of the head and the dorsal base of the caudal fin. The DHAA was direct distance between the most anterior extension of the head and the anterior insertion of the first anal fin. The DADAA was the direct distance between the anterior insertion of the first dorsal fin and the anterior insertion of the first anal fin. The DAAPA was the direct distance between the anterior insertion of the first anal fin and the posterior insertion of the last anal fin. The LFRsA was the length of the fin rays of the anal fin, and the LFRsD was the length of the fin rays of the dorsal fin. The symbols *, **, and *** represent for *p*<0.05, 0.01, and 0.001, respectively, with significant differences between CR and GF models at the same life stages.

### Behavioral differences between the GF and CR marine medaka models

To explore the developmental differences and effects on host health of GF animals, the behavioral differences of GF marine medaka compared to CR fish at 1 mpf were examined with a total of n=21 replicates in each group (shown in **Fig. S8**). In detail, the fish samples were placed under the same conditions and parameters under video tracking with EthoVision XT, and the results were then analyzed (**Fig. S8A**). The GF fish appeared to have less locomotion ability than the CR fish as shown by the visual images and the heatmap of the fish tracking data from 10 min experimental collection (**Fig. S8 B** and **C**). According to the 10-min tracking data, the average distance moved for the GF fish (3675.81 mm) was significantly less than that for the CR fish (5868.30 mm), and the average velocity of the GF fish (6.13 mm/s) was significantly slower than that of the CR fish (9.78 mm/s) (**Fig. S8D** and **E**). However, there were no obvious differences in the max and min accelerations between GF and CR groups with values of 1042.32 and 1170.04, -699.94 and -699.85 mm/s^2^, respectively (**Fig. S8F** and **G**).

For the average activity, the value of the GF fish (20.22%) was lower, but not significantly, than that of CR groups (26.14%) (**Fig. S8H**). Moreover, the static and active frequencies of GF marine medaka were 806.62 and 1096.00, which were both significantly lower than those of CR fish with 1065.10 and 1518.05, respectively (**Fig. S8I** and **J**). However, the manic frequency of the GF fish was not different from that of the CR fish (**Fig. S8K**). Notably, the cumulative duration for medaka compared to the total time of 10 min was calculated, and showed that the GF fish remained in the static state (63.37%) significantly longer than the CR fish (47.59%), and the GF fish remained in the active state (31.61%) significantly less time than the CR fish (45.47%) (**Fig. S8L**). Similarly as the manic frequency, the GF fish stayed in manic state (5.02%) lower but not significantly so than that of the CR group (6.94%) (**Fig. S8L**). All the results in this study proved that the behavioral ability of the GF fish was significantly weaker than those of the CR model, which probably contributed by the delayed development of GF marine medaka from the larvae to juvenile stages. Due to the limited number of fish, the behavior ability of the adult GF and CR models was not carried out, but research on this aspect is still needed in future studies. On the other hand, the absence of microbiota may lead to no effects on the manic state or highly mobile ability of GF fish at the juvenile stage when detected in normal water conditions without any outside light or noise stimulations.

### Histopathological analysis of GF and CR fish from the larval to adult stages

In this study, histopathological analysis of GF and CR marine medaka at 1, 2, and 3 mpf was performed with totally >180 sections (n=3 replicate samples for each group, and n=5 sections with a special stain for each sample), to explore the changes that occur in fish during the developmental process (**Fig. S9**). The later larvae to juvenile fish at 1 mpf from the CR and GF models were all in the state of growth and development, and could be separated by sex based on the developing gonads observed in the sections (**Fig. S9A** and **B**). Although the CR-1 and GF-1 fish could be observed in the same magnification of 50×(scale bar of 400 μm), the GF fish were still smaller and thinner, with sparse muscle arrangement, swim bladders and liver tissues than CR marine medaka at 1 mpf. Moreover, the early adult fish at 2 mpf showed developed tissues or organs, including the brain, eye, gill, kidney, peritoneum, pancreas, anal fin, gonad, liver, heart, gut and cloaca, and the developmental differences were significant between CR fish with 20×, 1.25 mm, and GF fish with 30×, 625 μm sections (**Fig. S9C** and **D**). The adult CR fish with 10× and 2.5 mm scale bar sections at 3 mpf exhibited completely developed tissues to be the mature ones, including the brain, eye, gill, kidney, anal fin, gonad, liver, heart, gut and cloaca, but GF fish with 20× and 1.25 mm sections still exhibited a distinctly smaller size and delayed maturation of organs (**Fig. S9E** and **F**). The muscle, gill, spinal cord, backbone, and, especially, ovary and testis of the CR fish were observed to be fully developed in terms of size, count, and density indexes, compared to those of the weak GF fish (Af: anal fin, B: brain, Bb: backbone, C: cloaca, E: eye, G: gill, Gu: gut, H: heart, K: kidney, L: liver, M: muscle, O; ovary, P: peritoneum, Pa: pancreas, S: swim bladder, Sc: spinal cord, Sp: spleen, T: testis, To: tongue,♀: female, ♂: male). Notably, the cloaca, which is connected to the kidney duct, gonad and gut, was increasingly developed along with CR and GF fish development from the 1 to 2 and 3 mpf stages, indicating the development of the fish body and system functions, especially after the spawning of early adult CR fish from 2.5 months.

Moreover, the developmental differences in major immune organs, including the liver, kidney, pancreas, and spleen, were investigated and compared between the GF and CR marine medaka from the juvenile to adult models (**Fig. S10**). The liver tissue of GF and CR fish at 1, 2, and 3 mpf, which showed different degrees of development during growth and between groups, indicated a decrease in hepatocyte levels and low hepatic density in GF fish along with development and growth (**Fig. S10A**). The kidney tissue of the GF and CR models showed a distinct and mature structure in normal fish head and trunk kidneys, but fewer tubules and hematopoietic tissue in GF fish. Moreover, in the kidney, the CR fish showed mature glomeruli, proximal and distal tubules, collecting ducts, hematopoietic tissue, and newly generated renal tubules, but the GF fish seemed to have a weak ability to grow renal tubules and showed less hematopoietic tissue in kidney tissues (**Fig. S10B**). The pancreas in both GF and CR fish had relatively intact and small tissues in at 1 mpf, but discrete multiple pancreas were scattered along the intestinal tract in GF and CR fish at 2 and 3 mpf, and a smaller area of GF fish pancreas and several vacuoles appeared in both GF and CR fish (**Fig. S10C**). The spleen of GF and CR fish at 3 mpf showed developed and relatively intact tissues near the gut and peritoneum, but that of the GF fish was smaller and showed an indistinct difference in red pulp (R) and white pulp (W), and more melano-macrophage centers (MMCs) in tissues, compared to that of the CR fish (**Fig. S10D**). However, not enough spleen sections were obtained at 1 and 2 mpf for the GF and CR fish, possibly due to the small size of the spleen before development to adult fish.

The developmental characteristics of the intestinal tissues of GF and CR fish were identified by histopathological analysis, and the critical indexes were measured from the juvenile to adult models (**Fig. 4**). In juvenile fish at 1 mpf, the anterior-middle and middle-posterior intestines of the GF fish were smaller than those of the CR fish, and the villus height, width and muscular thickness of the GF fish were slightly weaker, but not significantly, except for the villus width of the anterior-middle intestines, indicating the progression of development in both the GF and CR groups. The average values of villi height, width and muscularis thickness of the CR-1 and GF-1 fish anterior-middle intestines were 88.69, 85.74, 63.49, 49.60, 8.61, and 8.14 μm, and the indexes for the CR-1 and GF-1 middle-posterior intestines were 89.74, 87.76, 48.95, 48.81, 9.85, and 9.33 μm (**Fig. 4A**). In early-adult fish at 2 mpf, the anterior-middle and middle-posterior intestines of the GF fish were significantly different (*p*<0.01 and 0.001) from those of the CR fish except for the muscularis thickness of the anterior to middle intestine. The villi height, width and muscularis thickness of the GF fish seemed obviously lower, with a narrow enteric cavity and the villi width of anterior-middle intestines, indicating the serious developmental delay in gut tissue between the GF and CR groups. The average values of villi height, width and muscularis thickness of the CR-1 and GF-1 fish anterior-middle intestines were 102.19, 87.97, 69.29, 46.52, 12.28, 12.01 μm, and the indexes for the CR-1 and GF-1 middle-posterior intestines were 110.77, 93.49, 68.01, 56.94, 13.20, 10.74 μm (**Fig. 4B**). In adult fish at 3 mpf, the anterior-middle and middle-posterior intestines were observed to be fully developed with numerous and curly villi in both the CR and GF fish. However, the villi height, width and muscularis thickness of GF fish were significantly lower than those of the CR fish, except for the villi width of the anterior-middle intestines. The average values of villi height, width and muscularis thickness of the CR-1 and GF-1 fish anterior-middle intestines were 125.77, 105.21, 62.27, 60.10, 20.41, and 11.00 μm, and the indexes for the CR-1 and GF-1 middle-posterior intestines were 130.12, 116.12, 58.59, 45.28, 23.22, and 15.24 μm (**Fig. 4C**). The intestinal indexes from the histopathological sections showed a backward or slow increase in the GF fish and significant differences from the CR models during growth, indicating that the gut microbiota may contribute to the deficiency in intestinal system development and weak digestive or absorption ability.

**Figure 4.**
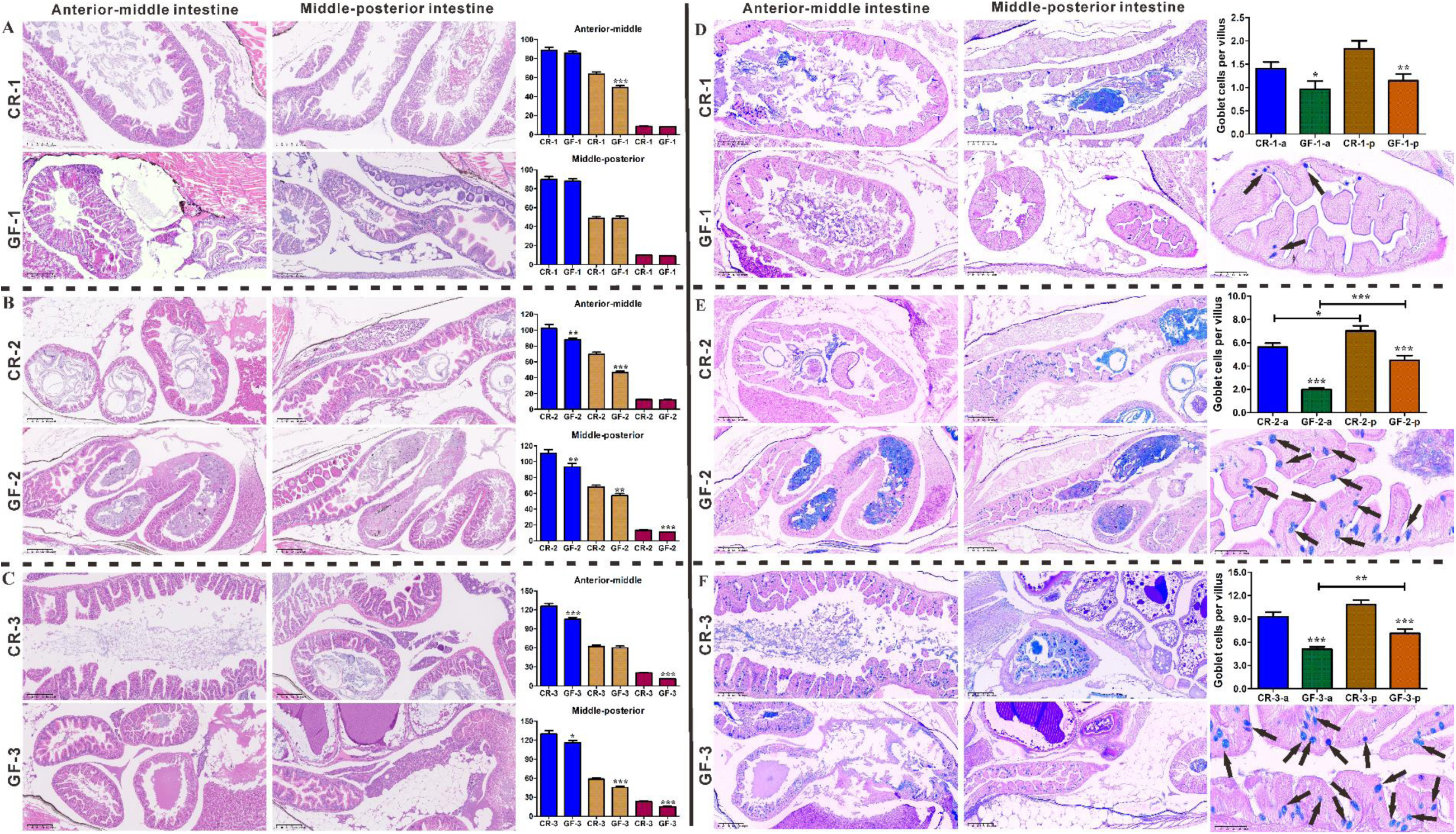
Micro-structure, distribution of mucous layers, and goblet cells of intestines in GF and CR marine medaka from juvenile to adults. (A) The anterior to middle intestines, and middle to posterior intestines of GF and CR fish at the 1 mpf juvenile stage, and the developmental indexes of intestines including the villus height, villus width, and muscularis thickness were measured from sections with magnification 200× and scale bar of 100 μm. (B) The anterior to middle intestines, and middle to posterior intestines of GF and CR fish at the 2 mpf early-adult stage, and the developmental indexes of intestines including the villus height, villus width, and muscularis thickness were measured from sections with magnification 100× and scale bar of 200 μm. (C) The anterior to middle intestines, and middle to posterior intestines of GF and CR fish at the 3 mpf adult stage, and the developmental indexes of intestines including the villus height, villus width, and muscularis thickness were measured from sections with magnification 100× and scale bar of 200 μm. Comparisons of the villus height (blue color), villus width (brown color), and muscularis thickness (deep red color) data were presented in histograms. (D) The anterior to middle intestines, and middle to posterior intestines of GF and CR fish at 1 mpf juvenile stage, the mucous layers and the goblet cells were highlighted by AB-PAS stain, and mucus-secreting goblet cells were measured from sections with magnification 200× and scale bar of 100 μm. (E) The anterior to middle intestines, and middle to posterior intestines of GF and CR fish at 2 mpf, the mucous layers and the goblet cells were highlighted by AB-PAS staining, and mucus-secreting goblet cells were measured from sections with magnification 100× and scale bar of 200 μm. (F) The anterior to middle intestines, and middle to posterior intestines of GF and CR fish at the 3 mpf adult stage, and the goblet cells per villus in gut tissues were measured from sections with magnification 100× and scale bar of 200 μm. There were n=9 sections of three biological replicates in each group, for a total of n=54 villi detected from the anterior-middle and middle-posterior intestines without sex tranches. Note that the magnified posterior intestines were GF-1, GF-2, and GF-3 with 500× magnification and scale bar of 50 μm, in which the black arrow indicted that the goblet cells were highlighted blue by AB-PAS stain. The measured goblet cells in the villi of CR anterior to middle intestines (CR-1-a, CR-2-a, CR-3-a, blue color), middle to posterior intestines (CR-1-p, CR-2-p, CR-3-p, green color), GF anterior to middle intestines (GF-1-a, GF-2-a, GF-3-a, brown color), and middle to posterior intestines (GF-1-p, GF-2-p, GF-3-p, green color) were presented in histograms. The goblet cells (black arrow) showed the goblet-like shape with mucous droplets filling up the cytoplasm and the mucous released into the gut lumen, and the symbols *, **, and *** stand for *p*<0.05, 0.01, and 0.001, respectively, with significant differences in goblet cell number per villus between CR and GF fish intestines.

Furthermore, the photomicrographs of the mucous layers and goblet cells in gut tissue of the CR and GF fish were subjected to AB-PAS staining (**Fig. 4**). In this study, it observed that the mucous layers in both CR and GF marine medaka at 1 mpf juvenile stage were filled with purple-red neutral mucins in the anterior to middle intestine. The more acid and mixed mucins stained with blue and blue-purple appeared in the middle to posterior intestines, but the diverse mucins were obviously more abundant in the CR fish than in the GF fish intestines. Moreover, the number of goblet cells highlighted by AB-PAS staining in the anterior to middle intestines and middle to posterior intestines of the GF fish, with an average of 0.963 and 1.148 cells per villus, was significantly decreased compared to that in CR fish, with 1.407 and 1.833 cells per villus (**Fig. 4D**). The mucous layers in marine medaka at 2 mpf early-adult stage showed a higher abundance and also purple-red neutral mucins in anterior intestine, and more acid and mixed mucins in middle to posterior intestines stained with blue and blue-purple, but the mixed mucins seemed to be the dominant type in the GF fish middle to posterior intestines. Similarly, the average number of goblet cells in the anterior to middle intestines was 1.981 cells per villus, and middle to posterior intestines of GF fish was 4.500 cells per villus, which was significantly reduced compared to that in CR fish with an average of 5.630 and 7.012 cells per villus, respectively (**Fig. 4E**). The posterior gut showed more goblet cells with longer villus and higher absorption function requirements. At 3 mpf, the mucous layers in adult CR marine medaka showed purple-red neutral mucins and blue-purple mixed mucin in the anterior intestine, and abundant acid and mixed mucins in the middle to posterior intestines. But, the neutral mucin in the anterior intestines and mixed mucins in the posterior intestines seemed to be the dominant type and were less distinct in GF fish. Compared to those in younger fish, the goblet cells in the anterior to middle intestines and middle to posterior intestines in both the GF and CR fish were increased, but the average of 5.09 and 7.13 cells per villus in GF fish was still significantly less than that in CR fish (9.278 and 10.833 cells per villus), respectively (**Fig. 4F**). The goblet cells (black arrow) showed a goblet-like shape with mucous droplets filling up the cytoplasm and the mucous released into the gut lumen. All these results provided strong evidence that the planting of the gut microbiota in fish critically influences intestinal development, goblet cells and mucin productions, and subsequent digestive functions.

In this study, the developmental differences in gonad tissues between GF and CR fish were also explored to reveal the impacts of gut microbiota deficiency on reproductive system (**Fig. 5**). The histopathological analysis of ovary and testis was referred to previous reports (5) and the “Guidance document on the diagnosis of endocrine-related histopathology in fish gonads” in the OECD Series on Testing and Assessment (No.123). In juvenile fish, the identified growing ovary tissues of GF and CR female fish exhibited primary germ cells in the early stages I (oogonia) and II (previtellogenic oocytes), and distinct testis tissues were observed in CR and GF male fish at 2 mpf. However, the germ cells in GF developing gonads were observed mainly the spermatogonia (SG) and spermatocytes (primary and secondary, SC) stages, compared with the almost fully grown testis in size and matured sperm in CR fish indicating the early-adult stage (**Fig. 5A**). The ovary tissues in GF fish at 3 mpf were immature, with oocyte number in stages I, II, III (vitellogenic oocytes) and IV (postvitellogenic oocytes) detected to be significantly decreased with an average of 3.13, 5.73, 2.33 and 1.27 cells in the set area of gonad tissue, compared with that in the CR fish (6.47, 13.0, 7.73 and 3.4 cells). Moreover, the percentages of the I, II, III and IV stages in GF fish were calculated to be 26.27, 43.94, 18.91, and 10.88%, respectively, compared with those in the CR fish (4.66, 42.31, 26.60, and 26.44%). These results indicated that the oocytes in the GF fish were immature at the early I stage and significantly fewer in number at III stage (**Fig. 5B**). The average number of germ cells in the set area of testis tissues in GF fish at 3 mpf was also significantly lower in the SG, SC, spermatids (ST) and spermatozoa (SZ) stages with the detected 4.60, 169.40, 119.33, and 92.53 cells detected, than in CR fish at the adult stage with 17.00, 228.33, 142.20, and 151.40 cells. Meanwhile, the percentages for the SG stages were significantly different between the GF and CR fish at 1.21 and 3.18%, respectively (**Fig. 5C**). Moreover, the total number of cells at different stages in the set area in ovary at the 3 mpf adult stage was counted to be 12.47 in GF fish, which was significantly less than that (30.6) in CR fish (**Fig. S11**). Similarly, the total cells in testis at different stages were calculated and significantly less (385.87) in GF fish, than that (538.93) in CR fish (**Fig. S11**).

**Figure 5.**
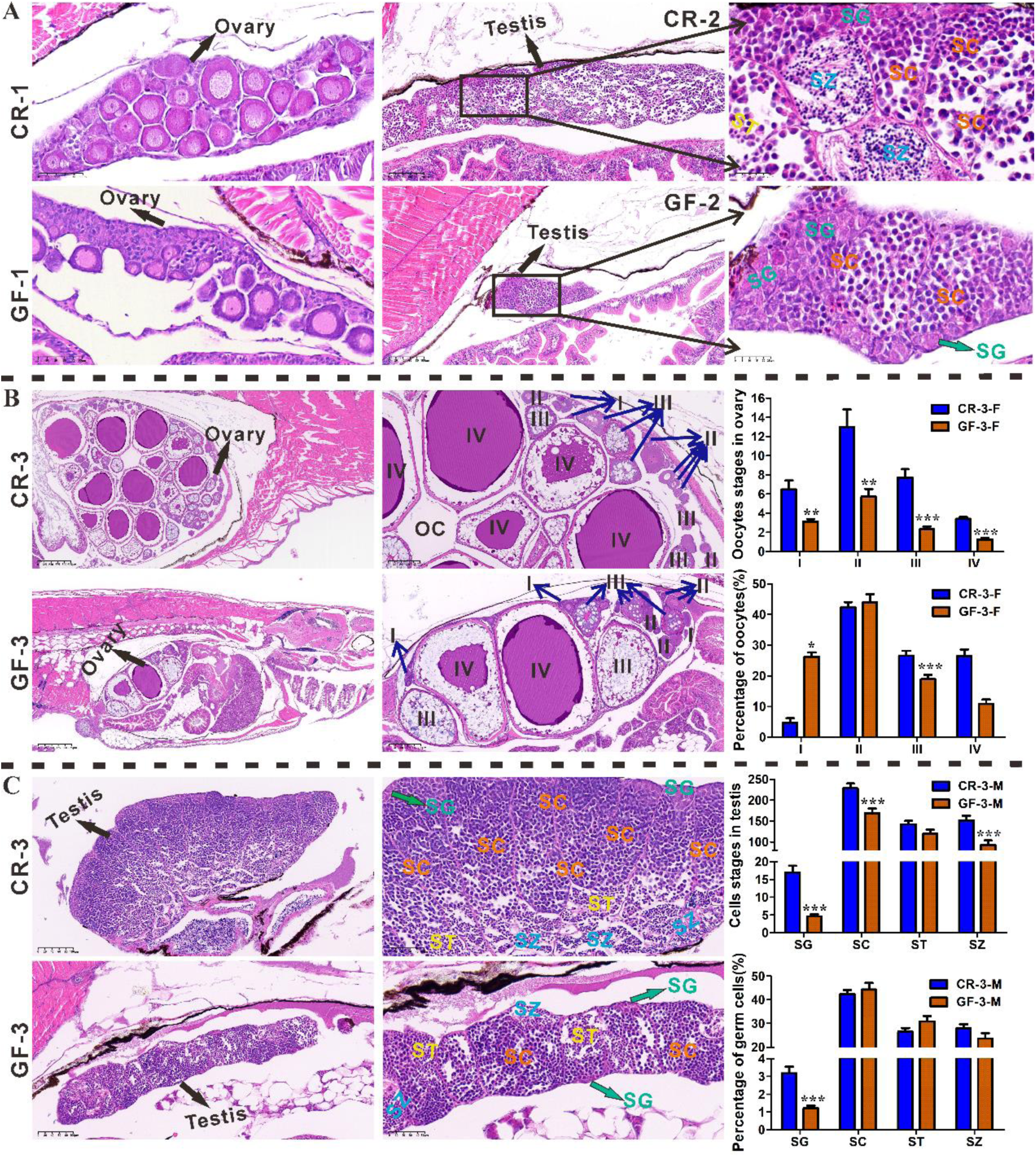
Developmental differences in reproductive systems between GF and CR marine medaka from juvenile to adult models. (A) The identified growing ovary tissues of GF and CR female fish at 1 mpf, which appeared with primary germ cells and the early stages I and II of oocytes in sections with 500× magnification (scale bar, 50 μm). The distinct testis tissues in CR and GF male fish at 2 mpf were represented in sections with 200× magnification (scale bar, 100 μm). The stages of germ cells in the developing gonad of GF fish were observed mainly at the SG and SC stages, compared with the almost fully grown testis in terms of size and cell stages in CR early-adult fish sections with 800× magnification (scale bar, 25 μm). (B) The ovary tissues in CR and GF fish at 3 mpf with 30× magnification (scale bar, 625 μm) showed mature and immature differences, with the numbers in the I, II, III and IV stages of oocytes in ovaries detected in sections with 90× magnification (scale bar, 200 μm) sections and the percentage calculated. (C) The testis tissues of CR and GF fish at 3 mpf with 200× magnification (scale bar, 100 μm) also showed significant differences, with the numbers in the SG, SC, ST and SZ stages of germ cells in testis detected in magnified 400× (scale bar, 50 μm) sections and the percentage analyzed. Ovary: I=oogonia, II=previtellogenic oocytes, III=vitellogenic oocytes IV=postvitellogenic oocytes, OC=ovarian cavity. Testis: SG=spermatogonia, SC=primary and secondary spermatocytes, ST=spermatids, SZ=spermatozoa, green arrow=SG. The symbols *, **, and *** stand for *p*<0.05, 0.01, and 0.001, respectively, with significant differences in germ cell number and percentages at various stages in the set area of gonad tissues between CR and GF fish separated by sex.

### Transcriptomic profile of GF and CR marine medaka at early-adult and adult stages

In this study, the transcriptomic profile of GF and CR fish was explored to illustrate the control mechanisms of early-adult and adult stages between different groups, which potentially provide the understanding of gut microbiota effects on fish growth and development. There were four different groups, namely, GF-2, CR-2, GF-3, and CR-3, with 4 replicate samples of each group, among which 2 females and 2 males were selected at 3 mpf. A total of 16 samples yielded with 934,879,796 clean reads and 28599 expressed genes. The reference genome used for analysis was that of *O. melastigma* (GCF_002922805.2) from https://www.ncbi.nlm.nih.gov/genome/?term=txid30732, and the total clean reads were mapped with 91.87-94.16% indicating the suitability of the reference genome and no experimental contamination of the samples. The functional annotation of reference genes in GF and CR groups at different developed stages was performed based on the databases included the GO, KEGG, COG, NR, Swiss-Prot, and Pfam (**Fig. S12**). The coverage of sequencing reads in each sample and the distribution of the sequencing reads showed the higher levels of expression profile in CR fish (**Fig. 6A** and **B**). In the PCA, different groups were presented, in which a similar distance appeared between the GF-3 and CR-2 fish (**Fig. 6C**). The commonly and specially expressed genes in the GF-2, CR-2, GF-3, and CR-3 groups were identified (**Fig. 6D**). The significant different expressed genes (DEGs) both up- and down-regulation in the compared groups of GF-2-VS-CR-2, GF-3-VS-CR-3, GF-3-VS-GF-2, and CR-3-VS-CR-2 were selected by using the *p*-adjust value <0.05 and FC≥2 or FC≤0.5. In the GF-2-VS-CR-2 group, a total of 1030 DEGs with 294 up-regulated and 736 down-regulated genes were screened, while in GF-3-VS-CR-3 group, totally 4046 DEGs with 1824 up-regulated and 2222 down-regulated genes were screened(**Fig. 6E** and **F**). A total of 1662 DEGs with 662 up- and 970 down-regulation appeared in the GF-3-VS-GF-2 comparison group, and totally 4530 DEGs with 1780 up- and 2750 down-regulation appeared in the CR-3-VS-CR-2 group (**Fig. 6G** and **H**). Among these DEGs, the important genes searched by key words “growth”, “development”, “gonad”, and “reproduction” from the various compared sets were selected and summarized in **Table S10**, which showed the fold changes in the GF and CR at different life stages and the related signaling pathways in marine medaka. Briefly, the *gsdf*, *igfals*, *dio2*, *tgfb5*, *egfl7*, *cgref1*, *igf1*, *gh1*, *esm1*, *epcam*, and *junbb* genes were all significantly down-regulated in GF-2 fish compared to CR-2 fish, which were related to the growth hormone synthesis, secretion and action, thyroid hormone signaling pathway, ovarian steroidogenesis, and PI3K-Akt signaling pathways (**Table S10**). In the GF-3-VS-CR-3 group, the genes *gsdf*, *fgf16*, *igfals*, *tgfbr2b*, *pdgfc*, *gdf6a*, *igf1*, *fgf23*, *pdgfrb*, *fgfr2*, *egr1*, *pkd2*, *fgf16*, *ngfb*, *gcm2*, *mmp14b*, *spaca4l*, *six5*, and *mmp14a* were significantly decreased. These key DEGs related to the growth hormone synthesis, secretion and action, thyroid hormone signaling pathway, ovarian steroidogenesis, PI3K-Akt signaling pathway, parathyroid hormone synthesis, secretion and action, GnRH and TNF signaling pathways (**Table S10**). Interestingly, in the GF-3-VS-GF-2 group, the genes *pdgfrl*, *tgfbr2b*, *tgfb5*, *mmp14b*, and *gnrh3* were significantly up-regulated. These key DEGs related to the cell cycle, pancreatic cancer, TGF-beta signaling pathway, parathyroid hormone synthesis, secretion and action, GnRH and TNF signaling pathways, while the genes *socs2* and *fgf1a* were significantly inhibited and were related to the growth hormone synthesis, secretion and action, and PI3K-Akt signaling pathways (**Table S10**). In the CR-3-VS-CR-2 group, the genes *pdgfrl*, *fgf7*, *fgfbp2a*, *tgfb1a*, *tgfb5*, *gdf10a*, *gdf6a*, *ngfra*, *fgf23*, *ngfb*, *mmp14b*, *bmp4*, and *mmp14a* were commonly significantly up-regulated. These key DEGs related to the cell cycle, growth hormone synthesis, secretion and action, parathyroid hormone synthesis, secretion and action, GnRH signaling pathway, and thyroid hormone signaling pathways. While, the genes *gadd45ga*, *socs3a*, *gadd45ba*, *gh1*, and *isl1*were significantly decreased and were related to the TNF signaling pathway, Jak-STAT signaling pathway, insulin signaling pathway, growth hormone synthesis, secretion and action pathways (**Table S10**).

**Figure 6.**
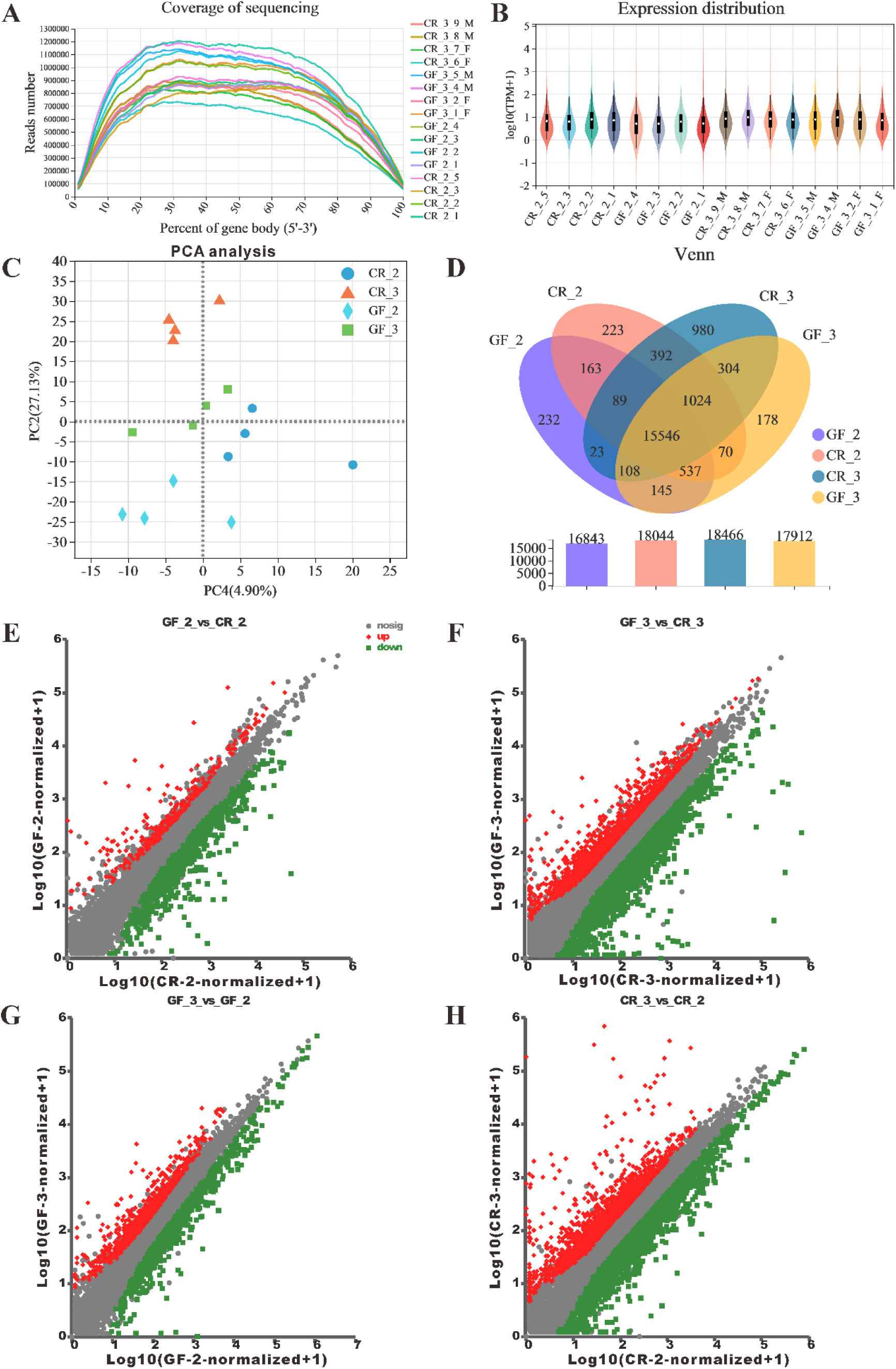
Transcriptomic analyses of GF and CR marine medaka at early-adult and adult stages. (A) The coverage of sequencing reads in each sample and the distribution (B) of the sequencing reads showed the higher-level expression profile of CR fish. (C) The PCA of different group samples showed a similar distance between GF-3 and CR-2 fish. (D) The common and special genes expressed in the GF-2, CR-2, GF-3, and CR-3 groups. (E-F-G-H) The significant different expressed genes (DEGs) with up- and down-regulation in the 4 comparison groups (GF-2-VS-CR-2, GF-3-VS-CR-3, GF-3-VS-GF-2, CR-3-VS-CR-2) selected based on the *p*-adjust value <0.05 and FC≥2 or FC≤0.5.

The annotation analysis of the GO database at level 2 showed the major terms enriched in the molecular function, cellular component, and biological process categories, which indicated the DEGs in the GF-3-VS-CR-3 and CR-3-VS-CR-2 groups (**Fig. S13A**). However, the summary of alteration splicing (AS) type in GF and CR fish clearly showed more skipping exons (SEs) at 2 mpf early-adult stage than at 3 mpf adult stage (**Fig. S13B**). The top 25 enriched KEGG pathways in GF-2-VS-CR-2 involved the Complement and coagulation cascades, ECM-receptor interaction, Focal adhesion, and Protein digestion and absorption, and especially Ovarian steroidogenesis (**Fig. S13C**). In the GF-3-VS-CR-3 group, a total of 17 significantly changed pathways were closely related to the Protein digestion and absorption, ECM-receptor interaction, Complement and coagulation cascades pathways (**Fig. S13D**). KEGG pathway enrichment between GF-3 and GF-2 fish showed totally 7 significantly changed pathways of Phototransduction and Protein digestion and absorption (**Fig. S13E**). The enriched KEGG pathways showed top 25 of totally 37 pathways in the CR-3-VS-CR-2 group, which involved in the Protein digestion and absorption, Nicotine addiction, Glutamatergic synapse, ECM-receptor interaction, PI3K-Akt signaling pathway, and Pancreatic secretion (**Fig. S13F**). Those enriched KEGG pathways were selected from the DEGs sets and are critical to organism growth, immune, and reproductive systems. Furthermore, the critical KEGG pathway of Ovarian steroidogenesis was significantly enriched in the GF-2-VS-CR-2 group, which showed the down-regulation of *insulin*, *igf1*, *cyp17*, and *cox2* genes in the GF-2 fish reproductive system (**Fig. S14A**). The protein digestive and absorption pathway was the most changed in the GF-3-VS-CR-3 group and CR-3-VS-CR-2 group (**Fig. S14B** and **C**). Importantly, the DEGs in the lumen including the intestines were down-regulated and up-regulated in cells to blood in GF-3 fish, compared to CR-3 fish. However, the expression of DEGs in the lumen including the intestines and blood was conversely expressed in CR-3 fish, compared to CR-2 fish, indicating the difference in gene expression between GF and CR, and among adult fish for protein digestion and absorption (**Fig. S14B** and **C**). Among the enriched KEGG pathways, the critical KEGG pathway of Phototransduction in the GF-3-VS-GF-2 group was significantly changed (**Fig. S15**). The Pi3k-akt pathway and the pancreatic secretion pathway were the significantly changed in the CR-3-VS-CR-2 group; these pathways closely related to the development and growth, intestinal functions and metabolism at different growth stages (**Fig. S16** and **S17**).

## Discussion

The ocean environment and pollution toxicity have increasingly attracted attentions in recent years, and the development and application of marine medaka as powerful animal models are urgently needed. Our study provided insights into the distribution and contribution of the intestinal microbiota in marine medaka at different life stages, as well as the physiological differences in the generated GF models compared with CR fish during growth and development. Moreover, the transcriptomic profiles of marine medaka from various groups and life stages were analyzed in depth to illustrate the potential effects and mechanisms of the intestinal microbiota on fish development from larval to juvenile and adult stages, as well as the underlying mechanisms.

In this study, the diversity and richness of the gut microbiota of CR marine medaka during different life stages were examined, and the distributions in female and male fish were identified. It proved that the dominant phyla of the gut microbiota from the juvenile to adult stages were Proteobacteria, Verrucomicrobia, Actinobacteria, Firmicutes, Bacteroidetes, and Cyanobacteria, most of which were decreased in both female and male marine medaka during growth. Furthermore, the domainant changed genera of the gut microbiota, including *Ruegeria*, *Haloferula*, *Sulfitobacter*, and *Vibrio*, belonging to phyla of Proteobacteria, Verrucomicrobia, Actinobacteria and Bacteroidetes were enriched, while *Pseudomonas*, *Leisingera*, *Ahrensia*, and *Ralstonia* belonging to phylum of Proteobacteria and Cyanobacteria were depleted. Interestingly, these significantly changed genera indicated bacterial appearance or disappearance during various life stages, with the major genera *Haloferula*, *Ruegeria*, *Iamia*, *Nautella*, *Actibacter*, and *Marimicrobium* being significantly and positively related to the BL, BW, BMI and ISI development indexes. The prediction of the gut microbiota in marine medaka with microbial phenotypes provided novel insights into bacterial functions and their isolation and subsequent identification of bacteria from fish intestines. The intestinal bacteria primarily contributed to the Gram-Negative, Gram-Positive, Biofilm Forming, Pathogenic, Mobile Element Containing, Oxygen Utilizing including Aerobic, Anaerobic, Facultatively-Anaerobic, and Oxidative Stress Tolerant phenotypes and can be applied for regulation and as probiotics in bio-medicine fields. All of these results indicated the important factors related to host health, growth, and functional changes, which can be evaluated to explain the mechanism underlying the effects of the gut microbiota along with the age and sex dependent manners.

A higher diversity of the intestinal microbiota was observed at 1 mpf, which may help fish adapt to changes in the external environment and promote growth and development. From 3 to 5 mpf, the dynamic changes in the intestinal microbiota in marine medaka tended to be stable during the adult period. Leon Cantas *et al.* researched on the culturable gut microbiota diversity and reported the totally 13 morphologically different bacterial species in zebrafish, which mainly belonged to *Aeromonas*, *Vibrio*, *Pseudomonas*, and *Staphylococcus* genera (14). Here, we isolated totally 22 bacterial strains belonging to 10 genera, among which the major strains of *Pseudomonas*, *Shewanella*, and *Vibrio* belonged to the dominant phylum Proteobacteria in marine medaka, according to morphological identification, 16S rRNA gene-based sequencing, and an the NCBI-BLAST search against various databases. However, the culturable gut bacterial strains in marine medaka were screened under aerobic and facultatively anaerobic conditions, and the sttrictly anaerobic bacteria in the gut microbiota should also be taken into consideration in the future.

Live food products such as *Artemia* serve as vectors for delivering compounds of diverse nutritions for larval stages of aquatic animals and are the most important factors influencing the survival rate and growth from the early larval to the juvenile stages, when many digestive enzymes are secreted before feeding but with relatively weak activities (33, 34). Among the common feeds, the *Artemia* nauplii as the biological feed providing exogenous digestive enzymes and a high abundance of nutrients for fish larvae, are the most suitable live feed compared to cooked egg yolk and micro-particle feed; and the nauplii are essential for good growth and development, resistance to disease, and reproduction performance (35–37). Due to the absence of a microbiome and the defects in digestive and immune systems, studies on GF fish models have been limited to the early life stages in which depended on the yolk energy supply without eating, such as in GF zebrafish with innate immunity before 7 dpf (22). In this study, the generation and application of GF *Artemia* for breeding the aseptic fish models was largely performed with simple hatching equipment and a simple operation process to prevent the continuous pollution *via* aeration, which is an important technical benefit for generating long-term GF fish models. Then, we designed a protocol for breeding GF marine medaka from larvae to adult fish, facilitating the application of animal models for studying microbial functions. In this protocol, the key steps were from the embryos treatment in sterile culture solution to the breeding of the GF marine medaka models from larvae to adult fish. During the progress of GF models generation, the detection of fish living conditions, waste water, and feeding samples collected at key life points was carried out to identify the success of obtaining GF fish. The detection of samples in aerobic and anaerobic conditions proved the success of GF *Artemia* models with this quick and effective protocol. In fact, the application of GF *Artemia* in feeding GF fish models daily was easy and suitable, which supplied the higher rich nutrient substance for animal survival at later life stages. Although the survival rate of GF fish at 2 and 3 mpf significantly decreased to less than 24.67 and 16.67%, respectively, compared to that of CR fish at the same life stages, the sufficient fish samples were successfully collected for subsequent histopathological and transcriptomic analyses.

During the development of GF medaka fish, the basic indexes were significant inhibited compared to those of CR fish at the same life stages. The average heart beats at 1 mpf, BL and BW of GF marine medaka at 1, 2, and 3 mpf were obviously lower than those of the CR groups, which were consistent with the morphometric characteristics of LS, DHAD, DHDC, DHAA, DADAA, DAAPA, LFRsA, LFRsD determined from the fish images. The GF marine medaka from larvae to juvenile to adults exhibited significant developmental delay and appeared distinct smaller sizes compare with CR fish at the same life stages. Notably, until the sampling, the GF fish at 3 mpf still could not spawn, while the CR fish began to produce embryos from 2.5 mpf, which also indicated an at least half-month delay in development between them. Possibly due to the developmental delay, the fish behavior proved that GF juvenile fish at 1 mpf appeared dramatically weaker movement ability in terms of the average distance moved, average velocity, average activity, and static and active frequency. Notably, the cumulative duration of GF medaka in the static state was significantly longer, and that in the active state was significantly shorter, than that of CR fish. Due to the limited number of surviving fish, the behavioral ability of the adult GF and CR models was not measured, but still needed in the future studies. On the other hand, we hypothesized that the absence of the microbiota may lead to impacts on nutrient absorption or energy metabolism in GF fish from juvenile, which contributed to the developmental and behavioral differences compared to CR fish.

Similar to mammals, medaka process an XY, YY sex-determination system with the male-determining locus on the Y chromosome, and show a distinct sexual dimorphism between female and male fish (38). In this study, it was difficult to distinguish the males and females of both GF and CR fish without obvious dimorphic dorsal fins during developmental stages from embryos/larvae to 1 mpf, until the later larvae to juvenile and to 2 mpf the early-adult fish with almost mature systems. The CR fish from early-adult showed distinct sexual dimorphism, wherein females had a flat distal surface of the anal fin, while that of males was convex due to longer fin rays in a portion of the structure. From the histopathological analysis, the development of the reproductive system was detected to be delayed in the female and male GF fish, compared to the normal CR fish. In later larvae to juvenile fish at 1 mpf, the CR and GF models all showed growth and development, and could be distinguished as females or males based on the developing gonads in the sections. The GF fish from the juvenile to “adult” stages were smaller and thinner with sparse muscle arrangement, swim bladders, liver tissues, pancreas, anal fin, gonad, gut and cloaca, which indicating that the absence of the gut microbiota probably delayed system functions, especially spawning ability of early adult fish from 2.5 months of age.

Marine medaka (*O. melastigma*) was considered as the potential marine fish model for studying immunity, with the development of immune organs and innate and adaptive immune defense from the larval stage (39, 40). In a previous study, the immune tissues and key organs in fish, such as mucosal tissues, spleen, head kidney, thymus and liver, were detected to evaluate host immune competence, among which the liver is a key innate immune organ and is larger than other immune organs in size (41). In this study, according to the histopathological analysis during growth, GF marine medaka showed developed organs at juvenile stage, and significantly different at early-adult, and finally delayed development at 3 months-old compared to CR fish. Here, the developmental differences in major immune organs, including the liver, kidney, pancreas, and spleen, were investigated and compared between GF and CR marine medaka from the juvenile to adult models.

In fish, the larval and juvenile to adult period are normally characterized by high energetic requirements to support high growth rates and striking transformations of tissues and organs (42). However, it was reported that in GF zebrafish and rodent animal models, absence of the microbiota reduced intestinal cell proliferation, including the goblet cells and associated immune cells, and disrupted metabolism and innate immunity, which suggesting the effects of the gut microbiota on host organs, especially on intestinal development (10, 11, 43). In this study, histopathological analysis of intestinal tissues showed differences between GF and CR models from juvenile to adult fish, providing insights into developmental influences of deficiency of gut microbiota. It was proven that at 1 mpf, the anterior-middle and middle-posterior intestines of GF juvenile fish were smaller than those of CR fish with the villi height, width and muscularis thickness of GF fish slightly lower. But the differences were not significant, except for the villi width of the anterior-middle intestines, indicating the developmental progress in both GF and CR groups. In early-adult fish at 2 mpf, the anterior-middle and middle-posterior intestines of GF fish were significantly different from those of CR fish, except the muscularis thickness of the anterior to middle intestine part. The villi height, width and muscularis thickness of GF fish seemed obviously weaker with narrow enteric cavity, which indicated the serious developmental gap of gut tissue between GF and CR groups. In adult fish at 3 mpf, the anterior-middle and middle-posterior intestines were observed to be fully developed with numerous and curly villi in both CR and GF fish. However, the villi height and width and muscularis thickness of GF fish were significantly lower than CR fish, except for the villi width of the anterior-middle intestines. All these results showed the significant defect during the growth, indicating that absence of the gut microbiota in fish may contribute to a deficiency in intestinal system development and weak digestive or absorption ability.

Furthermore, in fish, the goblet cells which distributed among the epithelial cells of the intestinal and respiratory tracts, can secrete mucus, a viscous fluid composed primarily of highly glycosylated proteins called mucins (44–46). Mucus layers serve important functions in intestinal metabolic and inflammation processes, including in protection against shear stress and chemical damage, and trapping and elimination of particulate microorganisms (47–49). In this study, it observed that the distribution of mucous layers in CR marine medaka showed more purple-red neutral mucins in the anterior intestine, and more acid and mixed mucins in the middle to posterior intestines stained with blue and blue-purple. However, the neutral mucins in the anterior intestines and the mixed mucins in the middle to posterior intestines seemed to be the dominant types in GF fish. Moreover, the goblet cells were increased along with the life stages from juvenile to adult, but were still significantly less abundant in the anterior to middle intestines and middle to posterior intestines in GF fish than in CR fish. The histopathological indexes of fish intestines strongly demonstrated that the existence of a normal gut microbiota in fish critically influenced the intestinal development, goblet cells, and mucin production, and potentially, subsequent digestive functions.

Reproductive stage and sex should be taken into consideration when assessing the risk posed by environmental contaminants to the immune competence of fish (41). Importantly, in this study, the reproductive system was proven to appear developmental differences in gonad tissues between GF and CR marine medaka, wherein the ovary and testis were not completely mature at the juvenile and early-adult stages, especially in female and male GF fish. Among juvenile fish, GF and CR female fish showed mostly the early stages I (oogonia) and II (previtellogenic oocytes) in growing ovary tissues. While at 2 mpf, the almost fully grown testis in size and cell stages with matured sperm indicated the early-adult stage of CR fish. However, the germ cells in GF developing testes were mainly in the spermatogonia (SG) and spermatocytes (primary and secondary, SC) stages. Finally, at 3 mpf, the ovary tissues in GF fish were still immature with the oocytes in I, II, III (vitellogenic oocytes) and IV (postvitellogenic oocytes) stages, with the significantly less oocytes at III stages. The testis tissues of GF fish also showed significantly fewer germ cells in the SG, SC, spermatids (ST) and spermatozoa (SZ) stages. However, the oocytes and germ cells in CR medaka at the adult stage were more mature with cell counts and highly differentiated stages, indicating the reproductive ability to produce high quality eggs or sperm of female and male fish. According to the biology and experimental book, the development and histopathological judgement of the ovary and testis in medaka were important indicators of fish maturity and offspring (5). A previous study also reported that the gut microbiota was closely related to host reproduction and that probiotic regulation positively influenced the physiological performance, including the development and reproduction of zebrafish models (50). Thus, we suspected that the GF fish of both zebrafish and marine medaka gained distinct developmental delay by the absence of a gut microbiota, as well as deficiencies in female and male gonad tissue size and cells stages, which possibly led to imperfect functions and the failure in spawning and fertilization.

John *et al*. reported the generation of GF zebrafish and their application in studying the effects of the microbiota on epithelial renewal and enterocyte morphology, as well as the host transcriptional responses to the microbiota in larval models (8). In this study, the transcriptomic profile revealed a total of 1030 DEGs including up- and down-regulated genes in GF-2-VS-CR-2 group, and 4046 DEGs in GF-3-VS-CR-3 group, 1662 DEGs in GF-3-VS-GF-2 group, and 4530 DEGs in CR-3-VS-CR-2 group. Among these DEGs, the important genes *gsdf*, *igfals*, *dio2*, *tgfb5*, *egfl7*, *cgref1*, *igf1*, *gh1*, *fgfr2*, *egr1*, *pkd2*, and *fgf16* were significantly inhibited in GF fish compared to CR fish at 2 and 3 mpf. The critical signaling pathways were significantly impacted in marine medaka involved in the growth hormone synthesis, secretion and action, thyroid hormone signaling pathway, ovarian steroidogenesis, PI3K-Akt signaling pathways, parathyroid hormone synthesis, secretion and action, the GnRH and TNF signaling pathways. On the other hand, in adult GF and CR fish compared to the corresponding early-adult fish, the key genes of *pdgfrl*, *tgfbr2b*, *tgfb5*, *mmp14b*, *gnrh3 gdf6a*, *ngfra*, *fgf23*, and *ngfb* were significantly up-regulated at 3 mpf. Moreover, these DEGs related to the cell cycle, growth hormone synthesis, secretion and action, pancreatic cancer, TGF-beta signaling pathway, parathyroid hormone synthesis, secretion and action, GnRH and TNF signaling pathways, while the genes *socs2*, *fgf1a*, *gadd45ga*, *socs3a*, *gadd45ba*, *gh1*, and *isl1* were significantly inhibited as depressors in these signaling pathways. The screened key DEGs in GF and CR marine medaka at early-adult and adult were examined first, and differences in the expression profile between the compared groups were demonstrated, which potentially provide insights into gut microbiota effects on fish growth and development.

Moreover, the inner mechanisms were analyzed with the expression type and enriched pathways based on the screened DEGs sets, which could be the “biomarkers” for evaluation of the healthy state of marine medaka fish. Complement and coagulation cascades, ECM-receptor interaction, Focal adhesion, and Protein digestion and absorption, and especially, the Ovarian steroidogenesis, were the major enriched KEGG pathways in GF fish compared to CR fish at 2 and 3 mpf. Furthermore, the critical KEGG pathways of Ovarian steroidogenesis enriched in the GF-2-VS-CR-2 group indicated the inhibition of the *insulin*, *igf1*, *cyp17*, and *cox2* genes in GF-2 fish reproductive system, which consistent with the developmental and histopathological analyses. Interestingly, the expression of DEGs in the intestines and blood in the GF-3-VS-CR-3 group may have contributed to the growth delay in the GF models and indicated the differences between adult fish in terms of protein digestive and absorption ability. As fish age increased, the expression of PI3K-Akt pathway and pancreatic secretion pathway were also significantly changed in the CR-3-VS-CR-2 group, and we hypothesized that intestinal function and host physiological metabolism trended to peak period at adult life stage. Overall, the application and elucidation of the GF fish model from juvenile to adult are significant for understanding bacterial functions or drug screening and disease prevention. Thus, the DEGs as well as the signaling pathways involved in regulating growth, development, reproduction, immune system in marine medaka, will serve as potential indicators in our subsequent studies to deeply explore the relationships between the gut microbiota and host health in depth.

In conclusion, in this study, we revealed for the first time the distribution of the gut microbiota in marine medaka during growth and successfully generated GF fish models from larvae to adults with GF *Artemia* as food, which was reported to be difficult or impossible in GF animal models. The innovative protocol of GF marine medaka from larvae to juvenile and adult fish and the special GF *Artemia* model as sterile food in this work will be important for GF animals’ application in bio-medical fields, which provide novel models for studying the functions of the gut microbiota. Moreover, the delayed development indicated by the basic indexes and weaker behavioral ability showed the negative influences of the absence of the microbiota in GF marine medaka. Importantly, the histopathological analysis presented further evidence of developmental differences in immune organs, intestinal villi, goblet cells, gonad tissues and cell maturation between GF and CR marine medaka at various life stages. Notably, the transcriptomic profile showing the significantly changed genes offered the possibility of elucidating the inner mechanisms or signaling pathways of GF fish models in terms of the gut microbiota and host health. Taken together, these findings provide the primary microbial and gene expression background, novel insights and important protocols for the generation and application of GF animal models from the juvenile to adult life stages in biomedicine and experimental animal research regions.

## Methods and materials

### The Ethic statement, chemicals and preparation methods were presented in the *SI Appendix*

#### Marine medaka maintenance and sampling of the gut microbiota

Wild type marine medaka (*O. melastigma*) were obtained from the South China Sea Institute of Oceanology, Chinese Academy of Sciences, and cultured with the following standard procedure in the laboratory. The marine medaka were maintained in glass tanks with the culture seawater refreshed every 48 h and were fed with newly hatched brine shrimp (*Artemia sp.*) twice daily, with a temperature of 28±2.0℃ and a 14:10 h light/dark cycle. The circulating seawater was prepared with ultra-pure water and activated salt, with a final salinity of 35‰, and then was UV-sterilized before use. The fertilized eggs of marine medaka were collected daily from the tank bottom and could be cultured as GF model fish for 3 months to be adult, as well as in conventionally raised (CR) models for 2.5 month to be adult fish.

To illustrate the microbial structure during marine medaka development, the fish in this study were maintained from embryos to adult and were sampled at different life stages. First, the marine medaka embryos with normal development were selected at 6-8 hours post-fertilization (hpf) and randomly placed into three replicate glass tanks. With the standard procedure, the fish embryos were hatched to larvae approximately at 14 days post-fertilization (dpf), and finally grown to adults by providing refreshed seawater daily and enough food (micro-particle food and shelled *Artemia* eggs for larvae, *Artemia* for later larvae to juveniles and to adults). During growth, the intestinal samples of female and male marine medaka were collected at 1, 3, and 5 months post-fertilization (mpf) from each replicate tank. Ther were totally 6 groups including the female groups (F1, F3, F5) and male groups (M1, M3, M5), wherein sequencing and analysis were performed to distinguish the differences induced by genders and developmental stages.

At each time point, the fish were washed with ultrapure water and anesthetized in 100 mg/L of MS-222, and then dissected for intestine sampling after the body weight and body length recorded. Next, the samples were immediately frozen in liquid nitrogen and stored at -80°C for subsequent analysis of microbial community.

#### 16S rRNA sequencing of the gut microbiota in marine medaka

In this study, totally 18 samples from 6 groups (F1, F3, F5, M1, M3, M5) with three entire fish intestines mixed as one biological replicate sample and three replicate samples per group were collected and measured. Briefly, the microbiota of male and female fish was detected *via* 16S rRNA high-throughput sequencing, and analyses were performed referred to our previous reports (46, 51). After total DNA extraction and quality measurement, the microbial community composition of marine medaka was detected by the Majorbio Cloud platform (www.majior.com) using the adapter primers 338F (ACTCCTACGGGAGGCAGCA) and 806R (GGACTACHVGGGTWTCTAAT) for the bacterial 16S rRNA V3-V4 regions (methods shown in **Text S1** and detailed software shown in **Table S1**). The analysis of microbiota was described in **Text S2**.

#### Isolation and identification of intestinal bacteria in adult marine medaka

To explore the intestinal bacterial characteristics and functions, several intestinal bacterial strains were isolated and identified, according to the approaches used in previous studies on fish (10, 14, 51). First, isolation included the following steps, which should be performed in the clean bench:

(A). Clean the adult marine medaka fish in plates with sterile water, wash 3 times×2 min

(B). Anaesthetize the fish with 100 mg/L MS-222 and then wash 3 times×2 min;

(C). Use the 75% alcohol to treat the fish body surface, and then wash 3 times×2 min;

(D). Move the fish to sterilized gauze, dissect the intact intestinal tissue using sterilized tweezers and scissors, and cut the tissue to three fragments as anterior-intestine, middle-intestine, and posterior-intestine;

(E). Extrude the intestinal contents with TSB medium in 1.5 mL EP tubes;

(F). Incubate the intestinal supernatant applying TSA, blood plate, TSB, BHI medium in aerobic and anaerobic conditions;

(G). Translate the bacteria to generations until signal strain, record the morphological characteristics of each bacterial colony, including size, color, surface and margin;

(H). Mix the bacteria within 30% glycerol and store at -80°C for subsequent experiments.

Then, identification of the intestinal bacteria was performed as the following steps:

(A). The genomic DNA of the intestinal bacteria was extracted according to the instructions for the kit;

(B). Then, PCR of the bacterial genome with the 27 F and 1492 R primers was performed;

(C). Sequencing of the products for each bacterial strain was detected and analyzed by the Chromos, DNAman, and Vector software;

(D). Then, NCBI-BLAST by using the Nucleotide database and 16S ribosomal RNA (Bacteria and Archaea) database;

(E). Select the best matched strains and consider the growth characters, to identify the isolated intestinal strains at the genus level by comprehensive analysis.

#### Generation and detection of the GF *Artemia* model by innovative incubation within 24 h

In this study, the germ-free (GF) *Artemia* model was generated with innovative treatment and hatching approaches, and the detection of sterile conditions. And the details were shown as the following steps in the biosafety cabinet:

(A). To prepare the materials: 5.0 g iodized salt was added to a blue cap bottle containing 200 mL ultrapure water to prepare 2.5% hatching brine, and the experimental materials such as hatching brine, ultra-pure water, ultra-dense strainer mesh, 100 mL blue-cap containers, 50 mL EP tubes, TSB and BHI medium, and 100 mL beaker were sterilized in an autoclave.

(B). 0.5 g normal shelled brine shrimp eggs (*Artemia sp*. cysts) were treated for 5 min with freshly prepared 2.4 g/L NaClO solution.

(C). After bleaching in step (B), the eggs were filtered through a short sterilized ultra-dense strainer mesh (about 17.5 cm in length, 6.0 cm in radius and 170 mesh in aperture), and rinsed 3 times (1-2 min per time) with sterilized ultrapure water.

(D). Approximately 0.5 g *Artemia* eggs after cleaning in step (C) were transferred to 200 mL sterilized saline prepared in step (A), mixed and separated according to the following methods for aseptic hatching and aseptic detection:

(a). The mixture of 50 mL sterilized *Artemia* eggs was incubated in a blue-cap glass bottle (100 mL), sealed with sealing film and shaken at 30°C for 24 h at 150 rpm in light conditions.

(b). 1 mL sterilized *Artemia* egg mixture was coated on TSA plates, sealed and placed in thee 30°C incubator for 24 h.

(c). 1 mL mixed solution of sterilized *Artemia* eggs was collected in an aerobic glass tube with 5 mL TSB and BHI medium, sealed and placed in 30°C shaker incubator, and cultured at 150 rpm for 24 h under light conditions.

(E). The hatched *Artemia* nauplii (approximately 30 mL) were taken out from the suspension of the middle and lower layers of the *Artemia* nauplii with a disposable sterile straw (taking care to avoid floating eggs and empty shells of the upper layer). And filtered with a sterile strainer, and then washed 3 times in a sterilizing beaker with sterile ultra-pure water, about 30 s-1 min each time. After washed and filtered, the sterile *Artemia* suspension was prepared with about 4 mL sterilized ultra-pure water, and the upper and middle layers of the active aseptic *Artemia* suspension (taking care to avoid unmoved or non-swimming *Artemia* at the bottom and several floating eggs/shells) were extracted again and moved into new aseptic EP tubes. Finally, about 4 mL of sterilized ultrapure water was added to prepare GF *Artemia* suspension for feeding and aseptic detection.

(F). The TSA plate, blood Plate, TSA double plate, TSB and BHI medium under aerobic and anaerobic conditions were applied to test samples of *Artemia* eggs after rinsing treatment and *Artemia* nauplii after hatching in sterile brine water. Moreover, the collected samples included untreated eggs, washed eggs after rinsing treatment, *Artemia* nauplii after incubation and washing, untreated and commonly incubated *Artemia* nauplii, and the sterilized ultrapure water as control. Additionally, the bacteria were incubated at 30°C and 150 rpm with light and shaking, and the plates were incubated in 30°C incubator without light. The results from the plates and liquid tubes were observed at 24 h, 48 h, 72 h, 96 h, and 120 h, and the observation and recording were extended until 168 h.

(G). The collected samples from all the groups were also detected for sterility by using PCR and 16S rRNA primers (27F and 1492R, 63F and 1387R) for bacteria.

(H). According to the results from the coated plate, incubated medium, PCR and other tests, successfully generated GF *Artemia* could be used to feed the hatched larvae of GF zebrafish or GF marine medaka beginning from the mouth-opening period. That is, GF larvae to juvenile fish were fed with 1-2 GF *Artemia* per fish each day at first, and sterility was tested. The next day, the waste water was changed to remove the residue and fed with freshly prepared GF *Artemia,* which could be increased to 3-5 GF *Artemia* or more per fish during growth until the adult stage.

In addition, the methods of TSA, TSB, and BHI at aerobic condition used in the sterile detection also successfully generated GF *Artemia* with hatching at 24 h.

#### Optimized protocol for generation GF marine medaka from larval to adult stages

In this study, the innovative protocol was optimized to generate and detect GF marine medaka from embryos to larvae to adults (the whole progress was shown in **Fig. 6A**), which included the following steps that were conducted within the biosafety cabinet:

• Preparation of the adult fish to spawn and collection of embryos.

(A). Adult mature marine medaka at sexual maturity (8-10 pairs) were selected after feeding in the afternoon and placed in a clean crystallizing dish with the GMM solution.

(B). The fertilized embryos that naturally fell off from the female’s cloacal pore or at the bottom of the tank were collected in the next morning with a straw, and the fused fertilized eggs immediately separated and wash in AB-MMEM solution for three times. Then, the embryos were moved to clean cell/tissue culture dishes with AB-GMM solution and culture at 28∼30°C for 6-8 hours.

• Sterile treatment to obtain GF embryos and cultured to GF larvae with detections.

(C). After incubation in AB-GMM, healthy embryos with normal development were selected and washed with AB-GMM for 3 times in 10 min, and then, gently immersed in 0.2 g/L PVP-I solution for 1 min. (Note: this should not be performed for more than 1.5 min, otherwise the death rate of the embryos will increase).

(D). The embryos above were rinsed with GMM solution for 3 times (about 10 minutes), and then bleached with 0.04 g/L sodium hypochlorite solution (diluted with GMM solution) for 10 min (Note: the suitable rang is 10-15 min). The tubes were gently rocked to ensure that no bubbles and the embryos were suspended in the bleach solution.

(E). After step (D), the embryos were washed with GMM for 3 times (about 10 minutes), and then transferred to 24-well plates for culture, and 1 embryo per well was cultured in 2 mL GMM solution and at 28∼30°C with a photoperiod 14: 10 h (light/dark) in the biochemical incubator.

(F). The GF embryos were observed, and the failed developed ones were moved when the medium solution was changed daily. Waste water and dead eggs were collected as samples for detections. Moreover, the GMM and plates were completely changed at 7 dpf with the same volume added in the new plates.

• From the larvae to juvenile and adult fish, GF marine medaka were treated and fed GF *Artemia* as food.

(G). After the GF fertilized eggs hatched at approximately 12-15 dpf, the GF larvae were moved to 12-well plates with 2 larvae per well and cultured in 4 mL GMM solution, which was completely changed every day.

(H). The GF larvae were fed sterile micro-particle fish food once a day for 3 days. 0.1 mL of sterile micro-particle food solution or 20∼30 sterile shelled *Artemia* eggs per 3 mL of GMM solution.

(I). The GF *Artemia* were prepared daily and fed to the GF marine medaka fish from larvae to juvenile and adults.

(J). The CR marine medaka were cultured at the same time as the control group under normal conditions and with incubated *Artemia* as food twice daily.

(K). The juvenile fish at 1 mpf were moved to sterile glass bottle containers that were easily sterilized by autoclaving, and increasing volume (30∼150 mL) GMM solution was added during fish growth until adult ones. The sterile conditions of GF models were detected every 3∼4 days.

- The developmental indexes of marine medaka, including the hatching rate, survival state, heart-beat rate, body length, and weight were recorded.
- The survival rate= the observed survival fish number/ selected embryo number in step (F) ×100%.
- The GF fish rate= GF fish number/ survival fish number ×100%.
- The fish samples in the GF and CR groups were collected at 1, 2, and 3 mpf for histopathological analysis of gut, liver, kidney tissues and so on, while the other samples were immediately frozen in liquid nitrogen and stored at -80°C for subsequent analysis.

#### Identification of GF and CR marine medaka models at the whole life stages

• Samples collection was performed as follows:

1). Waste water from the GF and CR fish models;

2). Food of larvae and adult fish, including sterile egg yolk solution and GF *Artemia*;

3). Fish medium and the agents used in the fish embryo treatment as the sterile control;

4). Waste medium included fragments of the embryo membrane at the hatching life stage, and food residue or feces during the feeding and eating periods from larvae to adult fish.

5). Dead fish bodies in the samples, to identify the model’s status: germ-free or bacterial contaminated;

• The key sampling points were as follows:

1). Sampling after GF models were prepared, to check waste water when the embryos were treated;

2). Samples collected each day form GF embryos to hatching larvae;

3). Sampling with the residue when feeding was performed during larvae to juvenile life stages;

4). Sampling every three days from juvenile to adult life stages;

5). Sampling when the fish culture plates or containers were newly changed.

• Detection approaches for collected samples from the GF and CR models at different key points during growth were as follows:

1). TSA plate: TSA plates were prepared with 400 mL medium containing 6 g tryptone, 2 g soy peptone, 2 g NaCl, 6 g agar powder. Then, 50-100 µL samples were added and the plates were coated. Finally, the plates were cultured in the 30°C incubator.

2). Blood plate: Sterile defibrinated sheep blood was added into TSA to prepare 5% blood plates. Then, 50-100 µL samples were added and coated on the blood TSA plates. Finally, the plates were cultured in the 30°C incubator.

3). TSB Aerobic: TSB medium was prepared, wherein 400 mL medium included 6 g tryptone, 2 g soy peptone, 2 g NaCl. Then, 100-200 µL samples were added to the TSB tubes and cultured in the shaking incubator with 30°C and 180 rpm.

4). TSB Anaerobic: TSB medium was prepared and divided to the anaerobic tubes, and then filled with high-purity nitrogen to exhaust air before autoclave sterilization. Then, 100-200 µL samples were added to TSB tubes and cultured in the 30°C incubator.

5). BHI Aerobic: BHI medium was prepared, wherein 100 mL medium included 3.7 g BHI powder. Then, 100-200 µL samples were added to BHI tubes and cultured in the shaking incubator at 30°C and 180 rpm.

6). BHI Anaerobic: BHI medium was prepared and divided to anaerobic tubes, and filled with high-purity nitrogen to exhaust air before autoclave sterilization. Then, 100-200 µL samples were added to TSB tubes and cultured in the 30°C incubator.

7). Double TSA plate: The second layer was added after 50-100 µL samples were coated on the TSA plates. Then, the plates were cultured in the 30°C incubator.

Note: The experimental materials above should be sterile after autoclave treatment, and then prepare the plates; all the plates and liquid medium should be stored at 4°C for subsequent use.

8). PCR: To examine the waste water samples with totally 20 µL, including 0.1 µL Taq DNA polymerase, 2.1 µL 10×Buffer (Mg^2+^), 1.6 µL dNTP, 1.0 µL bacterial DNA, 1.0 µL primers 27F and 1.0 µL primers 1492R, 13.2 µL DNA/RNA free H_2_O. PCR progress was performed as follows: 95°C 4 min; 35 cycles of 94°C 1 min, 55°C 1 min, 72°C 1 min; 72°C 10 min. Finally, the PCR products were detected by 1% agarose gel electrophoresis, which indicated the bacteria or sterility status of the CR and GF fish models and *Artemia* samples.

Note: the key period of these detections using plates and medium was observed at 24 hours until 7 days. During the observation time, no appearance of bacterial colony or liquid medium turbidity indicated that the samples were germ-free, and that the corresponding GF models were successfully prepared.

**Developmental indexes, behaviors, histopathological analysis, transcription profiles of GF and CR marine medaka at different life stages**, and the Statistical analysis were all presented in the ***SI Appendix***.

## Acknowledgments

We thank the supports from Chongqing Medical University Talent Project (No. R4014 to D.S.P., and R4020 to P.P.J.), National Natural Science Foundation of China (NSFC, No.32200386 to P.P.J), Three Hundred Leading Talents in Scientific and Technological Innovation Program of Chongqing (No. CSTCCXLJRC201714 to D.S.P.), Youth Innovation Program of Chongqing Institute of CAS (No.Y83A160 to P.P.J), the CAS Team Project of the Belt and Road (to D.S.P.), Chongqing Key Program of Basic Research and Advanced Exploration Project (No. cstc2019jcyj-zdxmX0035 to D.S.P.), Program of China-Sri Lanka Joint Center for Water Technology Research and Demonstration by Chinese Academy of Sciences (CAS)/China-Sri Lanka Joint Center for Education and Research by CAS, and International Partnership Program of CAS (No. 121311kysb20190071).

## Author contributions

D.S.P. designed research; Y.F.Y., and P.P.J. performed research; D.S.P., W.G.L., J.J.D., and Y.W. contributed new reagents/analytic tools; P.P.J. analyzed data; D.S.P., and P.P.J. wrote the paper, improved and submitted the manuscript.

## Declaration of Competing Interest

The authors declare no conflict of interest.

## Notes

### Competing Interest Statement

Chinese Patent: CN114287366A

