## Supplementary Information for "Breaking the mold: The first report on germ-free adult marine medaka (*Oryzias melastigma*) models"

*Corresponding author.

### Methods and materials

#### Ethic statement

The fish experiments were performed according to the "Guide for the Care and Use of Laboratory Animals" (Eighth Edition, 2011. ILARCLS, National Research Council, Washington, D.C.). The animal protocol was approved by the Animal Care and Use Committee of Chongqing, and the Institutional Animal Care and Use Committee of Chongqing Medical University, Chongqing, China, which was referred to the Administration of Experimental Animals issued by the Ministry of Science and Technology, and standards for experimental animals issued by the State Bureau of Quality and Technical Supervision (Approval ID: GB14922-2001 to GBT14927-2001).

#### Chemicals and preparation

In this study, the ocean activated salt was applied in preparation of the seawater and gnotobiotic marine medaka medium (GMM) with the reverse osmosis pure water. The amphotericin B (CAS: 1397-89-3, Cat#A8252), ampicillin sodium salt (CAS: 69-52-3, Cat#A8180), kanamycin sulfate (CAS: 25389-94-0, Cat#K8020), and penicillin-streptomycin mixed solution (100×, Cat#P1400) purchased from Solarbio Science &Technology Co., Ltd were used to prepare the antibiotic-containing GMM (AB-GMM) with the 0.22 μm membrane filter unit (Millipore, No. SLGP033RB). The sodium hypochlorite solution (NaClO) with 6% was purchased from the KESHI company (China, CAS: 7681-52-9). And the poly (vinylpyrrolidone)-iodine complex (PVP-I, CAS: 25655-41-8) was obtained from the Aladdin company. Moreover, the NaCl (CAS: 7647-14-5), soy peptone (Coolaber, Cat#CS10383), tryptone (Oxoid, Cat#LP0042), and agar powder (Solarbio, CAS: 9002-18-0) were used to prepare the tryptic soy agar (TSA) plates and tryptic soy broth (TSB) medium, and the sterile defibrinated sheep blood (Solarbio, Cat#TX0030) was added into TSA to prepare the 5% blood plates. The brain heat infusion (BHI, Cat#B8130) medium, glycerol (>99.0%, CAS: 56-81-5, Cat#G8190), and the bacterial genomic DNA extraction kit (Cat#D1600) were purchased form the Solarbio company. The Taq$^{TM}$ (TaKaRa, Cat#R001A) kits and the bacterial genomic DNA extraction kit (Solarbio, Cat#D1600) were used for the normal PCR assay to identify the bacterial strain and GF model samples.

The brine shrimp eggs (*Artemia* sp. cysts) and the shelled shrimp eggs were acquired from Shangjia Aquariums and stored at 4°C, and the micro-particle food (ZeiGLeR, USA, <50 microns) contained algae, pigments, and Vitality Pak formulated with highly unsaturated fatty acids (HUFA's Vpak) for fish larvae early feeding were purchased from the Shanghai Haisheng Biological Equipment Co., Ltd. In addition, the 3-Aminobenzoic acid ethyl ester or methane-sulfonate salt (MS-222, 98%) was obtained from Aladdin (Los Angeles, California, USA). The neutral paraformaldehyde fixative (Servicebio, Cat#G1101) was applied in animal and tissues histopathological analysis. All other chemical reagents used in this study were of analytical grade.

The GMM solution was prepared with the 1 L water and 35 g sea salt to the final 35 mg/mL concentration, then sterilized under the autoclave. The stock solution of amphotericin B with 500 ng/mL, and kanamycin with 10 mg/mL, ampicillin with 100 μg/mL were solvent in ultra-pure water, and the penicillin-streptomycin mixed solution (100×) were comprise of 10000 U/mL penicillin and 10000 μg/mL streptomycin. In addition, the AB-GMM solution were prepared with 49.55 mL GMM solution and 100 μL amphotericin B stock solution, 50 μL kanamycin solution, 250 μL ampicillin solution, and 50 μL penicillin-streptomycin solution and then mixed and filtered with 0.22 μm filter into 50 mL EP tube, and then stored at -20°C. The final content of amphotericin B in the AB-GMM solution was 500 ng/mL, and the content of kanamycin sulfate was 10 μg/mL, the content of ampicillin was 100 μg/mL, the content of penicillin and streptomycin were 10 U/mL and 10 μg/mL, and the content of sea salt was 35 mg/mL.

**The developmental indexes and behaviors of GF and CR marine medaka**

To investigate the developmental differences, marine medaka were collected at 1, 2, and 3 months-post-fertilization (mpf) from the GF and CR groups. There were totally six groups including the GF groups (GF-1, GF-2, GF-3) and CR groups (CR-1, CR-2,

CR-3), which were performed to analyze the growth and developmental differences. At the key points, the fish were washed with ultrapure water and anesthetized in 100 mg/L of MS-222, and the developmental indexes including survival rate, hatching rate, heart-beats, body length and weight were recorded and calculated from each group. Then, the samples were collected and fixed for histopathology, and immediately frozen in liquid nitrogen and stored at -80°C for the subsequent transcriptomic analysis.

In addition, the behaviors of CR and GF marine medaka at 1 mpf were examined and analyzed by the Danio Vision™ instrument with 20 fish randomly selected from each group. High-speed infrared camera was used to record the movement track of fish, and to quantitatively analyze the obtained behavioral data and the active ability. Before test, the fish were placed (with one fish per well) in 12-well plates containing GMM solution and adjusted for 10 min with the water temperature 28 ± 2.0°C. All fish in the plates were collected the swimming data for 10 min at rate of 25 times/sec under the maintained quiet environment during the experiment. Finally, the collected Video were analyzed using computer tracking software EthoVision XT® (Noldus, Netherlands), and the behavior data as well as the track heat-map can be analyzed and exported from the software.

**The histopathological analysis of GF and CR marine medaka during growth**

To compare the developmental differences between the GF and CR marine medaka, the sampling of fish from larvae to adult was carried out at various life stages. Briefly, the marine medaka were washed with ultra-pure water and anesthetized in 100 mg/L of MS-222, and then were fixed in the formaldehyde solution, then dehydrated in ethanol and embedded in paraffin to prepare the slices in the laboratory of Chongqing Yike Tianya Biotechnology Co., Ltd. The histopathological analysis of tissue sections stained with hematoxylin and eosin (H&E) were subsequently carried out to evaluate further the development. The magnified pictures with marked scales were stored and scanned by K-Viewer software (2.7.2.0 version). Additionally, the H&E section images with morphometric parameters at 10× magnification were taken by digital microscope

(Nikon, SMZ18) with NIS-Elements imaging software (version 4.30).

**The transcriptomic profile of GF and CR marine medaka at different life stages**

To explore the changes of transcriptomic profile between GF and CR marine medaka, the early-adult at 2 mpf and adult fish at 3 mpf life stage were washed with ultra-pure water and anesthetized in 100 mg/L of MS-222, and then sampled for the subsequent analysis. Then, the total RNA of the three replicate samples in each group was extracted by using TRIzol® Reagent according to the manufacturer's instructions (Invitrogen) and genomic DNA was removed using DNase I (Takara). The quality of RNA samples was determined by 2100 Bioanalyser (Agilent) and quantified using the ND-2000 (NanoDrop Technologies). The high-quality RNA samples (OD260/280=1.8~2.2, OD260/230 ≥ 2.0, RIN ≥ 6.5, 28S:18S ≥ 1.0, >1 μg) were used to construct the sequencing library. Then, RNA-seq transcriptome library was prepared following the TruSeq TM RNA sample preparation Kit from Illumina (San Diego, CA). Shortly, messenger RNA was isolated according to poly-A selection method by oligo (dT) beads and then fragmented by fragmentation buffer firstly. Secondly, the double-stranded cDNA was synthesized using a SuperScript double-stranded cDNA synthesis kit (Invitrogen) with random hexamer primers (Illumina). Then the synthesized cDNA was subjected to end-repair, phosphorylation and 'A' base addition according to Illumina's library construction protocol. Libraries were size selected for cDNA target fragments of 300 bp on 2% Low Range Ultra Agarose followed by PCR amplified using Phusion DNA polymerase (NEB) for 15 PCR cycles. After quantified by TBS380, the paired-end RNA-seq sequencing library was sequenced with the Illumina HiSeq xten/NovaSeq 6000 sequencer.

The raw paired-end reads were trimmed and quality controlled by SeqPrep (https://github.com/jstjohn/SeqPrep) and Sickle (https://github.com/najoshi/sickle) with default parameters. Then clean reads were separately aligned to reference genome with orientation mode using HISAT2 (http://ccb.jhu.edu/software/hisat2/index.shtml) software (1). The mapped reads of each sample were assembled by StringTie (https://ccb.jhu.edu/software/stringtie/index.shtml?t=example) in a reference-based approach (2). To identify differential expression genes (DEGs) between two different samples, the expression level of each transcript was calculated according to the transcripts per million reads (TPM) method. RSEM (http://deweylab.biostat.wisc.edu/rsem/) was used to quantify gene abundances. Essentially, differential expression analysis was performed using the DESeq2 and EdgeR with Q value$\leqslant$0.05 and |log2FC|>1 considered to be significantly differentially expressed genes. In addition, functional-enrichment analyses including GO and KEGG were performed to identify which DEGs were significantly enriched in GO terms and metabolic pathways at Bonferroni-corrected $p$-value$\leqslant$0.05 compared with the whole-transcriptome background. GO functional enrichment and KEGG pathway analysis were carried out by Goatools (https://github.com/tanghaibao/Goatools) and KOBAS (http://kobas.cbi.pku.edu.cn/home.do) tools.

**Statistical analysis**

The statistical analysis was performed using the Kolmogorov-Smirnov test and Leven's test to define the normality of data and the homogeneity of variance. The differences between the variables were calculated by t-test and one-way analysis of variance (ANOVA), followed by Dunnett's test using SPSS 20.0 software (SPSS, Chicago, IL, USA). The figures were represented by using the GraphPad Prism 5, and the correlation of the transcription and gut microbial profile was deeply analyzed using the Spearman's methods. The value of $p<0.05$ was denoted as statistically significant differences between control and treated groups, and all data were presented as the means ± standard error (SEM) of replicates in each group.

**Text S1. The methods of total DNA extraction and PCR amplification, 16S rRNA sequencing of fish intestinal microbiota**

At first, the total genomic DNA of intestinal samples was extracted and purified with FastDNA® SPIN Kit for Soil (Mpbio, USA). The concentration and purity of extracted DNA samples were assessed by a NanoDrop 2000 Spectrophotometer (Thermo Scientific, USA), and the integrity of DNA was checked by 1% agarose gel electrophoresis. The concentration of extracted DNA was assessed by a NanoDrop ND-2000 (Thermo Fisher Scientific, USA). Amplicons for sequencing were amplified as previously described (3). Briefly, extracted DNA was first amplified with high-fidelity Taq polymerase (Invitrogen, USA). Then equal quantities of three PCR reactions per sample were pooled, purified with the QIAquick PCR Purification Kit (Qiagen, Valencia, CA). The PCR amplification of 16S rRNA gene was performed as follows: initial denaturation at 95°C for 3 min, followed by 27 cycles of denaturing at 95°C for 30 s, annealing at 55°C for 30 s and extension at 72°C for 45 s, and single extension at 72°C for 10 min, and end at 4°C by an ABI GeneAmp® 9700 PCR thermocycler (ABI, CA, USA). Finally, the purified amplicons were pooled in equimolar and paired-end sequenced on an Illumina MiSeq PE300 platform (Illumina, San Diego, USA) according to the standard protocols by Majorbio Bio-Pharm Technology Co. Ltd. (Shanghai, China). Raw reads were sorted and can be analyzed using the platform of the company Majorbio (http://www.i-sanger.com/).

**Text S2. The analysis of fish intestinal microbiota**

The raw 16S rRNA sequencing reads were demultiplexed and quality-filtered by Fastp (0.19.6) and merged by (Flash version 1.2.11) with the following criteria: the maximum number of errors in the barcode was 0 and the number of mismatches in the primer was 2. Reads with more than 10% of bases with a quality score of $Q<20$, and ambiguous or unassigned characters and adapter contamination were removed. USEARCH (version 7.0) was used to classify the operational taxonomic units (OTUs) based on sequence similarity by cluster cut-off value of 97%. The taxonomy of each OTU representative sequence was analyzed by RDP Classifier (version 2.2) against the 16S rRNA database (Release 138) using confidence threshold of 0.7. The bacterial alpha diversity indexes were calculated by Mothur (version 1.30.2), and the alpha rarefaction curves showed that the reads number of each sample was sufficient for subsequent analysis.

The fish intestinal community at different life stages was analyzed based on the OTUs abundance in each sample with the coverage (the Good's coverage) and different alpha diversity indexes. Among them, the Shannon and Simpson reflected the community diversity, while the abundance-based coverage estimator (Ace) and Chao1 indicated the community richness. Moreover, the composition of phylum and genus with the community heatmap, as well as the principal component analysis (PCA) analysis were explored in male and female fish at different life stages. Additionally, the Kruskal-Wallis H test, and Student t test were applied to deeply explore the major changes of intestinal bacterial between different groups. Moreover, the Spearman correlation analysis was carried out to explore the relationship between developmental factors and microbial richness and functions. Finally, the bacterial functions and potential phenotype in marine medaka gut were predicted by PICRUSt software with EggNOG (evolutionary genealogy of genes: Non-supervised Orthologous Groups) database and KEGG (Kyoto Encyclopedia of Genes and Genomes) database, and BugBase (https://bugbase.cs.umn.edu/index.html) tool.

**Table S1. The analysis software or database of 16s rRNA sequencing used in the present study.**

| Analysis software/database | Version | Use | Links |
|---|---|---|---|
| Flash | 1.2.11 | Pair-end sequencing reads merged | https://ccb.jhu.edu/software/FLASH/index.shtml |
| Qiime | 1.9.1 | Taxonomies, Beta diversity, distance calculations | http://qiime.org/install/index.html |
| Uparse | 7.0.1090 | OTU Clustering | http://www.drive5.com/uparse/ |
| RDP Classifier | 2.11 | Sequence classification annotation | https://sourceforge.net/projects/rdp-classifier/ |
| Usearch | 7.0 | OTU Statistics | http://www.drive5.com/usearch/ |
| Mothur | 1.30.2 | alpha diversity | https://www.mothur.org/wiki/Download_mothur |
| PICRUSt | 1.1.0 | KEGG、COG、Pfam Functional prediction | http://picrust.github.io/picrust/ |
| Mega | 7.0 | Evolutionary Tree Analysis | https://www.megasoftware.net/ |
| SILVA | 138 | rRNA database | https://www.arb-silva.de/ |
| RDP | 11.5 | rRNA database | http://rdp.cme.msu.edu/ |
| GreenGenes | 135 | rRNA database | http://greengenes.secondgenome.com/ |
| FunGene | 9.6 | Functional Gene Database | http://www.fungene-db.fr/ |
| HPB | --- | human pathogens database | https://www.cerl.org/resources/hpb/content |
| Tax4fun | 0.3.1 | Tax4Fun Functional prediction | http://tax4fun.gobics.de/ |
| MAFFT | 7.2 | Multiple sequence alignment | https://mafft.cbrc.jp/alignment/software/ |
| IQ-TREE | 1.6.8 | Making evolutionary trees | http://www.iqtree.org/ |
| Fastp | 0.19.6 | Quality Control | https://github.com/OpenGene/fastp |
| PICRUSt2 | 2.2.0 | KEGG orthologys (KO), EC, COG, MetaCyc metabolic pathways | https://github.com/picrust/picrust2/ |

**Table S2. The diversity indexes of CR marine medaka at 1, 3, and 5 months life stages.***

| Groups | Sobs | Shannon | Simpson | Ace | Chao | Coverage |
|---|---|---|---|---|---|---|
| F1 | 656.333 | 3.459 | 0.135 | 694.945 | 705.190 | 0.998 |
| F3 | 323.667 | 2.397 | 0.247 | 416.003 | 393.282 | 0.997 |
| F5 | 298.000 | 2.124 | 0.315 | 431.554 | 401.127 | 0.997 |
| M1 | 395.667 | 3.108 | 0.183 | 446.210 | 447.361 | 0.998 |
| M3 | 335.667 | 2.849 | 0.133 | 449.933 | 431.323 | 0.997 |
| M5 | 275.667 | 2.817 | 0.125 | 423.836 | 368.071 | 0.997 |

*The indexes were represented with the average value of three replicate samples from each group.

**Table S3. The significantly changed genera of gut microbiota in marine medaka at different life stages.**

| Group 1: F1.vs.F3 | | | | | | |
|---|---|---|---|---|---|---|
| **No.** | **Name (genus level)** | **F1-Mean(%)** | **F3-Mean(%)** | **Significance** | **Changes** | **P-value** |
| 1. | *Ruegeria* | 2.098 | 46.02 | ** | Up | 0.0052 |
| 2. | *Haloferula* | 0.266 | 5.985 | * | Up | 0.0279 |
| 3. | *Nautella* | 0.036 | 3.460 | *** | Up | 0.0006 |
| 4. | *Iamia* | 0.277 | 1.413 | * | Up | 0.0187 |
| 5. | *Paracoccus* | 0.253 | 0.482 | * | Up | 0.0447 |
| 6. | *Marimicrobium* | 0.040 | 0.618 | * | Up | 0.0101 |
| 7. | *Dermacoccus* | 0.015 | 0.172 | *** | Up | 0.0003 |
| 8. | *Kangiella* | 0.003 | 0.063 | * | Up | 0.0275 |
| 9. | *unclassified_p_Proteobacteria* | 0.044 | 0 | ** | Down | 0.0067 |
| 10. | *Rothia* | 0.034 | 0.009 | ** | Down | 0.0053 |
| 11. | *Lawsonella* | 0.019 | 0.002 | * | Down | 0.0106 |
| 12. | *norank_p_WS6* | 0.021 | 0 | * | Down | 0.0442 |
| 13. | *[Eubacterium]_hallii_group* | 0.017 | 0.001 | * | Down | 0.0406 |
| 14. | *Klebsiella* | 0.011 | 0 | ** | Down | 0.0065 |
| 15. | *Aureimonas* | 0 | 0.008 | ** | Up | 0.0022 |
| 16. | *Idiomarina* | 0 | 0.005 | * | Up | 0.0161 |
| **Group 2: F1.vs.F5** | | | | | | |
| **No.** | **Name (genus level)** | **F1-Mean(%)** | **F5-Mean(%)** | **Significance** | **Changes** | **P-value** |
| 1. | *Ruegeria* | 2.098 | 54.25 | *** | Up | 0.0001 |
| 2. | *Iamia* | 0.277 | 1.639 | ** | Up | 0.0086 |
| 3. | *norank_o_PeM15* | 0.038 | 0.527 | * | Up | 0.0423 |
| 4. | *unclassified_p_Proteobacteria* | 0.044 | 0 | ** | Down | 0.0067 |
| 5. | *Rothia* | 0.034 | 0.008 | * | Down | 0.0169 |
| 6. | *norank_o_Thiotrichales* | 0.002 | 0.019 | * | Up | 0.0106 |
| 7. | *Lawsonella* | 0.019 | 0.001 | ** | Down | 0.0048 |
| 8. | *Gelria* | 0.001 | 0.013 | * | Up | 0.0254 |
| 9. | *Klebsiella* | 0.010 | 0.003 | * | Down | 0.0257 |
| 10. | *Parasutterella* | 0.005 | 0 | * | Down | 0.0161 |
| **Group 3: F1.vs.M1** | | | | | | |
| **No.** | **Name (genus level)** | **F1-Mean(%)** | **M1-Mean(%)** | **Significance** | **Changes** | **P-value** |
| 1. | *Lawsonella* | 0.019 | 0.003 | * | Down | 0.0249 |
| 2. | *Moorella* | 0 | 0.012 | * | Up | 0.0380 |
| **Group 4: F3.vs.F5** | | | | | | |
| **No.** | **Name (genus level)** | **F3-Mean(%)** | **F5-Mean(%)** | **Significance** | **Changes** | **P-value** |
| 1. | *Vibrio* | 0.296 | 5.344 | * | Up | 0.0476 |
| 2. | *Filomicrobium* | 4.442 | 0.929 | * | Down | 0.0337 |

| | | | | | | |
|---|---|---|---|---|---|---|
| 3. | *Nautella* | 3.460 | 0.706 | ** | Down | 0.0029 |
| 4. | *Marimicrobium* | 0.618 | 0.126 | * | Down | 0.0252 |
| 5. | *norank_f_OCS116_clade* | 0.262 | 0.078 | * | Down | 0.0193 |
| 6. | *Dermacoccus* | 0.172 | 0.049 | ** | Down | 0.0027 |
| 7. | *Dietzia* | 0.057 | 0.025 | ** | Down | 0.0044 |
| 8. | *Marixanthomonas* | 0.053 | 0.010 | ** | Down | 0.0013 |
| 9. | *norank_f_Chlamydiaceae* | 0.059 | 0 | * | Down | 0.0156 |
| 10. | *norank_o_Thiotrichales* | 0.006 | 0.019 | * | Up | 0.0327 |
| 11. | *Marinococcus* | 0.016 | 0.001 | ** | Down | 0.0044 |
| 12. | *Rheinheimera* | 0.011 | 0.001 | * | Down | 0.0158 |
| 13. | *Vogesella* | 0 | 0.009 | ** | Up | 0.0013 |
| **Group 5: F3.vs.M3** | | | | | | |
| **No.** | **Name (genus level)** | **F3-Mean(%)** | **M3-Mean(%)** | **Significance** | **Changes** | **P-value** |
| 1. | *Roseovarius* | 2.889 | 4.399 | * | Up | 0.0329 |
| 2. | *Dermacoccus* | 0.172 | 0.055 | * | Down | 0.0186 |
| 3. | *Dietzia* | 0.057 | 0.025 | ** | Down | 0.0054 |
| 4. | *norank_o_JG30-KF-CM45* | 0.054 | 0.015 | * | Down | 0.0310 |
| 5. | *Exiguobacterium* | 0.028 | 0.006 | * | Down | 0.0172 |
| 6. | *Marinococcus* | 0.016 | 0.006 | * | Down | 0.0158 |
| 7. | *Fusicatenibacter* | 0 | 0.015 | * | Up | 0.0227 |
| 8. | *Janibacter* | 0.009 | 0.001 | ** | Down | 0.0078 |
| 9. | *Sphingobium* | 0.005 | 0 | * | Down | 0.0161 |
| 10. | *Idiomarina* | 0.005 | 0 | * | Down | 0.0161 |
| **Group 6: F5.vs.M5** | | | | | | |
| **No.** | **Name (genus level)** | **F5-Mean(%)** | **M5-Mean(%)** | **Significance** | **Changes** | **P-value** |
| 1. | *Ruegeria* | 54.25 | 23.52 | * | Down | 0.0155 |
| 2. | *Filomicrobium* | 0.929 | 0.252 | * | Down | 0.0466 |
| 3. | *norank_o_Thiotrichales* | 0.019 | 0.005 | * | Down | 0.0255 |
| 4. | *Vogesella* | 0.009 | 0 | ** | Down | 0.0013 |
| **Group 7: M1.vs.M3** | | | | | | |
| **No.** | **Name (genus level)** | **M1-Mean(%)** | **M3-Mean(%)** | **Significance** | **Changes** | **P-value** |
| 1. | *Ruegeria* | 1.853 | 29.29 | ** | Up | 0.0061 |
| 2. | *Sulfitobacter* | 1.291 | 5.986 | * | Up | 0.0327 |
| 3. | *unclassified_f_Aurantimonadaceae* | 0.899 | 3.800 | ** | Up | 0.0018 |
| 4. | *Nautella* | 0.028 | 4.217 | * | Up | 0.0371 |
| 5. | *Haloferula* | 0.055 | 2.745 | * | Up | 0.0293 |
| 6. | *Marimicrobium* | 0.002 | 1.064 | * | Up | 0.0321 |
| 7. | *Iamia* | 0.086 | 1.064 | * | Up | 0.0189 |
| 8. | *unclassified_o_Legionellales* | 0.067 | 0.690 | * | Up | 0.0478 |
| 9. | *Dietzia* | 0.009 | 0.025 | * | Up | 0.0352 |
| 10. | *Moorella* | 0.012 | 0 | * | Down | 0.0380 |

| | | | | | | |
|---|---|---|---|---|---|---|
| 11. | *norank_o_Thiotrichales* | 0 | 0.009 | * | Up | 0.0390 |
| **Group 8: M1.vs.M5** | | | | | | |
| **No.** | **Name (genus level)** | **M1-Mean(%)** | **M5-Mean(%)** | **Significance** | **Changes** | **P-value** |
| 1. | *Ruegeria* | 1.853 | 23.52 | * | Up | 0.0355 |
| 2. | *Haloferula* | 0.055 | 13.99 | * | Up | 0.0238 |
| 3. | *Actibacter* | 0.012 | 1.250 | *** | Up | 0.0001 |
| 4. | *Iamia* | 0.086 | 0.912 | * | Up | 0.0485 |
| 5. | *norank_f_Erysipelotrichaceae* | 0.081 | 0.443 | * | Up | 0.0166 |
| 6. | *Rhodococcus* | 0.088 | 0.019 | ** | Down | 0.0061 |
| 7. | *unclassified_o_Cellvibrionales* | 0 | 0.020 | * | Up | 0.0449 |
| 8. | *Moorella* | 0.012 | 0 | * | Down | 0.0380 |
| 9. | *Aeromicrobium* | 0 | 0.01118 | * | Up | 0.0194 |
| 10. | *Exiguobacterium* | 0.002 | 0.008 | * | Up | 0.0241 |
| 11. | *Oceaniovalibus* | 0 | 0.006 | ** | Up | 0.0075 |
| **Group 9: M3.vs.M5** | | | | | | |
| **No.** | **Name (genus level)** | **M3-Mean(%)** | **M5-Mean(%)** | **Significance** | **Changes** | **P-value** |
| 1. | *Haloferula* | 2.745 | 13.99 | * | Up | 0.0486 |
| 2. | *Roseovarius* | 4.399 | 2.090 | * | Down | 0.0186 |
| 3. | *unclassified_f_Aurantimonadaceae* | 3.800 | 1.166 | ** | Down | 0.0012 |
| 4. | *Nautella* | 4.217 | 0.396 | * | Down | 0.0500 |
| 5. | *Marimicrobium* | 1.064 | 0.111 | * | Down | 0.0465 |
| 6. | *Paracoccus* | 0.594 | 0.128 | * | Down | 0.0367 |
| 7. | *norank_f_Erysipelotrichaceae* | 0.112 | 0.443 | * | Up | 0.0223 |
| 8. | *norank_f_OCS116_clade* | 0.172 | 0.031 | ** | Down | 0.0039 |
| 9. | *Pseudonocardia* | 0.113 | 0.039 | * | Down | 0.0198 |
| 10. | *norank_f_Chlamydiaceae* | 0.025 | 0.003 | ** | Down | 0.0020 |
| 11. | *Ilumatobacter* | 0.017 | 0 | * | Down | 0.0307 |
| 12. | *unclassified_f_Intrasporangiaceae* | 0.010 | 0.002 | * | Down | 0.0249 |
| 13. | *Aureispira* | 0.008 | 0 | * | Down | 0.0249 |
| 14. | *Stenotrophomonas* | 0.001 | 0.006 | * | Up | 0.0474 |
| 15. | *Oceaniovalibus* | 0.001 | 0.006 | * | Up | 0.0474 |

Note: The totally nine groups were compared and represented the significantly changed genera between them with the $p$ value of <0.05, 0.01, 0.001.

**Table S4. The KEGG pathway of gut microbiota in marine medaka.**

| Pathway level1 | Pathway level2 | F1 | F3 | F5 | M1 | M3 | M5 |
|---|---|---|---|---|---|---|---|
| Metabolism | Carbohydrate metabolism | 0.12616 | 0.12623 | 0.12623 | 0.12685 | 0.12730 | 0.12969 |
| Metabolism | Lipid metabolism | 0.03806 | 0.03744 | 0.03428 | 0.03695 | 0.03761 | 0.03529 |
| Metabolism | Metabolism of cofactors and vitamins | 0.06816 | 0.07335 | 0.07266 | 0.06686 | 0.07331 | 0.07034 |
| Metabolism | Energy metabolism | 0.06705 | 0.07289 | 0.07287 | 0.06521 | 0.07174 | 0.06822 |
| Metabolism | Nucleotide metabolism | 0.04924 | 0.05391 | 0.05443 | 0.04933 | 0.05414 | 0.05432 |
| Metabolism | Biosynthesis of other secondary metabolites | 0.00891 | 0.00872 | 0.00821 | 0.00842 | 0.00866 | 0.00796 |
| Metabolism | Amino acid metabolism | 0.12526 | 0.13878* | 0.13971* | 0.12031 | 0.13567** | 0.12959 |
| Metabolism | Metabolism of terpenoids and polyketides | 0.03617 | 0.03022 | 0.02347 | 0.03497 | 0.03050 | 0.02527 |
| Metabolism | Xenobiotics biodegradation and metabolism | 0.04412 | 0.04701 | 0.04317 | 0.04022 | 0.04528 | 0.03935 |
| Metabolism | Metabolism of other amino acids | 0.02599 | 0.02849* | 0.02857* | 0.02504 | 0.02815** | 0.02724 |
| Metabolism | Glycan biosynthesis and metabolism | 0.02171 | 0.01883 | 0.01912 | 0.02303 | 0.02044 | 0.02319 |
| Genetic Information Processing | Translation | 0.03897 | 0.03937 | 0.03883 | 0.03951 | 0.04099 | 0.04132 |
| Environmental Information Processing | Membrane transport | 0.11839 | 0.14424 | 0.15374** | 0.11460 | 0.13551 | 0.13835 |
| Environmental Information Processing | Signal transduction | 0.08734 | 0.06369 | 0.06771 | 0.09508 | 0.06619 | 0.07579 |
| Cellular Processes | Cell motility | 0.02725 | 0.01759 | 0.01943 | 0.03057 | 0.01894 | 0.02253 |
| Genetic Information Processing | Folding, sorting and degradation | 0.02165 | 0.02074 | 0.02039 | 0.02221 | 0.02170 | 0.02201 |
| Genetic Information Processing | Transcription | 0.00187 | 0.00171 | 0.00163 | 0.00194 | 0.00179 | 0.00182 |
| Genetic Information Processing | Replication and repair | 0.03767 | 0.03353 | 0.03234 | 0.03910 | 0.03557 | 0.03708 |

| | | | | | | | |
|---|---|---|---|---|---|---|---|
| Organismal Systems | Endocrine system | 0.00392 | 0.00386 | 0.00344 | 0.00371 | 0.00391 | 0.00342 |
| Environmental Information Processing | Signaling molecules and interaction | 0.00002 | 0.00001 | 0.00001 | 0.00002 | 0.00002 | 0.00001 |
| Cellular Processes | Cell growth and death | 0.01606 | 0.01286* | 0.01309* | 0.01700 | 0.01336* | 0.01421 |
| Cellular Processes | Transport and catabolism | 0.00337 | 0.00366 | 0.00350 | 0.00317 | 0.00367 | 0.00358 |
| Organismal Systems | Circulatory system | 0.00014 | 0.00013 | 0.00014 | 0.00014 | 0.00013 | 0.00013 |
| Cellular Processes | Cellular community - eukaryotes | 0.00001 | 0.00001 | 0.00001 | 0.00001 | 0.00001 | 0.00001 |
| Organismal Systems | Immune system | 0.00049 | 0.00060 | 0.00060 | 0.00046 | 0.00064 | 0.00060 |
| Organismal Systems | Environmental adaptation | 0.00241 | 0.00216 | 0.00227 | 0.00255 | 0.00222 | 0.00239 |
| Organismal Systems | Nervous system | 0.00146 | 0.00145 | 0.00141 | 0.00147 | 0.00143 | 0.00137 |
| Organismal Systems | Sensory system | 0.00000 | 0.00000 | 0.00000 | 0.00000 | 0.00000 | 0.00000 |
| Human Diseases | Endocrine and metabolic disease | 0.00048 | 0.00047 | 0.00043 | 0.00048 | 0.00049 | 0.00049 |
| Organismal Systems | Excretory system | 0.00018 | 0.00016 | 0.00015 | 0.00018 | 0.00017 | 0.00018 |
| Organismal Systems | Digestive system | 0.00110 | 0.00054 | 0.00043* | 0.00115 | 0.00083 | 0.00083 |
| Human Diseases | Neurodegenerative disease | 0.00188 | 0.00184 | 0.00188 | 0.00192 | 0.00193 | 0.00197 |
| Human Diseases | Substance dependence | 0.00032 | 0.00031 | 0.00031 | 0.00031 | 0.00030 | 0.00028 |
| Human Diseases | Infectious disease: bacterial | 0.01333 | 0.00739* | 0.00756* | 0.01565 | 0.00895* | 0.01181 |
| Cellular Processes | Cellular community - prokaryotes | 0.00627 | 0.00325 | 0.00359 | 0.00720 | 0.00382 | 0.00501 |
| Human Diseases | Infectious disease: parasitic | 0.00090 | 0.00056* | 0.00047** | 0.00089 | 0.00070 | 0.00065 |
| Human Diseases | Infectious disease: viral | 0.00009 | 0.00019* | 0.00021** | 0.00007** | 0.00019* | 0.00017 |
| Human Diseases | Cancer: overview | 0.00153 | 0.00161 | 0.00163 | 0.00148 | 0.00155 | 0.00148 |
| Human | Cancer: specific types | 0.00051 | 0.00072** | 0.00075*** | 0.00048 | 0.00069** | 0.00065* |

| | | | | | | | |
|---|---|---|---|---|---|---|---|
| Diseases | | | | | | | |
| Human Diseases | Immune disease | 0.00052 | 0.00043* | 0.00042* | 0.00054 | 0.00046 | 0.00048 |
| Human Diseases | Cardiovascular disease | 0.00005 | 0.00009* | 0.00010* | 0.00003 | 0.00009* | 0.00007 |

Note: The table represented the average value of KEGG pathways level 2 and the belonged level 1. The pathways marked with yellow background with the *, **, *** mean the significantly differed between various life stages (F1 vs F3, F1 vs F5, M1 vs M3, M1 vs M5) with the p value <0.05, 0.01, 0.001.

**Table S5. The isolation and identification of zebrafish gut bacteria in NCBI-BLAST 16S ribosomal RNA (Bacteria and Archaea) database.**

| No. | Blast matched results from 16S Database | Identity (%) | Genus group | Sequence length (bp) | Samples' name and Similarity |
|---|---|---|---|---|---|
| 1. | Pseudomonas khazarica strain TBZ2 16S ribosomal RNA；<br>NR_169334.1；<br>Pseudomonas khazarica sp. nov., a polycyclic aromatic hydrocarbon-degrading bacterium isolated from Khazar Sea sediments；<br>Antonie Van Leeuwenhoek 113 (4), 521-532 (2020). | 99.79 % | Proteobacteria;<br>Gammaproteobacteria;<br>Pseudomonadales;<br>Pseudomonadaceae;<br>Pseudomonas. | 1416 | No.1=g1907088210-Y-2;<br>No.8=g1907088215-Y-7;<br>No.9=g1907303530-3;<br>N0.13=g2001223468-4;<br>No.20=g2007015126-7;<br>No.1-No.8-No.13-No.20=100%；<br>No.1-No.8-No.9-No.13-No.20=99.93%; |
| 2. | Shewanella seohaensis strain S7-3 16S ribosomal RNA;<br>NR_108852.1;<br>Shewanella seohaensis sp. nov., isolated from a tidal flat sediment;<br>Antonie Van Leeuwenhoek 102 (1), 149-156 (2012). | 99.86 % | Proteobacteria;<br>Gammaproteobacteria;<br>Alteromonadales;<br>Shewanellaceae;<br>Shewanella. | 1421 | No.2=g1907088211-Y-3;<br>N0.16=g2001223473-9;<br>No.2-No.16=100%; |
| 3. | Vibrio owensii CAIM 1854 = LMG 25443 strain DY05 16S ribosomal RNA；<br>NR_117424.1；<br>Vibrio owensii sp. nov., isolated from cultured crustaceans in Australia；<br>FEMS Microbiol. Lett. 302 (2), 175-181 (2010). | 99.65 % | Proteobacteria;<br>Gammaproteobacteria;<br>Vibrionales;<br>Vibrionaceae;<br>Vibrio. | 1428 | No.3=g1907088213-Y-5; |
| 4. | Vibrio alfacsensis strain CAIM 1831 16S ribosomal RNA；<br>NR_118129.1；<br>Vibrio alfacsensis sp. nov., isolated from marine organisms；<br>Int. J. Syst. Evol. Microbiol. 62 (PT 12), 2955-2961 (2012) | 99.93 % | Proteobacteria;<br>Gammaproteobacteria;<br>Vibrionales;<br>Vibrionaceae;<br>Vibrio. | 1435 | No.4=g1907088217-10-2;<br>No.3-No.4=96.38%; |

| | | | | | |
|---|---|---|---|---|---|
| 5. | Vibrio alginolyticus strain NBRC 15630 16S ribosomal RNA；<br>NR_113781.1；<br>Direct Submission；<br>Submitted (23-APR-2014) NCBI, NIH, Bethesda, MD 20894, USA. | 99.72% | Proteobacteria;<br>Gammaproteobacteria;<br>Vibrionales;<br>Vibrionaceae;<br>Vibrio. | 1429 | No.5=g1907088218-10-3;<br>No.17=g2007015121-2;<br>No.5-No.17=99.79%;<br>No.3-No.5=99.02%;<br>No.4-No.5=96.80%;<br>No.3-No.17=98.95%；<br>No.4-No.17=96.94%； |
| 6. | Vibrio brasiliensis LMG 20546 16S ribosomal RNA；<br>NR_117887.1；<br>Vibrio caribbeanicus sp. nov., isolated from the marine sponge Scleritoderma cyanea；<br>Int. J. Syst. Evol. Microbiol. 62 (PT 8), 1736-1743 (2012). | 98.60% | Proteobacteria;<br>Gammaproteobacteria;<br>Vibrionales;<br>Vibrionaceae;<br>Vibrio;<br>Vibrio oreintalis group | 1431 | No.6=g1907088221-10-6；<br>No.3-No.6=96.37%;<br>No.4-No.6=95.61%;<br>No.5-No.6=96.65%;<br>No.17-No.6=96.72%; |
| 7. | Staphylococcus succinus subsp. succinus strain AMG-D1 16S ribosomal RNA;<br>NR_028667.1;<br>Staphylococcus succinus sp. nov., isolated from Dominican amber;<br>Int. J. Syst. Bacteriol. 48 (Pt 2), 511-518 (1998). | 99.93% | Firmicutes;<br>Bacilli;<br>Bacillales;<br>Staphylococcaceae;<br>Staphylococcus. | 1429 | No.7=g1907303530-1；<br>No.10=g1907303530-2；<br>N0.11=g2001223466-2;<br>No.7-No.10-No.11 =100%; |
| 8. | Dietzia aurantiaca strain CCUG 35676 16S ribosomal RNA;<br>NR_108518.1;<br>Dietzia aurantiaca sp. nov., isolated from a human clinical specimen;<br>Int. J. Syst. Evol. Microbiol. 62 (PT 3), 484-488 (2012). | 99.06% | Actinobacteria;<br>Corynebacteriales;<br>Dietziaceae;<br>Dietzia. | 1375 | N0.12=g2001223467-3;<br>N0.14=g2001223471-7;<br>N0.15=g2001223472-8;<br>No.21=g2007015127-8<br>No.22=g2007015128-9;<br>No.14-No.15-No.21-No.22 =100%；<br>No.12-No.14-No.15-No.21-No.22=99.93%; |
| 9.* | Vibrio fluvialis strain NBRC 103150 16S ribosomal RNA；<br>NR_114218.1；<br>Direct Submission | 100% | Proteobacteria;<br>Gammaproteobacteria;<br>Vibrionales;<br>Vibrionaceae;<br>Vibrio. | 1433 | No.18=g2007015122-3;<br>No.3-No.18=96.15%;<br>No.4-No.18=97.14%;<br>No.5-No.18=96.72%;<br>No.6-No.18=96.73%;<br>No.17-No.18=96.86%; |

| | | | | | |
|---|---|---|---|---|---|
| | Submitted (23-APR-2014)<br>NCBI, NIH, Bethesda, MD<br>20894, USA. | | | | |
| 9.* | Allomonas enterica strain<br>LMG 21754 16S ribosomal<br>RNA；<br>NR_114900.1;<br>Direct Submission<br>Submitted (23-APR-2014)<br>NCBI, NIH, Bethesda, MD<br>20894, USA. | 99.65% | Proteobacteria;<br>Gammaproteobacte<br>ria;<br>Vibrionales;<br>Vibrionaceae;<br>Allomonas. | 1433 | No.18=g2007015122-3; |
| 10. | Bacillus cereus ATCC 14579<br>16S ribosomal RNA (rrnA);<br>NR_074540.1;<br>Genome sequence of Bacillus<br>cereus and comparative<br>analysis with Bacillus<br>anthracis;<br>Nature 423 (6935), 87-91<br>(2003). | 99.79% | Firmicutes;<br>Bacilli;<br>Bacillales;<br>Bacillaceae;<br>Bacillus;<br>Bacillus cereus<br>group. | 1444 | No.19=g2007015125-6; |

**Note:** 1.The results represented the best matched strains with the identity >98.6%, which included the information of bacterial name and version, source and submission, published journal and years. 2.The publish information was the first sample if there were multiple similar bacterial strains. 3.The identity of similar bacterial strains (marked in red color) were also searched by BLAST, and which less than <96% was considered as the different species. *means the first and second best matched bacteria were distinct at genus level.

**Table S6. The samples collection and sterile detection of GF *Artemia* models**

| Observe time / Samples | 24 h | 48 h | 72 h | 96 h | 120 h | 144 h | 168 h | Note |
|---|---|---|---|---|---|---|---|---|
| 1-TSA-normal cysts | + | + | + | + | + | + | + | |
| 2-TSA-treated cysts | - | - | - | - | - | - | - | 24 h-incubated nauplii |
| 3-TSB normal cysts | + | + | + | + | + | + | + | |
| 4-TSB-treated cysts | - | - | - | - | - | - | - | 24 h-incubated nauplii |
| 5-BHI normal cysts | + | + | + | + | + | + | + | |
| 6-BHI-treated cysts | - | - | - | - | - | - | - | 24 h-incubated nauplii |
| 7-TSA-GF-A | | - | - | - | - | - | - | |
| 8-TSA-hatching brine | | - | - | - | - | - | - | |
| 9-Blood TSA-GF-A | | - | - | - | - | - | - | |
| 10-Blood TSA-hatching brine | | - | - | - | - | - | - | |
| 11-Double TSA-GF-A | | - | - | - | - | - | - | |
| 12-Double TSA-hatching brine | | - | - | - | - | - | - | |
| 13-TSB-aerobic-GF-A | | - | - | - | - | - | - | |
| 14-TSB-aerobic-hatching brine | | - | - | - | - | - | - | |
| 15-BHI- | | - | - | - | - | - | - | |

| | | | | | | | |
|---|---|---|---|---|---|---|---|
| aerobic-GF-A | | | | | | | |
| 16-BHI-aerobic-hatching brine | | - | - | - | - | - | - |
| 17-TSB-anaerobic-GF-A | | - | - | - | - | - | - |
| 18-TSB-anaerobic-hatching brine | | - | - | - | - | - | - |
| 19-BHI-anaerobic-GF-A | | - | - | - | - | - | - |
| 20-BHI-anaerobic-hatching brine | | - | - | - | - | - | - |

Note: The time was begin at the treatment of *Artemia* cysts, and “+” means the bacterial colony or liquid, “-”means no bacterial cultured from the samples, which indicated the GF condition of models.

**Table S7. The developmental indexes of GF *Artemia* after 24 h incubation.**

| GF *Artemia* | Hatching rate % | Survival rate % | Malformation rate % | Body length (μm) | Body weight (mg) |
|---|---|---|---|---|---|
| Group 1 | 93.33 | 90.00 | 2.78 | 555.00 | 3.50 |
| Group 2 | 86.67 | 73.33 | 3.57 | 552.57 | 2.90 |
| Group 3 | 90.00 | 80.00 | 0.00 | 548.60 | 3.10 |
| Group 4 | 92.86 | 85.71 | 3.13 | 549.91 | 3.70 |
| Group 5 | 86.67 | 73.33 | 3.33 | 552.41 | 3.60 |
| Group 6 | 86.67 | 77.78 | 0.00 | 530.24 | 4.10 |
| Group 7 | 90.00 | 80.00 | 2.86 | 536.64 | 3.90 |
| Group 8 | 86.84 | 73.68 | 0.00 | 537.22 | 4.30 |
| Group 9 | 86.67 | 73.33 | 0.00 | 548.86 | 5.10 |
| Group 10 | 81.82 | 66.67 | 3.03 | 555.57 | --- |
| Average | 88.15 | 77.38 | 1.87 | 546.70 | 3.80 |
| Note | N=30-50 *Artemia* nauplii and cysts in each group | N=30-50 *Artemia* nauplii in each group | N=30-50 *Artemia* nauplii in each group | N=10 *Artemia* nauplii in each group | N=100 *Artemia* nauplii in each group, wet weight |

**Table S8. The samples and sterile detection of sterile micro-particle food and shelled shrimp eggs.**

| Detections / Samples | TSA | TSA double layer | Blood TSA | TSB-aerobic | TSB-anaerobic | BHI-aerobic | BHI-anaerobic |
|---|---|---|---|---|---|---|---|
| 1-micro-particle food (<50) | - | - | - | - | - | - | - |
| 2-micro-particle food (<50) after stored for 5 days | - | - | - | - | - | - | - |
| 3-micro-particle food (100-150) | - | - | - | - | - | - | - |
| 4-micro-particle food (100-150) after stored for 5 days | - | - | - | - | - | - | - |
| 5-sterile shelled shrimp eggs | - | - | - | - | - | - | - |
| 6- sterile shelled shrimp eggs after stored for 5 days | - | - | - | - | - | - | - |
| 7-GF-A | - | - | - | - | - | - | - |
| 8-GF-A after stored for 2 days | - | - | - | - | - | - | - |
| 9-negtive control | - | - | - | - | - | - | - |
| 10-positive control | + | + | + | + | + | + | + |

Note: The negative control was the GMM solution; the positive control was the culture water of *Artemia* in normal conditions.

**Table S9. The detections of samples collected during the GF marine medaka model from larvae to juvenile and adult stages.**

| Detections / Samples | TSA plates | TSA double layer plates | Blood TSA | TSB-aerobic | TSB-anaerobic | BHI-aerobic | BHI-anaerobic |
|---|---|---|---|---|---|---|---|
| 1-waste water | - | - | - | - | - | - | - |
| 2-broken embryo membrane and dead fish | - | - | - | - | - | - | - |
| 3-faeces | - | - | - | - | - | - | - |
| 4-negtive control | - | - | - | - | - | - | - |
| 5-positive control | + | + | + | + | + | + | + |

Note: The negative control was the GMM solution; the positive control was the culture water of *Artemia* in normal conditions.

**Table S10. The critical DEGs selected by key words of "growth" and "development" in different GF and CR marine medaka at early adult and adult stages.**

| Gene name | Gene description | Pathway definition (presented the crucial functions related to growth and development) | COG functional categories | Regulate and fold change (FC) | P-adjust value |
|---|---|---|---|---|---|
| | | **GF-2-VS-CR-2, Growth, Development** | | | |
| *gsdf* | gonadal somatic cell derived factor | --- | S:Function unknown | Down; 0.346 | 0.0112 |
| *LOC112152490* | growth hormone receptor | Growth hormone synthesis, secretion and action; Jak-STAT signaling pathway; PI3K-Akt signaling pathway. | U:Intracellular trafficking, secretion, and vesicular transport | Down; 0.261 | 0.0077 |
| *igfals* | insulin-like growth factor binding protein, acid labile subunit | Growth hormone synthesis, secretion and action | S | Down; 0.329 | 0.0223 |
| *dio2* | iodothyronine deiodinase 2 | Thyroid hormone signaling pathway | S | Down; 0.157 | 0.000002 |
| *tgfb5* | transforming growth factor, beta 5 | Cell cycle; Renal cell carcinoma; Pancreatic cancer; | O:Posttranslational modification, protein turnover, chaperones | Down; 0.243 | 0.0054 |
| *egfl7* | EGF-like-domain, multiple 7 | --- | S | Down; 0.440 | 0.0306 |
| *cgref1* | cell growth regulator with EF-hand domain 1 | --- | S | Down; 0.146 | 0.0001 |
| *igf1* | insulin-like growth factor 1, transcript variant X1 | Ovarian steroidogenesis; Growth hormone synthesis, secretion and action;PI3K-Akt signaling pathway; | K:Transcription | Down; 0.197 | 0.00007 |
| *gh1* | growth hormone 1 | Growth hormone synthesis, secretion and action; Jak-STAT signaling pathway; PI3K-Akt signaling pathway | O | Down; 0.159 | 0.000000002 |
| *esm1* | endothelial cell-specific molecule 1 | --- | O | Down; 0.173 | 0.0005 |
| *epcam* | epithelial cell adhesion molecule | --- | U | Down; 0.388 | 0.0229 |
| *junbb* | JunB proto-oncogene, AP-1 transcription factor subunit b | Osteoclast differentiation; Growth hormone synthesis, secretion and action; TNF signaling pathway | K:Transcription | Down; 0.401 | 0.0241 |
| | | **GF-3-VS-CR-3, Growth, Development** | | | |
| *gsdf* | gonadal somatic cell derived factor | --- | S:Function unknown | Down; 0.423 | 0.0090 |
| *eps8a* | epidermal growth factor receptor pathway substrate 8a, transcript variant X1 | --- | T:Signal transduction mechanisms | Up; 2.055 | 0.0001 |
| *fgf16* | fibroblast growth factor 16, transcript | Ras signaling pathway; MAPK signaling pathway; PI3K-Akt signaling | T | Down; 0.423 | 0.0090 |

| | | | | | |
|---|---|---|---|---|---|
| | variant X1 | pathway | | | |
| *igfals* | insulin-like growth factor binding protein, acid labile subunit | Growth hormone synthesis, secretion and action | S | Down; 0.448 | 0.0022 |
| *tgfbr2b* | transforming growth factor beta receptor 2b | Pancreatic cancer; Th17 cell differentiation; MAPK signaling pathway; TGF-beta signaling pathway | T | Down; 0.456 | 0.0022 |
| *pdgfc* | platelet derived growth factor c | Ras signaling pathway; Gap junction; MAPK signaling pathway; EGFR tyrosine kinase inhibitor resistance; PI3K-Akt signaling pathway | A:RNA processing and modification | Down; 0.406 | 0.000002 |
| *gdf6a* | growth differentiation factor 6a | Cytokine-cytokine receptor interaction; TGF-beta signaling pathway; Hippo signaling pathway | O | Down; 0.462 | 0.0042 |
| *igf1* | insulin-like growth factor 1, transcript variant X1 | Ovarian steroidogenesis; Growth hormone synthesis, secretion and action;PI3K-Akt signaling pathway; | K:Transcription | Down; 0.241 | 4.15E-21 |
| *egr2a* | early growth response 2a | Human T-cell leukemia virus 1 infection; Hepatitis B | S | Up; 4.664 | 0.00045 |
| *fgf23* | fibroblast growth factor 23 | Rap1 signaling pathway; Ras signaling pathway; MAPK signaling pathway; Parathyroid hormone synthesis, secretion and action; PI3K-Akt signaling pathway | T | Down; 0.184 | 0.0210 |
| *pdgfrb* | platelet-derived growth factor receptor, beta polypeptide | Focal adhesion; Regulation of actin cytoskeleton; Calcium signaling pathway; Jak-STAT signaling pathway; EGFR tyrosine kinase inhibitor resistance;PI3K-Akt signaling pathway | T | Down; 0.490 | 0.0051 |
| *fgfr2* | fibroblast growth factor receptor 2, transcript variant X1 | Ras signaling pathway; Signaling pathways regulating pluripotency of stem cells; EGFR tyrosine kinase inhibitor resistance; PI3K-Akt signaling pathway | T | Down; 0.472 | 0.0000006 |
| *egr1* | early growth response 1 | Parathyroid hormone synthesis, secretion and action; GnRH signaling pathway | S | Down; 0.296 | 0.000002 |
| *pkd2* | polycystic kidney disease 2, transcript variant X1 | --- | S | Down; 0.437 | 0.000000002 |
| *fgf16* | fibroblast growth factor 16, transcript variant X1 | Ras signaling pathway; MAPK signaling pathway; PI3K-Akt signaling pathway | T | Down; 0.316 | 0.0070 |
| *LOC112150763* | insulin-like growth factor-binding protein 3 | Growth hormone synthesis, secretion and action; Cellular senescence; p53 signaling pathway | A | Down; 0.497 | 0.0007 |
| *ngfb* | nerve growth factor b (beta polypeptide) | Rap1 signaling pathway; Ras signaling pathway; MAPK signaling pathway; Neurotrophin signaling pathway; PI3K-Akt signaling pathway | T | Down; 0.054 | 0.0058 |
| *gcm2* | glial cells missing transcription factor 2, transcript variant X2 | Parathyroid hormone synthesis, secretion and action | K | Down; 0.376 | 0.0000009 |
| *mmp1* | matrix | Parathyroid hormone synthesis, | O | Down; | 0.0058 |

| | | | | | |
|---|---|---|---|---|---|
| *4b* | metallopeptidase 14b (membrane-inserted) | secretion and action; GnRH signaling pathway; TNF signaling pathway | | 0.054 | |
| *spaca 4l* | sperm acrosome associated 4 like | --- | S | Down; 0.470 | 0.0041 |
| *gnrh3* | gonadotropin-releasing hormone 3 | GnRH secretion; GnRH signaling pathway; Neuroactive ligand-receptor interaction | U | Up; 97.94 | 0.00000009 |
| *LOC112140833* | solute carrier family 2, facilitated glucose transporter member 1 | Thyroid hormone signaling pathway; Bile secretion; Insulin secretion; Insulin resistance | G:Carbohydrate transport and metabolism | Up; 8.197 | 0.00000000004 |
| *six5* | SIX homeobox 5 | Transcriptional misregulation in cancer | K | Down; 0.337 | 0.0010 |
| *mmp14a* | matrix metallopeptidase 14a (membrane-inserted) | Parathyroid hormone synthesis, secretion and action; GnRH signaling pathway; TNF signaling pathway | O | Down; 0.224 | 4.33E-16 |
| **GF-3-VS-GF-2, Growth, Development** | | | | | |
| *pdgfrl* | platelet-derived growth factor receptor-like | --- | T | Up; 2.705 | 0.0040 |
| *tgfbr2b* | transforming growth factor beta receptor 2b | Pancreatic cancer; TGF-beta signaling pathway; Adherens junction | T | Up; 3.388 | 0.0031 |
| *tgfb5* | transforming growth factor, beta 5 | Cell cycle; TGF-beta signaling pathway | O | Up; 3.550 | 0.0005 |
| *socs2* | suppressor of cytokine signaling 2, transcript variant X1 | Insulin signaling pathway; Growth hormone synthesis, secretion and action; Jak-STAT signaling pathway; Prolactin signaling pathway | O | Down; 0.414 | 0.0041 |
| *fgf1a* | fibroblast growth factor 1a | Regulation of actin cytoskeleton; Ras signaling pathway; PI3K-Akt signaling pathway | T | Down; 0.296 | 0.0019 |
| *mmp14b* | matrix metallopeptidase 14b (membrane-inserted) | Parathyroid hormone synthesis, secretion and action; GnRH signaling pathway; TNF signaling pathway | O | Up; 4.569 | 0.00001 |
| *gnrh3* | gonadotropin-releasing hormone 3 | GnRH secretion; GnRH signaling pathway; Neuroactive ligand-receptor interaction | U | Up; 17.409 | 0.0002 |
| **CR-3-VS-CR-2, Growth, Development** | | | | | |
| *pdgfrl* | platelet-derived growth factor receptor-like | --- | T | Up; 3.129 | 0.0000000006 |
| *gadd45ga* | growth arrest and DNA-damage-inducible, gamma a | Cell cycle; p53 signaling pathway; Pancreatic cancer; MAPK signaling pathway; Cellular senescence; Apoptosis; FoxO signaling pathway; NF-kappa B signaling pathway | O | Down; 0.196 | 6.58E-13 |
| *fgf7* | fibroblast growth factor 7 | Regulation of actin cytoskeleton; Ras signaling pathway; MAPK signaling pathway; PI3K-Akt signaling pathway | T | Up; 2.443 | 0.0015 |
| *fgfbp2a* | fibroblast growth factor binding | --- | A | Up; 10.516 | 0.00002 |

| | | | | | |
|---|---|---|---|---|---|
| | protein 2a, transcript variant X1 | | | | |
| *LOC112136844* | platelet-derived growth factor receptor beta, transcript variant X1 | MAPK signaling pathway; Calcium signaling pathway; Jak-STAT signaling pathway; EGFR tyrosine kinase inhibitor resistance; PI3K-Akt signaling pathway | T | Up; 3.027 | 0.000003 |
| *socs3a* | suppressor of cytokine signaling 3a | TNF signaling pathway; Jak-STAT signaling pathway; Insulin signaling pathway; Growth hormone synthesis, secretion and action | O | Down; 0.339 | 0.00004 |
| *tgfb1a* | transforming growth factor, beta 1a, transcript variant X2 | Cell cycle; TGF-beta signaling pathway | O | Up; 3.012 | 0.000001 |
| *tgfb5* | transforming growth factor, beta 5 | Cell cycle; TGF-beta signaling pathway; MAPK signaling pathway | O | Up; 3.221 | 0.0002 |
| *gdf10a* | growth differentiation factor 10a | Cytokine-cytokine receptor interaction | O | Up; 3.787 | 0.000004 |
| *gadd45ba* | growth arrest and DNA-damage-inducible, beta a | Cell cycle; FoxO signaling pathway; NF-kappa B signaling pathway | O | Down; 0.403 | 0.0000008 |
| *gdf6a* | growth differentiation factor 6a | Cytokine-cytokine receptor interaction; TGF-beta signaling pathway; Hippo signaling pathway | O | Up; 2.836 | 0.0000009 |
| *ngfra* | nerve growth factor receptor a (TNFR superfamily, member 16) | Rap1 signaling pathway; Ras signaling pathway; MAPK signaling pathway; Cytokine-cytokine receptor interaction; PI3K-Akt signaling pathway | U | Up; 3.229 | 0.0024 |
| *gh1* | growth hormone 1 | Cytokine-cytokine receptor interaction; Neuroactive ligand-receptor interaction; Growth hormone synthesis, secretion and action; Jak-STAT signaling pathway; PI3K-Akt signaling pathway | O | Down; 0.197 | 0.0077 |
| *fgf23* | fibroblast growth factor 23 | Ras signaling pathway; MAPK signaling pathway; Parathyroid hormone synthesis, secretion and action; PI3K-Akt signaling pathway | T | Up; 13.196 | 0.0001 |
| *ngfb* | nerve growth factor b (beta polypeptide) | Rap1 signaling pathway; Ras signaling pathway; MAPK signaling pathway; Neurotrophin signaling pathway; Cytokine-cytokine receptor interaction; PI3K-Akt signaling pathway | T | Up; 47.054 | 0.0004 |
| *isl1* | ISL LIM homeobox 1, transcript variant X1 | Signaling pathways regulating pluripotency of stem cells | K | Down; 0.315 | 0.0001 |
| *mmp14b* | matrix metallopeptidase 14b (membrane-inserted) | Parathyroid hormone synthesis, secretion and action; GnRH signaling pathway; TNF signaling pathway | O | Up; 7.186 | 1.36E-13 |
| *bmp4* | bone morphogenetic protein 4, transcript variant X2 | Signaling pathways regulating pluripotency of stem cells; TGF-beta signaling pathway; Thyroid hormone | O | Up; 3.078 | 4.82E-10 |

| | | | | | |
|---|---|---|---|---|---|
| | | signaling pathway | | | |
| *mmp14a* | matrix metallopeptidase 14a (membrane-inserted) | Parathyroid hormone synthesis, secretion and action; GnRH signaling pathway; TNF signaling pathway | O | Up; 4.049 | 4.04E-9 |

Note: “---” means no description or definition, and the short letter means the same gene and information as in the previous table. Some of the DEGs researched by key words “Growth”, “Development” included the DEGs related to key words of “Gonad”, “Reproduction”.

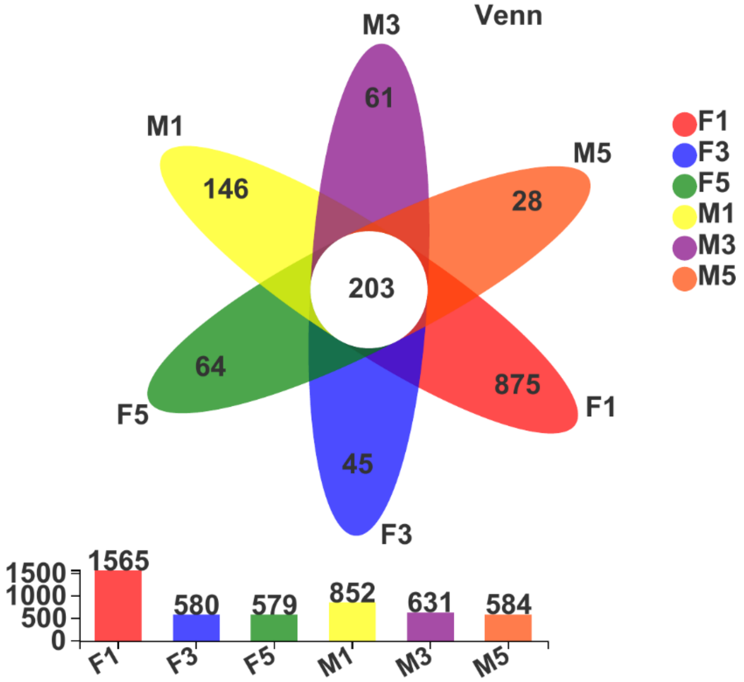

**Figure S1. The Venn map of gut microbiota in marine medaka at 1, 3, and 5 mpf based on the OTUs level**. From the map, it showed that the common OTUs of the six groups was 203, and the F1 group specially owned 875 OTUs, F3 was 45, F5 was 64, M1 was 146, M3 was 61, and M5 was 28. Moreover, the total number of each group was shown in the bottom.

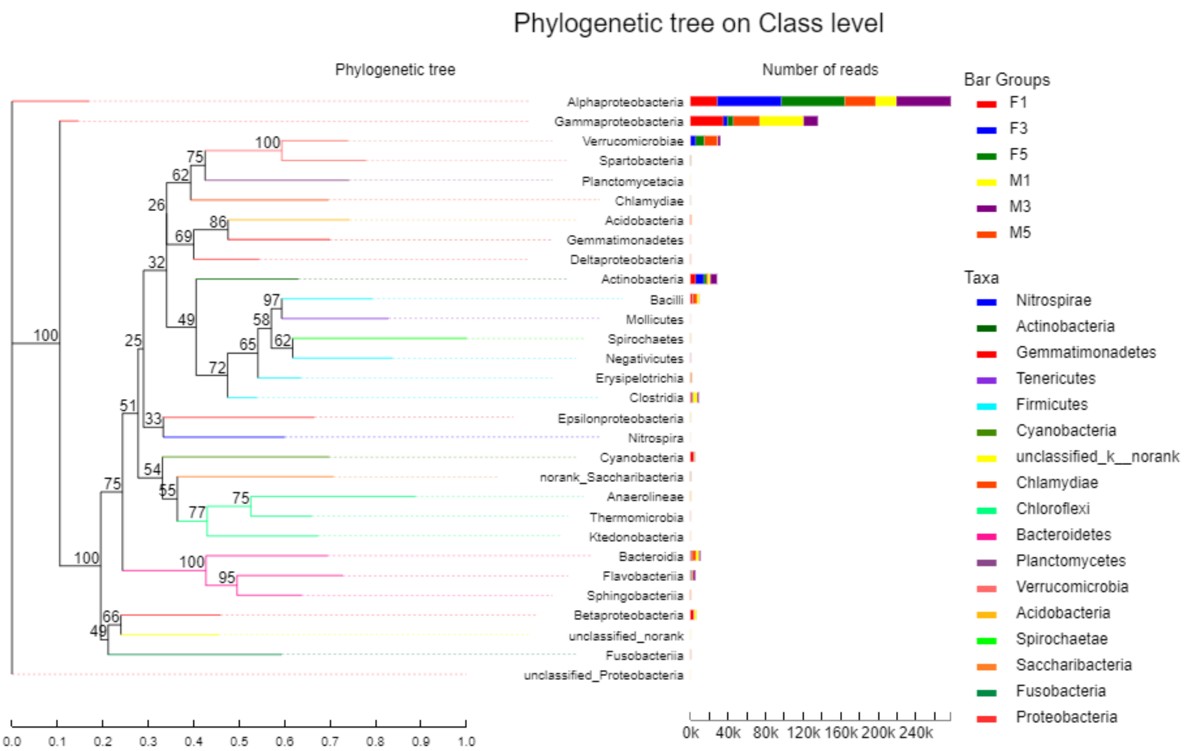

**Figure S2. The phylogenetic tree of gut microbiota in marine medaka at 1, 3, and 5 mpf based on class level.** The map showed that the distance and contains of different bacteria in the six groups on the class level, and the major of the class were belonged to Alphaproteobacteria, Gammaproteobacteria, Verrucomicrobia, Actinobacteria, and so on.

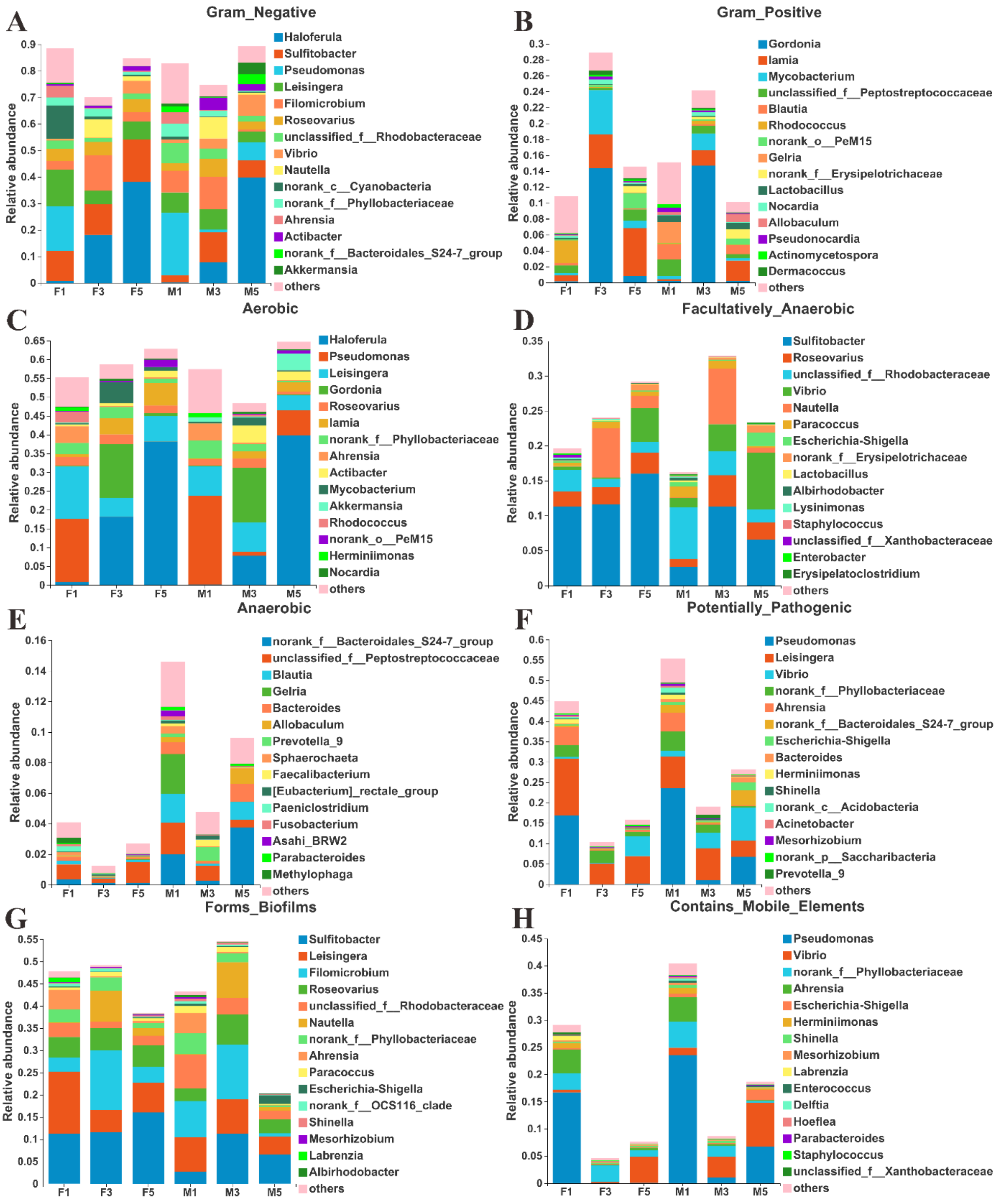

**Figure S3. The predicted microbial phenotypes of gut microbiota in marine medaka from juvenile to adult.** It showed that the predicted microbial phenotypes included the 8 major Gram_Negative (A) and Gram_Positive (B), Aerobic (C) and Facultatively-Anaerobic (D), Anaerobic (E) and Potentially-Pathogenic (F), Forms_Biofilms (G) and Contains_Mobile_Elements (H) phenotypes of intestinal microbiota, and analyzed the dominant genera presented with top 15 species contributed to phenotypes and potential functions in female and male fish at different life stages.

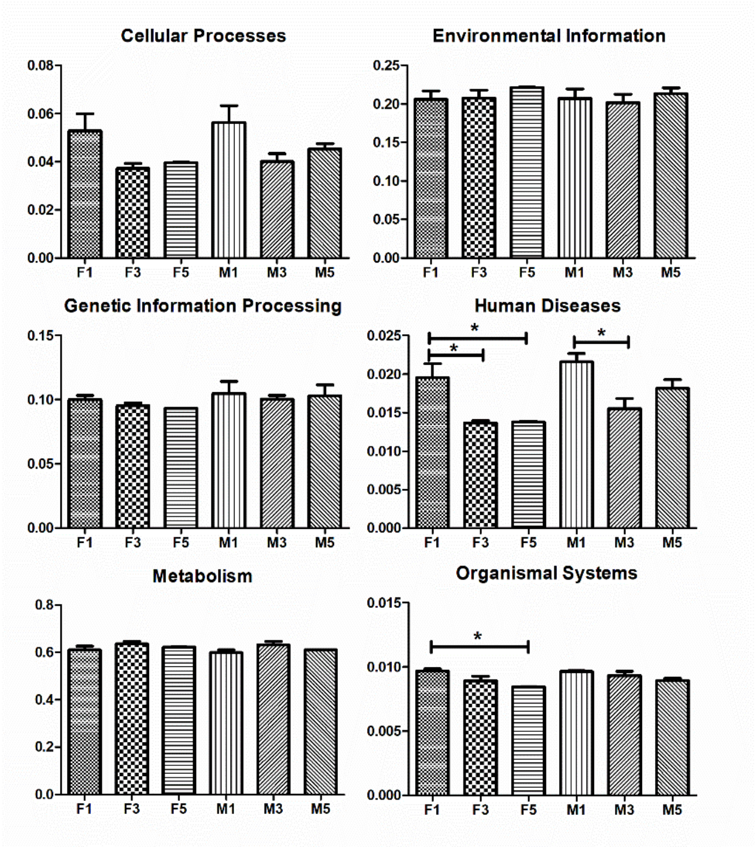

**Figure S4. The KEGG pathways of gut microbiota in marine medaka at different life stages.** In the level 1, the pathways of human diseases and organismal systems showed significantly different between F1 and F3, F5 groups.

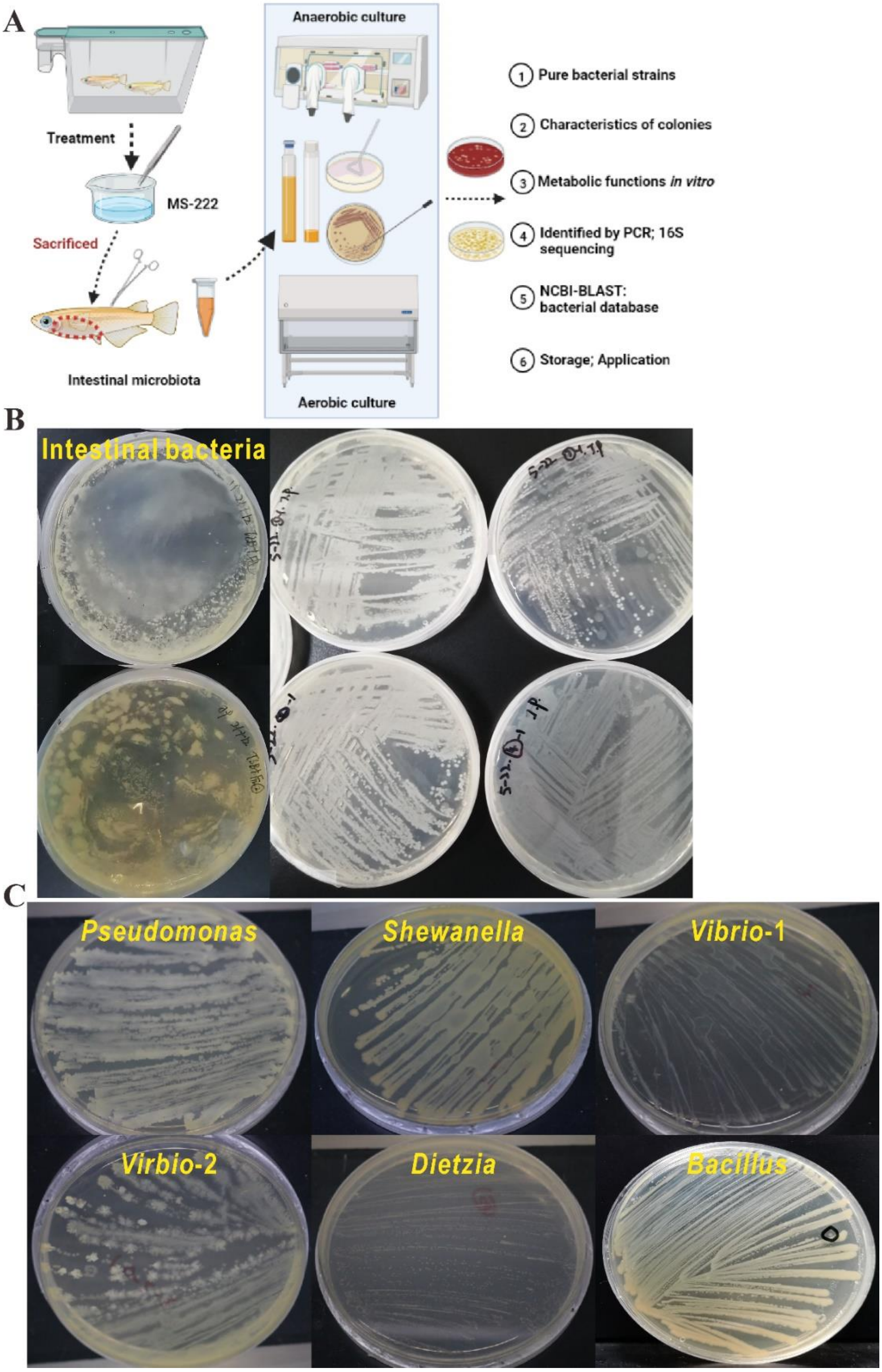

**Figure S5. The isolation and identification of intestinal bacteria in the adult marine medaka.** (A) The diagram of the progress, which involved the treatment of adult fish with washing and anaesthetized in MS-222, sacrificed to dissect the gut tissues, then culture the gut microbiota in the TSB and BHI medium, selected distinct bacterial colonies and purify the strains on TSA and Blood plates, identify the strains by PCR and 16S rRNA gene sequencing as well as the BLASTn search, finally store the bacteria and submitted the information on NCBI database. (B) It showed the multiple colonies on TSA plates after intestinal microbiota coated, and the subcultures before the pure strain gained. (C) It represented the bacterial colonies on plates with individual strains purified and genus identified.

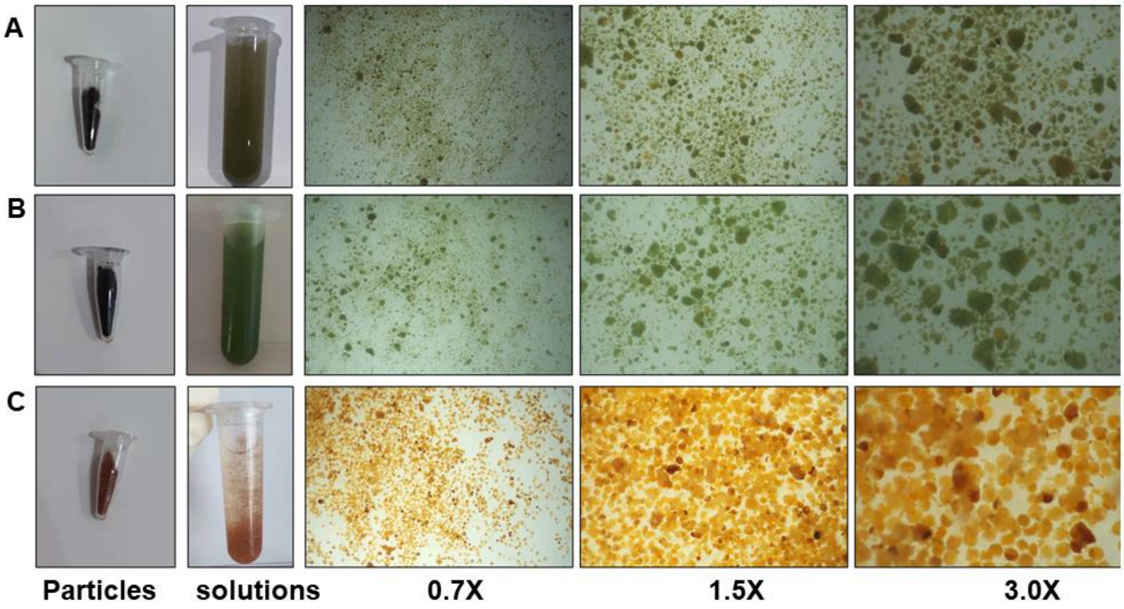

**Figure S6. The characters and comparison of the sterile micro-particle food and shelled shrimp eggs before feeding.** (A) The micro-particle food with <50 microns and (B) with 100-150 microns, and (C) shelled shrimp eggs samples prepared in this study.

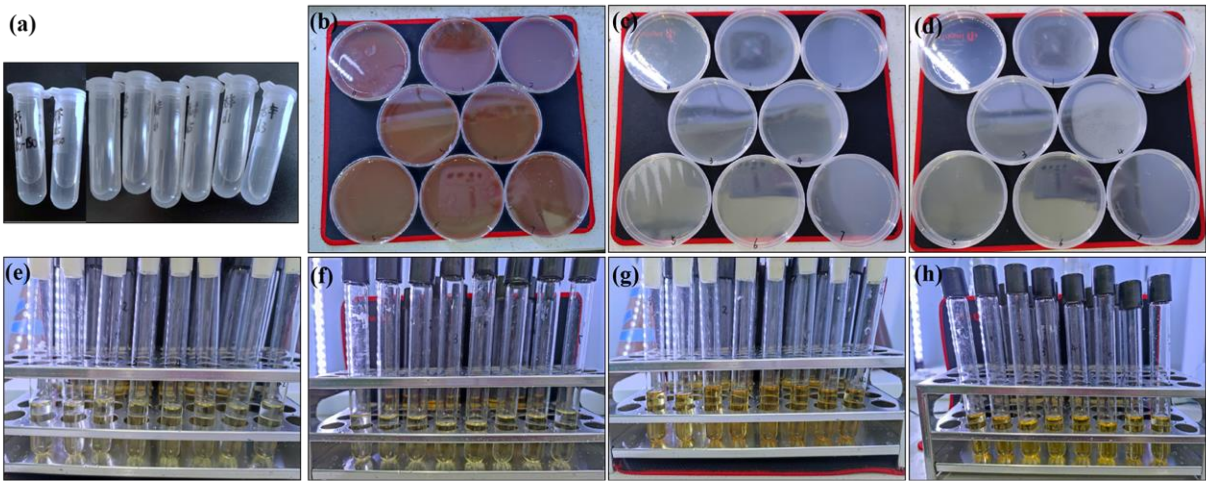

**Figure S7. The detection of the sterile micro-particle food and shelled shrimp eggs before feeding.** (a) The samples collected during the experiment. (b-d) The TSA, Blood plates and double layer plates. And (e-h) the TSB and BHI medium detection in the aerobic and anaerobic conditions.

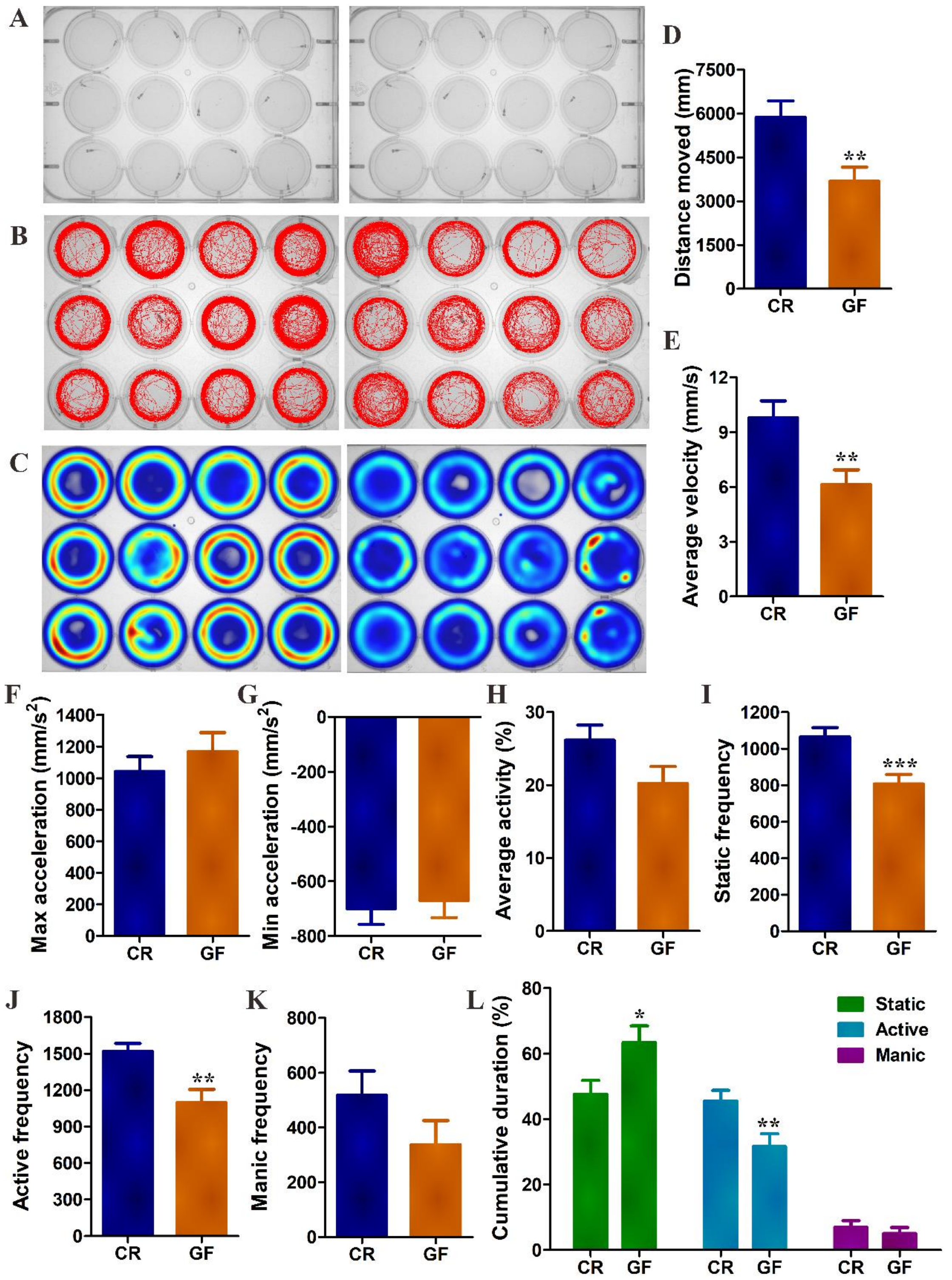

**Figure S8. The behavior differences between GF and CR marine medaka at 1 mpf.** (A) The juvenile fish in the multiple wells plate (n=21 replicates for each group) for the tracking analysis, (B) the visual images and (C) the heatmap for GF and CR fish tracking during 10 min data collection. (D) The distance moved (mm), and (E) the average velocity (mm/s), and (F, G) the max and min acceleration (mm/s$^2$) of marine medaka. (H) The average activity (%), and (I-J-K) the static, active, manic frequency in marine medaka. (L) The cumulative duration of marine medaka compared to total time 10 min was calculated and presented as the time ratio of fish at static, active, and manic states. The symbols of *, **, and *** stands for $p<0.05$, 0.01, and 0.001 with significant differences between CR and GF models.

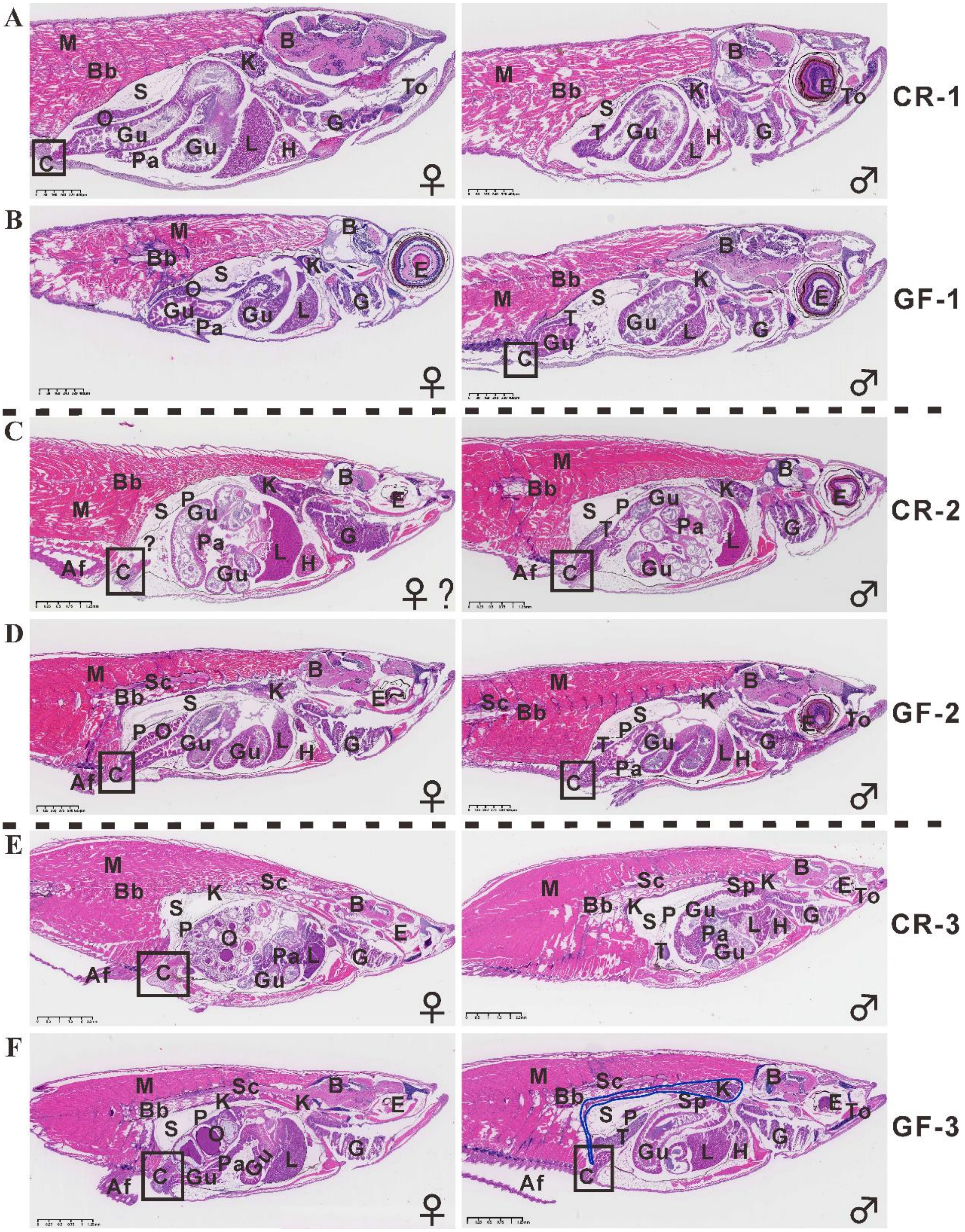

**Figure S9. The histopathological analysis of GF and CR marine medaka from juvenile to early adult and adult models with H&E stain.** (A, B) The juvenile fish at 1 mpf from CR and GF models, which departed by the female and male based on the developing gonad in sections with magnification 50×, scale bar of 400 μm. (C, D) The early adult fish at 2 mpf showed developed tissues or organs, but with obvious difference between CR fish with 20×, 1.25 mm and GF fish with 30×, 625 μm sections.

(E, F) The adult CR fish with 10×, 2.5 mm scale bar sections at 3 mpf appeared complete developed tissues to be the mature ones, but GF fish with 20×, 1.25 mm sections still become smaller size and delayed matured organs. There were n=3 replicate samples for each GF and CR group at different stages, and n=5 sections with special stain for each sample, which totally >180 sections were carried out in those experiments. Af: anal fin, B: brain, Bb: back bone, C: cloaca, E: eye, G: gill, Gu: gut, H: heart, K: kidney with the blue irregular circle, L: liver, M: muscle, O; ovary, P: peritoneum, Pa: pancreas, S: swim bladder, Sc: spinal cord, Sp: spleen, T: testis, To: tongue, ♀: female, ♂: male.

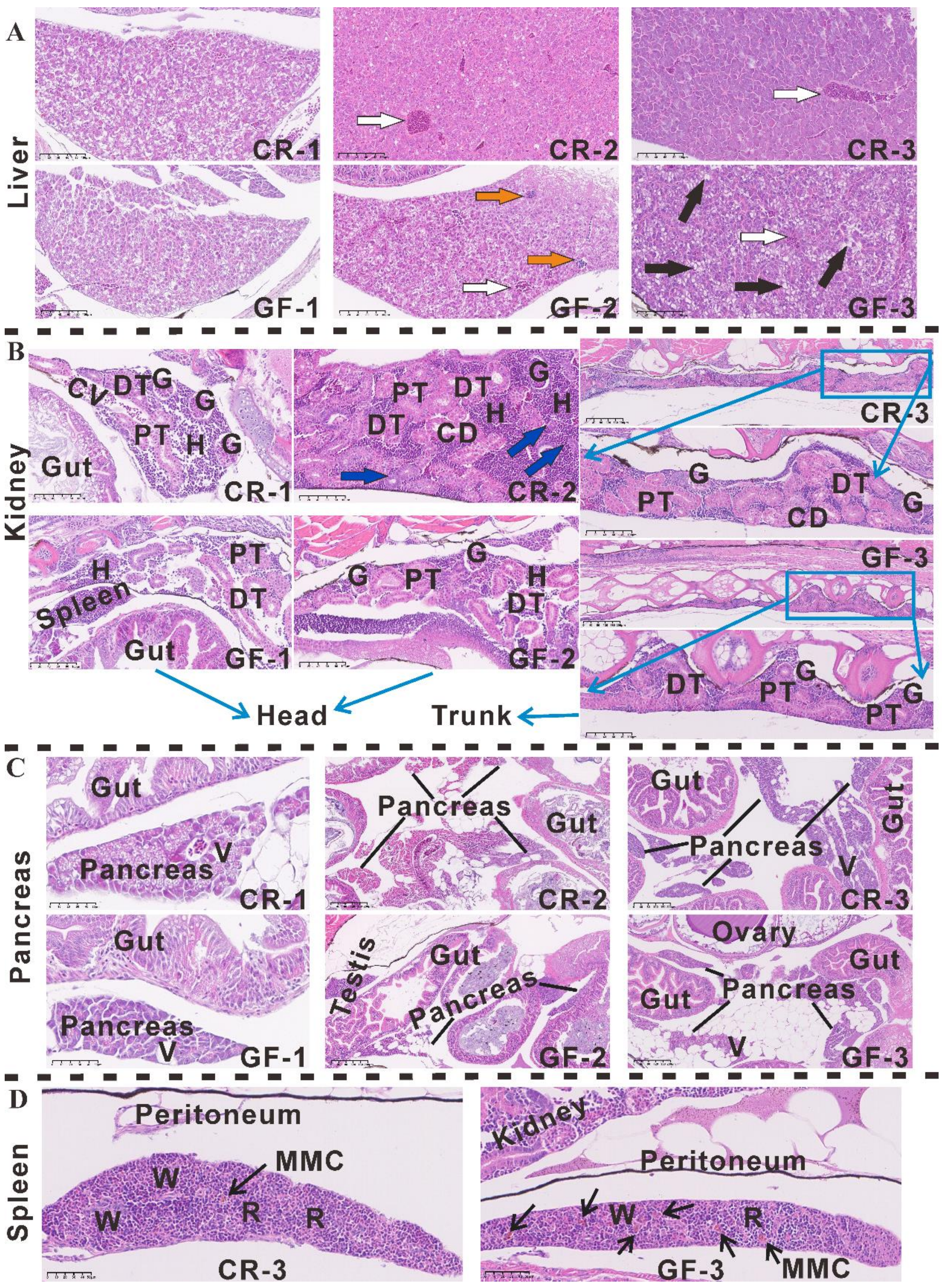

**Figure S10. The developmental differences of major immune organs between the GF and CR marine medaka.** (A) The liver tissue of GF and CR fish at 1, 2, 3 mpf, which showed the different developing degrees during the growth and between groups in sections with magnification 250×, scale bar of 100 μm. (B) The kidney tissue of GF and CR models, which showed the distinct and matured structure in normal fish head and trunk kidney, but less tubules and hematopoietic tissue in GF fish under the sections with magnification 250×, scale bar of 100 μm at 1, 2, 3 mpf, and 100×, scale bar of 200 μm in adult fish overall trunk. (C) The pancreas in GF and CR fish showed the relative intact and small tissues in both GF and CR fish at 1 mpf with magnification 500× and 50 μm scale bar, but the discrete multiple pancreas were scattered along the intestinal tract in GF and CR fish at 2 and 3 mpf with 100× and 200 μm sections. (D) The spleen of GF and CR fish at 3 mpf life stage with magnification 400× and 50 μm scale bar can be observed the grow-up and relative intact tissues nearby the gut and peritoneum, but GF fish still showed smaller size, indistinct difference of R and W, and more MMC in spleen tissue. In liver, the central veins (white arrow), suspected lymphocytes (orange arrow), and black arrow indicated the decrease of hepatocytes with rare hepatic density in GF fish along with the development and growth from 1 to 2 and 3 mpf stages. In the kidney: G=glomerulus, PT= proximal tubules, DT=distal tubules, CD=collecting duct, CV=central vein, H=hematopoietic tissue, and the newly generation of renal tubules (blue arrow). In pancreas, the smaller area of GF and serval vacuoles (V) were appeared in both GF and CR fish tissues. The spleen tissue showed the red pulp (R) and white pulp (W), and melano-macrophage center (MMC, thin arrow) with deep brown color by H&E stain.

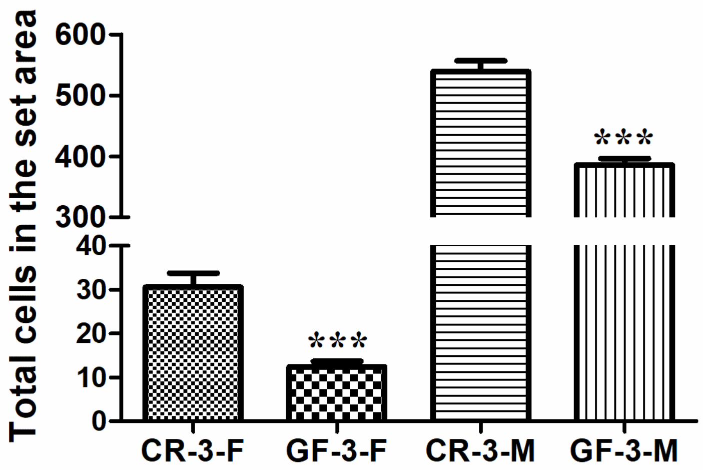

**Figure S11. The total cells of the set area in ovary and in testis from CR and GF fish at 3 mpf adult stage.** The total cells of oocytes at different stages were counted to be 12.47 in GF fish, which significantly less than 30.6 in CR fish. Similarly, the total cells in testis at different stages were calculated to be 385.87 in GF fish, which significantly less than 538.93 in CR fish. And the "***" stands for the significant difference with $p$ value <0.001 between CR and GF groups.

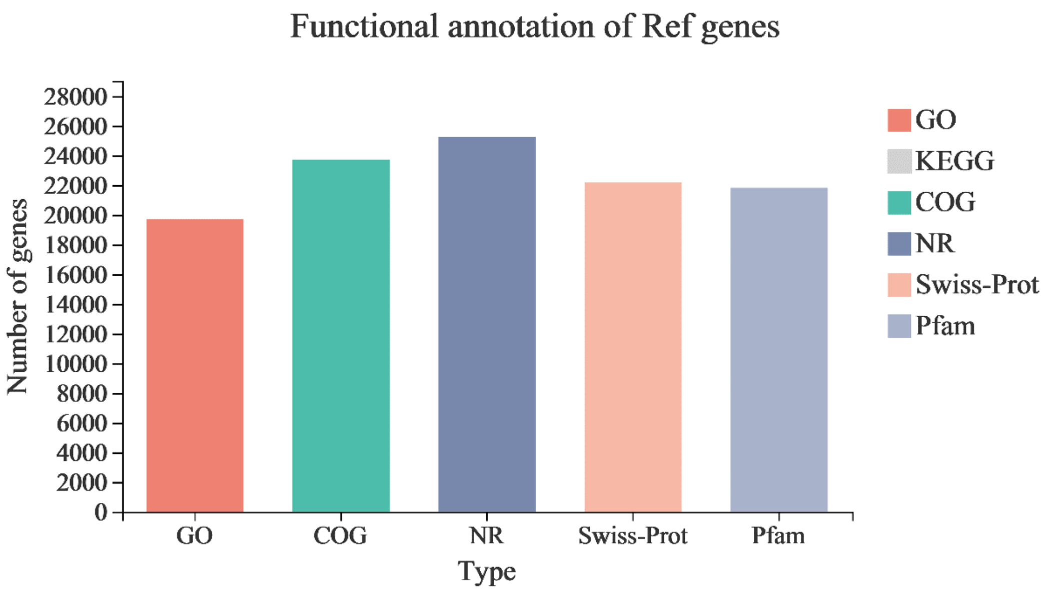

**Figure S12. The functional annotation of reference genes in GF and CR groups at different developed stages**. The database included the GO, KEGG, COG, NR, Swiss-Prot, and Pfam.

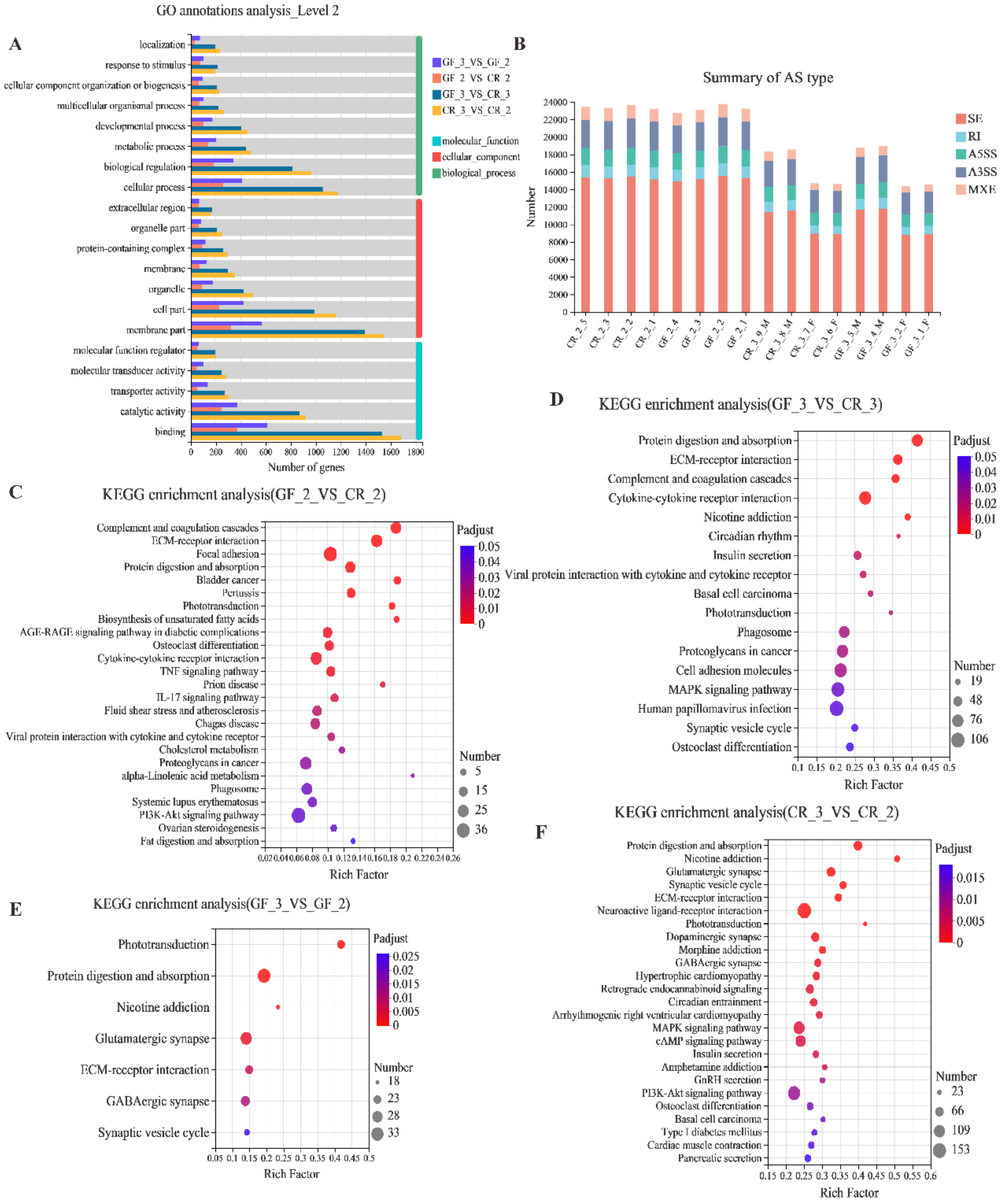

**Figure S13. The expression profile and changed pathways of GF and CR marine medaka at early-adult and adult stages.** (A) The annotations analysis of GO database at level 2 for expression profile in each sample, and the summary of AS type (B) of the sequencing reads in GF and CR fish at 2 and 3 mpf ages. (C) The enriched analysis of different KEGG pathway between GF-2 and CR-2 fish showed with top 25 of totally 28 significant changed pathways. (D) The enriched KEGG pathway between GF-3 and CR-3 fish was totally 17 significant changed pathways. (E) The enriched KEGG pathway between GF-3 and GF-3 fish showed totally 7 significant changed pathways.

(F) The enriched KEGG pathways showed top 25 of totally 37 pathways in CR-3-VS-CR-2 group. All the enriched KEGG pathways were selected with *p*-adjust value <0.05 from the DEGs sets saved by the *p*-adjust value <0.05 and presented with the FC ≥ 2 or FC ≤ 0.5. The vertical axis indicates KEGG terms and the horizontal axis represents the rich factor, and the enrichment degree was stronger with a bigger rich factor, and the size of dots indicates the number of DEGs.

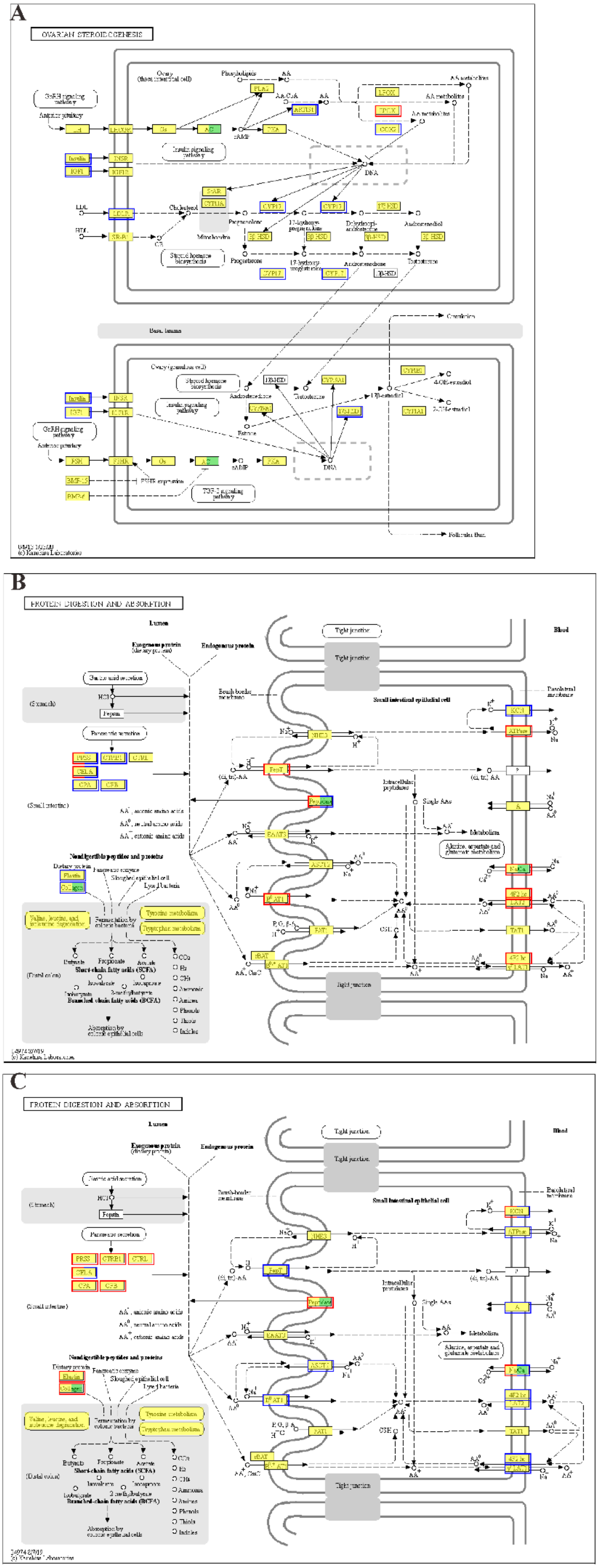
A
OVARIAN STEROIDOGENESIS
B
PROTEIN DIGESTION AND ABSORPTION
C
PROTEIN DIGESTION AND ABSORPTION

**Figure S14. The significantly changed KEGG pathways in GF and CR compared groups and at different developed stages.** (A) The critical KEGG pathway of Ovarian was significantly enriched in GF-2-VS-CR-2 group. The protein digestive and absorption pathway was the most changed in GF-3-VS-CR-3 (B) group and CR-3-VS-CR-2 (C) group. (D) The Pi3k-akt pathway was also significantly changed in CR-3-VS-CR-2 group. All the KEGG pathways were selected with the *p*-adjust value <0.05 among the saved DEGs sets, and then presented the detailed pathway maps with DEGs marked by red color of up-regulation and blue color of down-regulation.

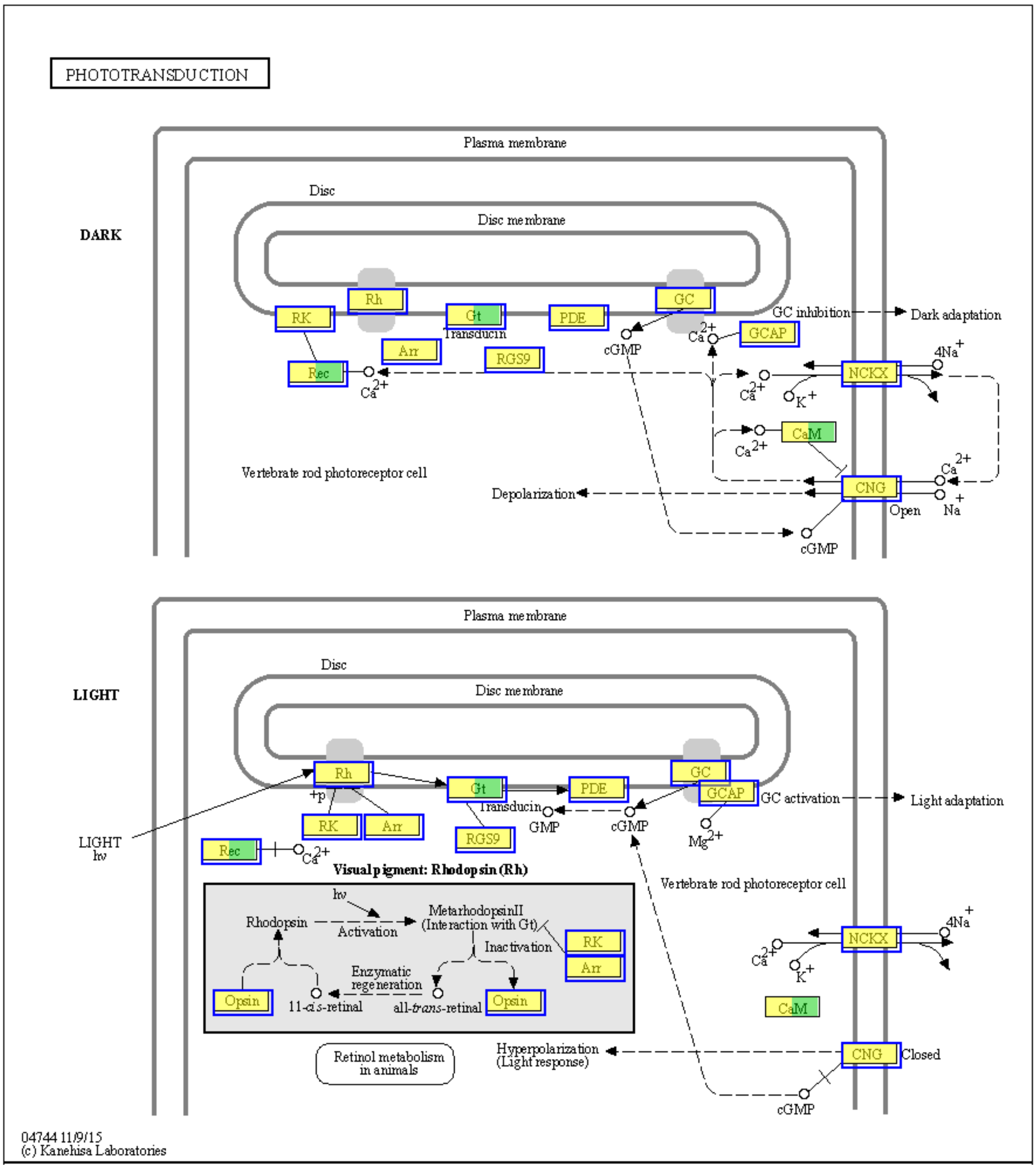

**Figure S15. The significantly changed KEGG pathways in GF and CR compared groups and at different developed stages.** The critical KEGG pathway of Phototransduction in GF-3VS-GF-2 group was the significantly changed and selected with the *p*-adjust value <0.05 among the saved DEGs sets, and then presented the detailed pathway maps with DEGs marked by red color of up-regulation and blue color of down-regulation.

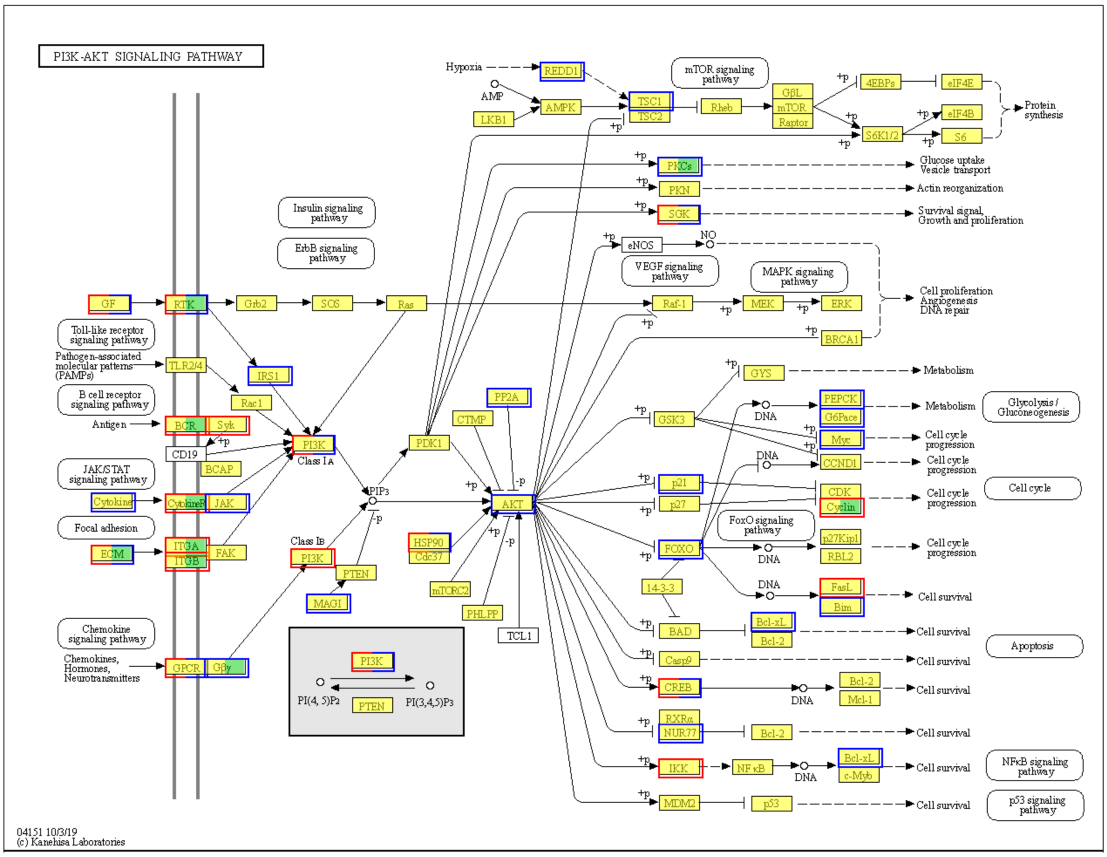

**Figure S16. The significantly changed KEGG pathways in GF and CR compared groups and at different developed stages.** The critical KEGG pathway of Pi3k-akt pathway was the significantly changed in CR-3-VS-CR-2 group selected with the *p*-adjust value <0.05 among the saved DEGs sets, and then presented the detailed pathway maps with DEGs marked by red color of up-regulation and blue color of down-regulation.

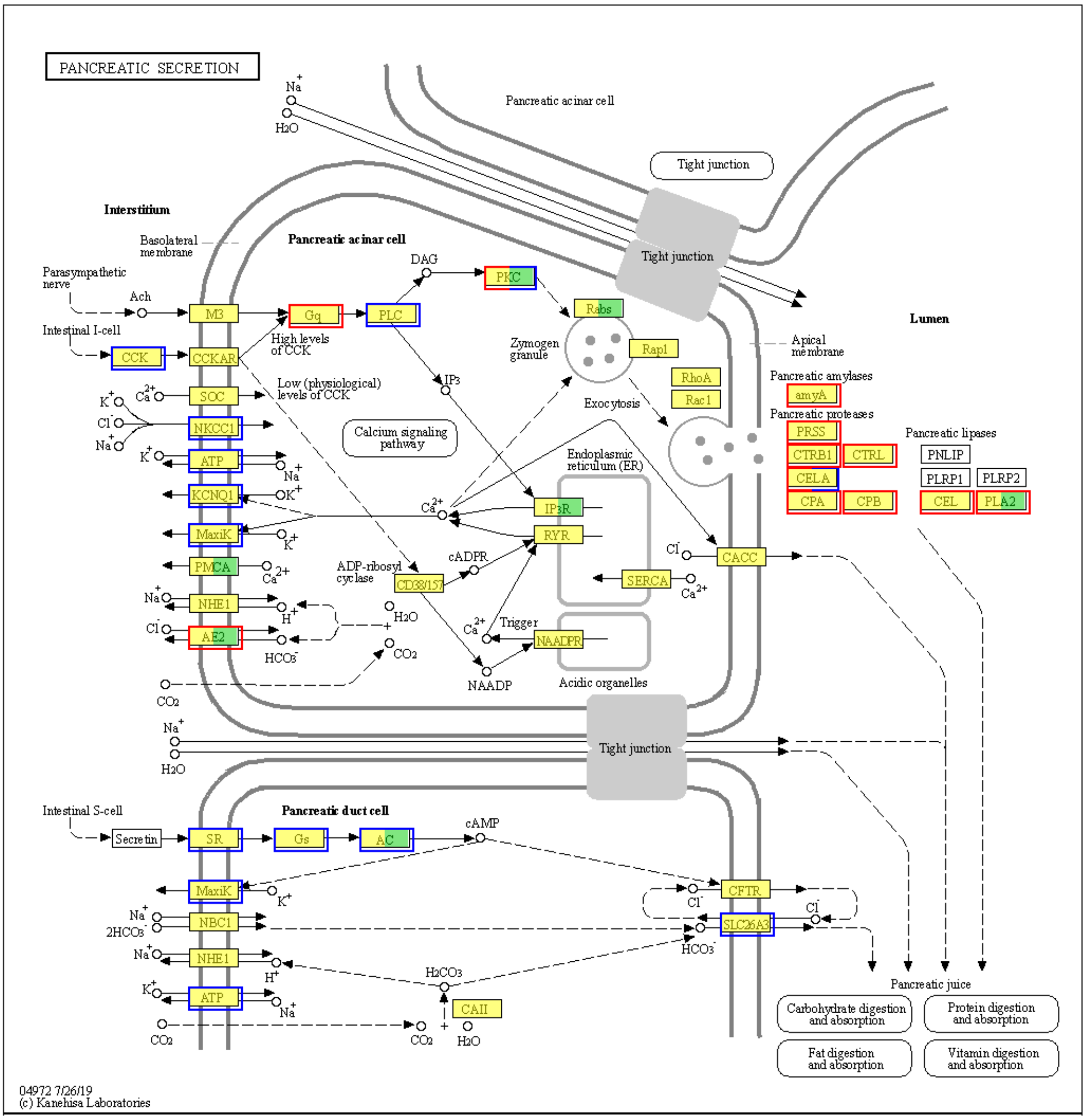

**Figure S17. The significantly changed KEGG pathways in GF and CR compared groups and at different developed stages.** The critical KEGG pathway of pancreatic secretion pathway was the significantly changed in CR-3-VS-CR-2 group selected with the *p*-adjust value <0.05 among the saved DEGs sets, and then presented the detailed pathway maps with DEGs marked by red color of up-regulation and blue color of down-regulation.
